# Specialized high-capacity mitochondria fuel cell invasion

**DOI:** 10.1101/2025.05.02.651978

**Authors:** Isabel W. Kenny-Ganzert, Lena P. Basta, Lianzijun Wang, Qiuyi Chi, Caitlin Su, Katherine S. Morton, Joel N. Meyer, David. R Sherwood

## Abstract

Cell invasion through basement membrane (BM) is energetically intensive, and how an invading cell produces high ATP levels to power invasion is understudied. By generating 20 endogenously tagged mitochondrial proteins, we identified a specialized mitochondrial subpopulation within the *C. elegans* anchor cell (AC) that localizes to the BM breaching site and generates elevated ATP to fuel invasion. These ETC-enriched high-capacity mitochondria are compositionally unique, harboring increased protein import machinery and dense cristae enriched with ETC components. High-capacity mitochondria emerge at the time of AC specification and depend on the AC pro-invasive transcriptional program. Finally, we show that netrin signaling through a Src kinase directs microtubule polarization, which facilitates metaxin adaptor complex dependent ETC-enriched mitochondrial trafficking to the AC invasive front. Our studies reveal that an invasive cell produces high ATP by generating and localizing high-capacity mitochondria. This might be common strategy used by other cells to meet energy demanding processes.

## Introduction

Cell invasion through basement membranes (BMs), a dense sheet-like extracellular matrix (ECM) that surrounds and separates tissues^1^, is crucial for development and immune cell trafficking. Despite the formidable barrier formed by BMs, many cells acquire the specialized ability to invade and transmigrate BMs. This includes trophoblasts during embryo implantation, neural crest cells that populate diverse, mesodermal cells during gastrulation, neuronal axons that breach BM to innervate the spinal cord, and immune cells trafficking to sites of infection and injury^2–6^. Dysregulation of cell invasion also underlies many human diseases, such as rheumatoid arthritis, pre-eclampsia, and endometriosis^7–10^. Most notably, the acquisition of invasive behavior initiates cancer metastasis, which is the primary cause of cancer lethality^11^. Despite the physiological and clinical importance of BM invasion, the molecular and cellular mechanisms that drive this specialized behavior are not fully understood.

A unique feature of invasive cells is the ability to form F-actin-rich invadosomes. These membrane-associated protrusive structures harbor and secrete matrix metalloproteinases (MMPs) to breakdown and physically displace BM barriers^12–14^. To develop invasive capabilities, cells require upregulation of ribosome biogenesis, translation of numerous pro-invasive proteins, and de novo lipid synthesis to form dynamic invadosome membrane structures^15–18^. Protein translation, lipid biogenesis, membrane trafficking, and F-actin turnover, all require significant energy in the form of ATP^19–25^. Thus, another distinctive trait of invasive cells that breach BMs are mechanisms to generate high ATP levels to fuel the many molecular and cellular processes required to breakdown BM^26^.

Studies in cancer cell lines and tumor samples have revealed that metastatic cancers largely depend on mitochondrial OXPHOS to power invasion^26–29^. For example, there is an upregulation of OXPHOS and mitochondrial biogenesis genes in invasive circulating cancer cells, a decrease in breast and melanoma cancer cell invasion *in vitro* and suppression of metastasis *in vivo* after blocking OXPHOS, and a reliance on OXPHOS for proper invasion of invasive lung cancer leader cells *in vitro*^30^. Mitochondria are also localized to the leading edge of many cancer cells, such as pancreatic, prostate, ovarian, breast, glioblastoma, lung, and hepatocellular, and mitochondria enrich at the leading edge of migrating fibroblasts^31–35^. Work in several cancer cell lines and migrating fibroblasts has found that mitochondrial localization to the leading edge is dependent on kinesin-mediated trafficking along microtubules, although the signals that direct transport and many of the mechanisms mediating trafficking remain poorly understood^31,34,36^. Mitochondrial enrichment is thought to ensure high levels of localized ATP production to fuel F-actin polymerization, membrane dynamics, actomyosin contractility, and focal adhesion turnover required for cell movement and invasion^26,37^. Studies in pancreatic ductal carcinoma cells have further revealed that mitochondria fuse, increase in size and generate more ATP within protrusions invading through artificial matrices^32^. Emerging evidence indicates that mitochondria are uniquely tailored for distinct functions^38–40^, whether mitochondria that localize to the leading edge of invasive cells have other unique attributes that facilitate production of high ATP levels required for cell invasion, however, is unknown.

Anchor cell (AC) invasion through BM in *C. elegans* is a visually accessible, highly stereotyped, and genetically tractable *in vivo* model of cell invasion^41,42^. The AC is a specialized uterine cell that invades through the underlying linked gonadal and ventral epithelial BMs to initiate uterine-vulval attachment^43,44^. During AC invasion, a netrin (*C. elegans* UNC-6) cue secreted from the underlying vulval cells polarizes F-actin-rich invadosomes to the invasive front of the AC, where they dynamically assemble and disassemble and depress the BM until one breaches the BM^14,45^. At the site of BM breaching, a large invasive protrusion forms via lysosome exocytosis to expand the BM opening^14,16^. The AC’s pro-invasive transcriptional program, including the proto-oncogenes Fos and MECOM transcription factors (*C. elegans* FOS-1 and EGL-43, respectively), and BM breaching machinery, including actin regulators Arp2/3, cofilin, and matrix-degrading MMPs, are shared with invasive metastatic cancer cells^43,46,47^. The AC also harbors a robust energy acquisition and delivery system to fuel the invasive machinery, including polarized glucose transporters, glycolytic enzymes, and mitochondria that enrich at the AC’s invasive front and provide high levels of ATP to fuel invadosomes and the invasive protrusion^48,49^. The visual and molecular genetic accessibility of AC invasion, combined with a recently completed AC transcriptome^43^, provide a powerful model to establish the cellular and molecular underpinnings of mitochondria formation and composition within invasive cells.

Here, using whole-body ATP biosensors, we show that the AC harbors elevated ATP levels during its differentiation as an invasive cell and that ATP peaks and is enriched at the invasive front during BM invasion. By examining AC mitochondrial gene expression, we reveal that genes encoding components of the electron transport chain (ETC) complexes, which work to establish the electrochemical gradient that drives OXPHOS, are highly expressed and enriched in the AC. We endogenously tagged 15 components across all five ETC complexes as well as 5 mitochondrial proteins involved in import and cristae formation with mNeonGreen (mNG). These strains along with electron microscopy, dyes for mitochondrial transmembrane potential and mitochondrial lipid composition, and targeted disruptions, revealed that the AC contains a population of specialized, ETC-enriched, transport enhanced, and cristae dense, mitochondria that are preferentially localized to the invasive front and are required for the high ATP levels for BM invasion. We show these high-capacity mitochondria are specified early during AC invasive differentiation by the AC’s proto-oncogenic transcription factor network and reveal that netrin signaling guides their trafficking to the site of BM breaching through mitochondrial metaxin adaptor complexes, microtubules, and the Src family kinase, SRC-1. Together, we present the first extensive endogenously tagged mitochondrial component toolkit and discover a specialized subset of high-capacity mitochondria with increased OXPHOS capability that are preferentially trafficked to the BM breach site to fuel invasion.

## Results

### AC mitochondria produce high ATP without increasing volume, number or morphology

Anchor cell (AC) invasion is a highly stereotyped basement membrane (BM) transmigration that can be staged with the underlying 1° fated P6.p vulval precursor cell (VPC) divisions^41,42,50^. The AC is a specialized uterine cell, which is specified between the L2/L3 larval stages (early P6.p 1-cell stage, Figure 1A)^51^. During the L3 stage, the AC grows and expresses many pro-invasive proteins (P6.p 1-cell stage)^52–54^. At the P6.p early 2-cell stage many protrusive F-actin rich invadosomes rapidly form and turn over until one penetrates the BM, which triggers the exocytosis of lysosomes and focused F-actin generation at the breach site to form a single large protrusion^45,55,56^. The protrusion clears a path through the BM, allowing the AC to contact the underlying VPCs and initiate direct uterine-vulval contact (Figure 1A). Mitochondria polarize towards the AC invasive front and generate ATP required to fuel the invasion process^49,57,58^. We previously used an AC-expressed ratiometric ATP:ADP biosensor, PercevalHR^58,59^, which revealed a dramatic increase in the ATP:ADP ratio at the site of mitochondria enrichment when BM breaching and protrusion formation occur^58^. This suggested that ATP generation might be uniquely regulated in the AC.

**Figure 1.**
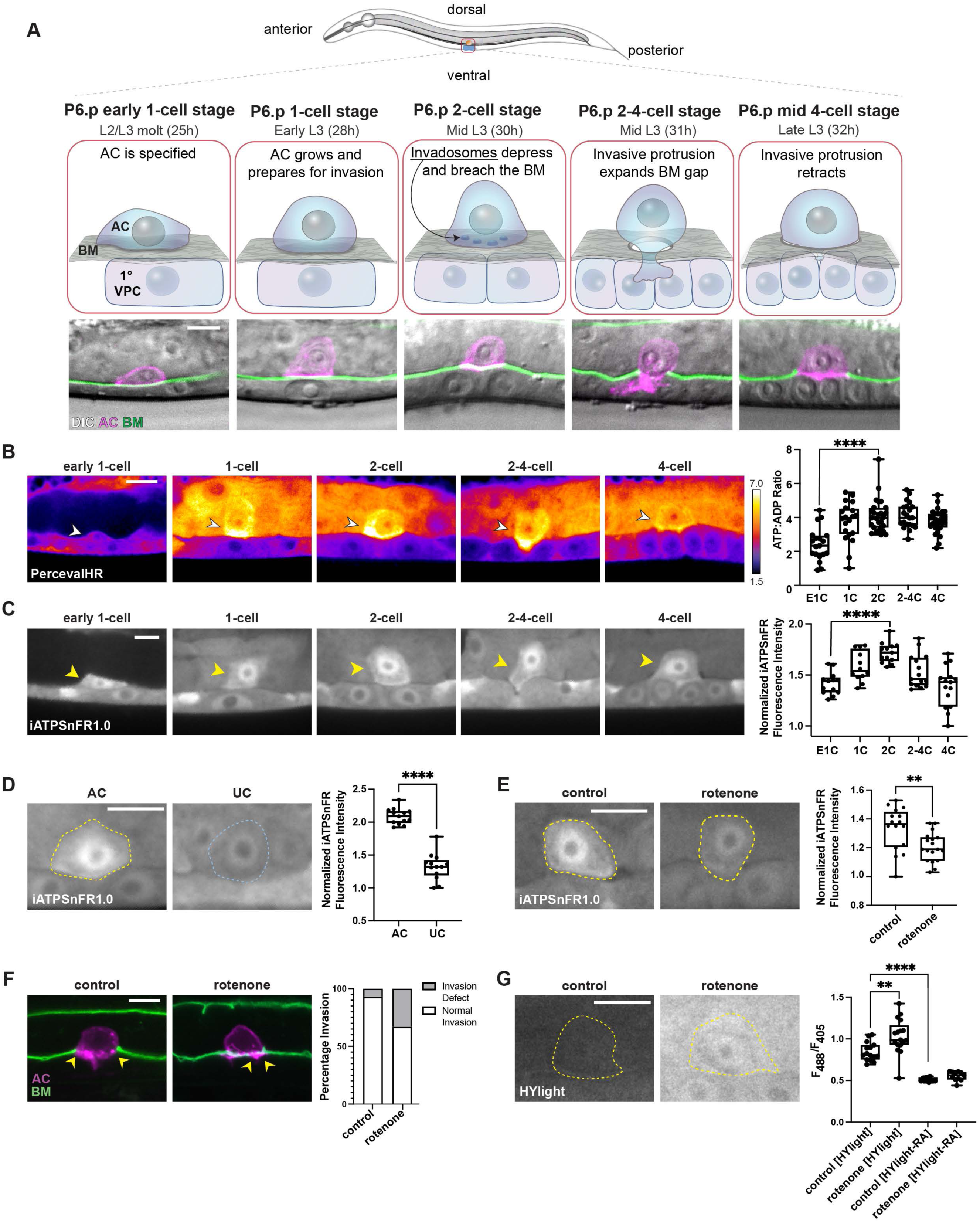
An increase in ATP accompanies AC invasion. (**A**) A timeline of anchor cell (AC) invasion through the basement membrane (BM) in L3 *C. elegans* larva (top image). Schematics (top panel) and micrographs (bottom panel; differential interference contrast (DIC, gray) microscopy with the AC (magenta, *lin-29p*::2xmKate2::PLCδPH) and the BM (green, laminin::mNG) of the lateral view of AC invasion from the P6.p early 1-cell stage of the 1° vulval precursor cells (VPCs) to the P6.p 4-cell stage. (**B**) (Left) Spectral fluorescence intensity maps of the ATP:ADP ratio as visualized by PercevalHR (*eef-1A.1p*::PercevalHR) in the AC (arrowhead) before (P6.p early 1-cell (E1C) and 1-cell stage (1C)) and during (P6.p 2-cell (2C), 2-4 cell (2-4C) and 4-cell (4C) stages) BM breaching. This and all subsequent color calibration bar represent the minimum and maximum pixel value range of the acquired data. (Right) Quantification of total ATP:ADP ratio over time, n ≥ 19 animals per time point. One-way ANOVA with Tukey’s post hoc test for multiple comparisons, **** *p* < 0.0001. (**C**) (Left) ATP levels as visualized by the ATP biosensor iATPSnFR1.0 (*eef-1A.1p*::iATPSnFR1.0) in the AC (arrowhead) before (E1C and 1C), during (2C, 2-4C), and after (4C) stages, BM breaching. Scale bar, 5 μm. (Right) Quantification of normalized iATPSnFR levels over time, n ≥ 12 animals per time point. One-way ANOVA with Tukey’s post hoc test for multiple comparisons, **** *p* < 0.0001. (**D**) (Left) ATP levels (*eef-1A.1p*::iATPSnFR1.0) at the P6.p 2-cell stage in the AC (yellow outline) and uterine cell (UC, blue outline). (Right) Quantification of normalized iATPSnFR levels in AC versus UC at the P6.p 2-cell stage, n = 13 animals. Paired two-tailed t-test, **** *p* < 0.0001. (**E**) (Left) ATP levels (*eef-1A.1p*::iATPSnFR1.0) in the AC of control and rotenone-treated animals at the P6.p 2-cell stage. (Right) Quantification of normalized iATPSnFR levels in AC of control and rotenone-treated animals, n ≥ 16 animals per condition. Unpaired two-tailed t-test, ** *p* < 0.01. (**F**) (Left) Control and rotenone-treated animals were scored for AC (magenta, *cdh-3p*::2xmKate2mCherrry::PLCδPH) invasion through the BM (green, laminin::GFP) at the P6.p early 4-cell stage. Arrowheads indicate site of BM breaching. (Right) Quantification of invasion defects of control and rotenone-treated animals, n = 42 animals per condition. (**G**) (Left) Glycolytic ratiometric sensor, HYlight (*eef-1A.1p*::HYlight), in the AC of control and rotenone-treated animals at the P6.p 2-cell stage. (Right) Quantification of HYlight and HYlight-RA (HYlight control) ratiometric levels in the AC of control and rotenone-treated animals, n ≥ 10 animals per condition. One-way ANOVA with Tukey’s post hoc test for multiple comparisons, ** *p* < 0.01, **** *p* < 0.0001. Representative ratiometric images of HYlight-RA control in Figure S1D.Scale bar, 5 μm.

To compare AC ATP metabolism with neighboring non-invasive uterine cells, we expressed the genetically encoded biosensors PercevalHR and iATPSnFR1.0 under the ubiquitous *eef-1a.1* promoter^60^. PercevalHR measures the ATP:ADP ratio, which reflects the free energy of ATP hydrolysis available for driving energy demanding processes^61^. iATPSnFR1.0 is formed from circularly permuted superfolder GFP inserted between the ATP-binding helices of the ε-subunit of a bacterial F0-F1 ATPase and is responsive to physiological levels of cytoplasmic ATP, but not ADP^62^. Interestingly, we found an increase in the ATP:ADP ratio in the AC compared to neighboring uterine cells at the P6.p 1-cell stage (n = 20/20 animals examined) several hours prior to invasion when the AC is growing and translating pro-invasive proteins (Figure 1B). This ratio peaked and was polarized toward the invasive front at the time of BM breaching (Figure 1B, 2-cell n = 27/27 and 2-4-cell stage, n = 21/21). We similarly found that the fluorescence levels of iATPSnFR1.0 were elevated at the P6.p 1-cell stage and increased ∼50% at the time of BM breaching (Figure 1C). Given the known dynamic response range of iATPSnFR1.0 fluorescence to ATP *in vitro*, this likely represents over a 3-fold increase in ATP in the AC at the time of breaching^63^. We also found a ∼40% increase in AC iATPSnFR1.0 fluorescence compared to neighboring uterine cells (Figure 1D). Importantly, we have previously shown that *eef-1a.1* drives similar expression levels in the AC and uterine cells and expression does not increase at the time of AC invasion (Figure S1A)^60^. Unlike PercevalHR, which has rapid fluorescence kinetics with exposure to ATP:ADP^61^, iATPSnFr1.0 takes up to 10 seconds to return to baseline fluorescence after ATP exposure^63^. This likely accounts for the uniform cytosolic florescence and nuclear localization of iATPSnFR1.0 in the AC^64^. iATPSnFR1.0 is a GFP based sensor and, like GFP, is sensitive to pH^63^. Importantly, we have previously shown that GFP does not show pH sensitivity in the AC^58^. Together, both PercevalHR and iATPSnFR1.0 show that the AC uniquely produces high ATP levels prior to invasion that peak during BM transmigration.

Cellular ATP can be produced via mitochondrial oxidative phosphorylation (OXPHOS) or glycolysis^65^. To establish the contribution of glycolytic metabolism to ATP production during AC invasion, we used the glycolysis ratiometric biosensor HYlight^66^. The fluorescence ratio emitted by HYlight serves as a proxy for glycolytic activity, as it measures the glycolytic metabolite fructose 1, 6-biphosphate (FBP), which is the glycolysis commitment step^66,67^. We found that glycolytic activity as measured by HYlight was the same in the AC as neighboring non-invasive uterine cells (Figure S1B), suggesting that the AC uses OXPHOS to generate higher ATP levels. To directly test this, we reduced OXPHOS by treating worms with rotenone, a potent mitochondrial toxin ^68^. Rotenone treatment caused a significant decrease in iATPSnFR1.0 fluorescence (∼40%), similar to the reduction in the ATP:ADP ratio we previously reported using PercevalHR^58^, and penetrant invasion defects (Figures 1E and 1F; Table S1). In addition, we dissipated the AC’s mitochondrial proton gradient by generating transgenic animals with AC-specific overexpression of the mitochondrial uncoupling protein UCP-4 (*lin-29p::UCP-4::SL2::mKate2::PH*)^69,70^, which also reduced the ATP:ADP AC ratio (Figure S1C). Impaired mitochondria respiration shifts neuronal metabolism towards glycolysis^67^. We thus examined HYlight after rotenone treatment and similarly found increased glycolysis (30% increase in HYlight ratio, Figure 1G, HYlight-Reduced Affinity (-RA) control Figure S1D). This might explain the residual ATP production in AC and moderate invasion defect after OXPHOS inhibition (Figures 1E and 1F). Together, these results implicate a key role for mitochondrial OXPHOS in the increased ATP production necessary for AC invasion.

Mitochondria polarize to the AC invasive front and associate with glycolytic enzymes^58^. Whether AC mitochondria have other features that contribute to high ATP levels is unknown. Increased mitochondrial biogenesis, increased mitochondrial volume, and alterations in mitochondrial morphology are implicated in enhanced ATP production^71–74^. We examined mitochondrial volume and network morphology but found no differences between the AC and neighboring uterine cells (Figure S1E and S1F). Further, we examined the localization of mitochondrial transcription factor-A (TFAM::GFP, *C. elegans* HMG-5::GFP)^75^. TFAM drives mtDNA replication during mitochondrial biogenesis, compacts mitochondrial nucleoids, and TFAM puncta mark mitochondrial nucleoids^75–78^. There was no difference, however, in the number of TFAM puncta in the AC compared to neighboring uterine cells (Figure S1G), strongly suggesting a lack of enhanced AC mitochondrial biogenesis. Together these results offer compelling evidence that the AC does not require greater mitochondria number, volume, or network morphology to drive higher OXPHOS dependent ATP levels required for invasion.

### AC mitochondria are enriched for electron transport chain components

We were next interested in determining if AC mitochondria have a distinct molecular composition that might facilitate high ATP production. Mitochondria have two phospholipid bilayers, an outer membrane (OMM) that separates the mitochondria from the cytoplasm, and a folded inner membrane (IMM) that encloses the mitochondrial matrix (Figure 2A)^79^.These mitochondrial compartments house diverse proteins that perform a variety of functions, including OXPHOS, apoptosis, and calcium regulation^79^. To determine if any mitochondrial molecular functions might be augmented in the AC, we referenced Human MitoCarta3.0, a publicly available dataset of 1136 mitochondrially-associated genes with mitochondrial pathway annotations^80^ and identified 1076 known or predicted *C. elegans* orthologs (Table S2). From the *C. elegans* mitochondrial genes, we found genes in 7 mitochondrial pathways were enriched in a recently published AC transcriptome (Figure S2A)^52^. Of those, six pathways encoded proteins involved in the electron transport chain (ETC) and OXPHOS (Figure S2A; Table S2), suggesting that increased levels of ETC components might contribute to elevated AC ATP production.

**Figure 2.**
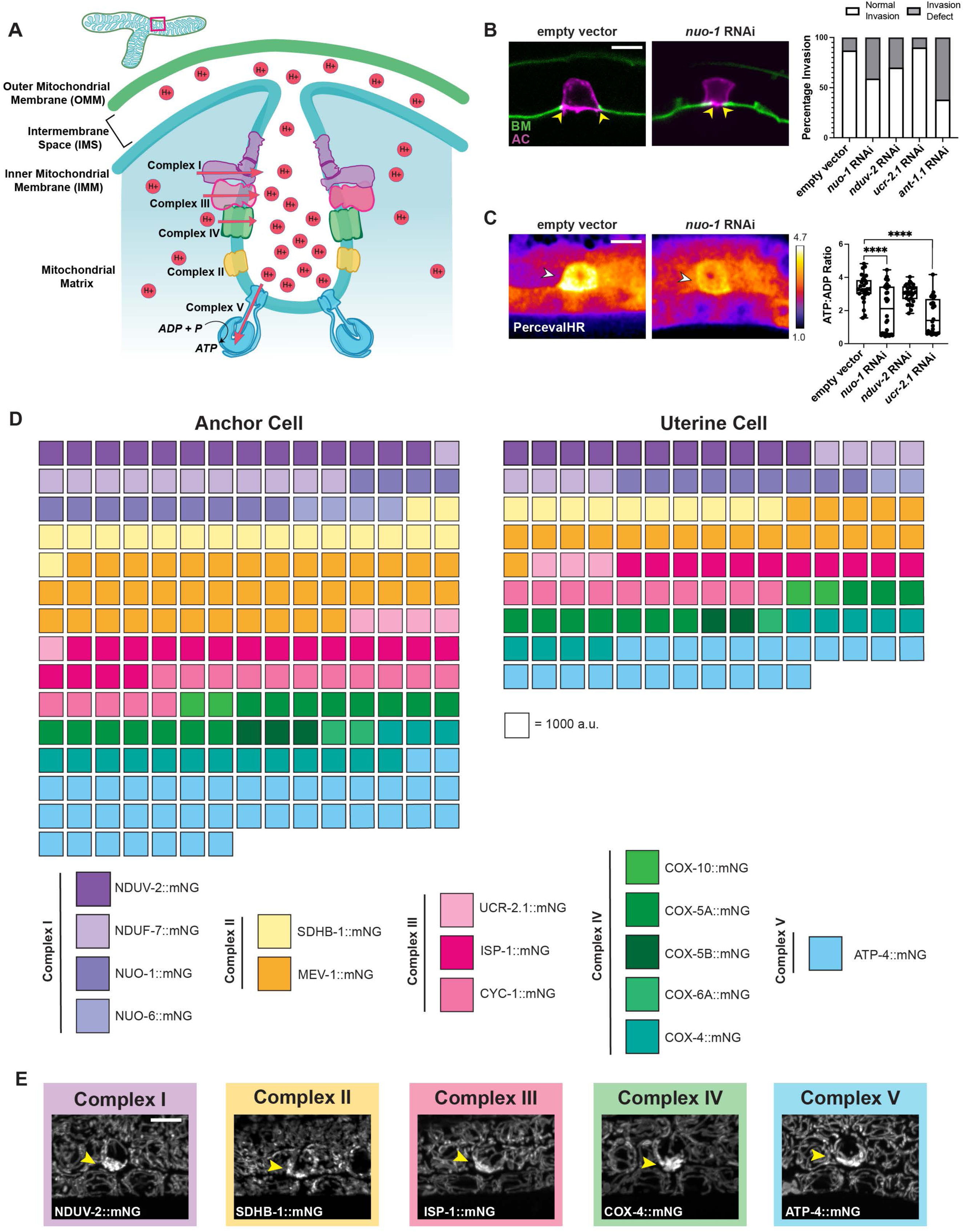
ETC components are enriched in the AC mitochondria. (**A**) Schematic of mitochondrial membrane structure and electron transport chain (ETC) localization and composition in the inner mitochondrial membrane (IMM). The ETC is comprised of five multi-protein complexes (CI-CV) that generate an electrochemical proton gradient (red H+, arrows depict proton movement) used by Complex V (ATP synthase) to phosphorylate ADP to ATP. (**B**) (Left) Empty RNAi vector control and *nuo-1* RNAi-treated animals were scored for AC (magenta, *cdh-3p*::mCherry::PLCδPH) invasion through the BM (green, laminin::dendra) at the P6.p 4-cell stage. Arrowheads indicate site of BM breaching. Note smaller breach in *nuo-1* RNAi treated animals compared to empty vector control animals. (Right) Quantification of invasion defects of empty vector control and *nuo-1*, *nduv-2*, *ucr-2.1,* and *ant-1.1* RNAi-treated animals, n = 20 animals per condition. (**C**) (Left) Spectral fluorescence intensity maps of the ATP:ADP ratio (*eef-1A.1p*::PercevalHR) in the AC (arrowhead) of empty vector control and *nuo-1* RNAi-treated animals at the P6.p 2-cell stage. (Right) Quantification of ATP:ADP ratio in the P6.p 2-cell stage AC of empty vector control and *nuo-1*, *nduv-2*, and *ucr-2.1* RNAi-treated animals, n ≥ 28 animals per condition. One-way ANOVA with Dunnett’s test for multiple comparisons, **** *p* < 0.0001. (**D**) Waffle plot of fluorescence intensity of ETC components per mitochondrion in the P6.p 2-cell stage AC versus uterine cell (UC). Each square represents 1000 a.u., n ≥ 10 animals for each ETC component, AC and UC measurements made in the same animal. (**E**) Representative images of one endogenously tagged ETC components per ETC complex in the P6.p 2-cell stage AC (arrowhead). Note the enriched expression of ETC components in the AC. Scale bar, 5 μm

The ETC is comprised of five multi-protein complexes (CI-V) localized to the IMM and generates an electrochemical proton gradient that is used to drive the rotational catalysis mechanism of ATP synthase (Figure 2A)^81^. RNAi mediated reduction of the transcriptionally enriched complexes I-IV ETC components resulted in invasion defects (Figures 2B and S2B; Table S1), consistent with an important role for the ETC in AC invasion. As components of the OXPHOS complexes might have modulatory roles in ATP production^82^, we also used RNAi to target *ant-1.1*, which encodes the dominant *C. elegans* adenosine nucleotide transporter (ANT) that transports ATP out of the mitochondria to the cytoplasm^83–85^. RNAi mediated depletion of ANT-1.1 resulted a strong invasion defect (60%) (Figures 2B and S2B, Table S1). Using the PercevalHR ATP:ADP biosensor, we further found that RNAi mediated reduction of *nuo-1, nduv-2,* and *ucr-2.1* decreased the ATP:ADP ratio in the AC (Figures 2C and S2C). As with rotenone treatment, HYlight revealed increased glycolysis after RNAi mediated reduction to *nuo-1* (Figure S2D).

To determine whether the transcriptional enrichment of ETC components correlated with protein abundance in mitochondria, we used genome-editing to insert mNeonGreen (mNG) at the C-terminus of 15 ETC proteins (Methods). This included genes encoding 4 Complex I (CI) *(nduv-2, nduf-7, nuo-1, nuo-6),* 2 Complex II (CII) (*sdhb-1, mev-1*), 3 Complex III (CIII) (*ucr-2.1, isp-1, cyc-1*), 5 Complex IV (CIV) (*cox-10, cox-5a, cox-5b, cox-6a, cox-4*) proteins, and 1 Complex V (CV) (*atp-4*) protein. We assessed the knock-in lines for viability, mitochondrial respiratory capacity, and AC invasion. Of the 15 tagged strains, 4 (*nuo-6, isp-1, cyc-1, atp-4*) were homozygous sterile. Interestingly, when maintained as heterozygotes, progeny of these lines were viable through adulthood (Table S3; see Methods), suggesting a sensitive mitochondrial germline requirement where mitochondria have high activity^86,87^. The growth rates of 7 of the 11 homozygous strains were like wild-type animals, however, 4 were slower growing (Table S3). Seahorse analysis for mitochondrial respiratory capacity of 8 homozygous viable strains revealed normal mitochondria health, with the exception of COX-4::mNG and SDHB-1::mNG, with only COX-4::mNG showing impaired basal function (Figure S2E)^88,89^. Importantly in all 15 tagged strains, homozygous animals displayed normal AC invasion, suggesting ETC function in the AC was largely unaffected (Table S1).

We next measured the fluorescence intensity of each mNG tagged ETC component per mitochondrion and compared ETC component abundance in the AC versus neighboring uterine cell mitochondria at the P6.p 2-cell stage (Figure 2D, 2E, S2F, and S2G). To control for mitochondrial density, we utilized adaptive thresholding and quantified ETC component intensity per mitochondrion (Methods). Strikingly, we found that 14 of the 15 ETC proteins were enriched (∼1.2 to 2.1-fold) in the AC mitochondria compared to non-invasive neighboring uterine cell mitochondria (Figures 2D and S2G). One of the main functions of the ETC is to generate the mitochondrial membrane potential (Δ*ψ*), which is harnessed for ATP production. Using a mitochondrial membrane potential sensitive dye, Tetramethylrhodamine Ethyl Ester (TMRE), we found that the AC has a higher membrane potential than neighboring uterine cells (Figure S2H). We conclude that the AC has ETC-enriched mitochondria that generate an increased membrane potential and produce higher ATP levels. We hereafter refer to these as high-capacity mitochondria.

### High-capacity AC mitochondria have elevated protein import machinery and dense cristae

We next investigated the cellular and molecular composition of the high-capacity AC mitochondria. The ETC localizes to inner mitochondrial membrane (IMM) folds, or cristae, which increase surface area and allow for the dense packing of ETC proteins^90,91^. To assess cristae architecture, we used transmission electron microscopy (TEM) and found that AC mitochondria contain a higher density of cristae per mitochondrion than neighboring non-invasive uterine cells (43% versus 34%, respectively, Figure S3A).

We next asked if there were molecular differences that support high ETC enriched AC mitochondria. We reasoned that high-capacity mitochondria would require greater import of nuclear encoded ETC proteins and mechanisms to support dense cristae. We thus examined the requirement of the mitochondrial protein translocases (*C. elegans* TOMM and TIMM complexes) that facilitate the import of proteins from the cytoplasm, mitochondrial contact site and cristae organizing system (MICOS) components that stabilize cristae, and the cardiolipin synthase (CRLS) enzyme that synthesizes cardiolipin, which stabilizes cristae structure (Figure 3A)^92–94^. We performed an RNAi screen targeting these genes and found that knockdown of nearly all decreased AC mitochondria enrichment of the ETC components NUO-1::mNG (CI), NDUV-1::mNG (CI) and UCR-2.1::mNG (CIII) (Figures 3B, S3B, and S3C; Table S4). Furthermore, RNAi knockdown of *tomm-20* (TOMM complex), *immt-1* (MICOS component), *crls-1* (cardiolipin synthase), disrupted AC invasion and decreased the AC ATP:ADP ratio (Figures 3C and 3D; Table S1). We created endogenous mNG knock-ins in TOMM-20, IMMT-1, and CRLS-1 and found each were enriched in the AC mitochondria compared to the neighboring uterine cells (1.3-1.6-fold, Figures 3E, 3F, and S3D). Similarly, nonyl acridine orange (NAO), which stains for cardiolipin^95^, showed ∼2-fold more cardiolipin accumulation in AC mitochondria compared with uterine mitochondria (Figure S3E). We conclude that the AC high-capacity mitochondria are built with dense cristae and an increased import system to harbor ETC enriched mitochondria that increase ATP production for invasion.

**Figure 3.**
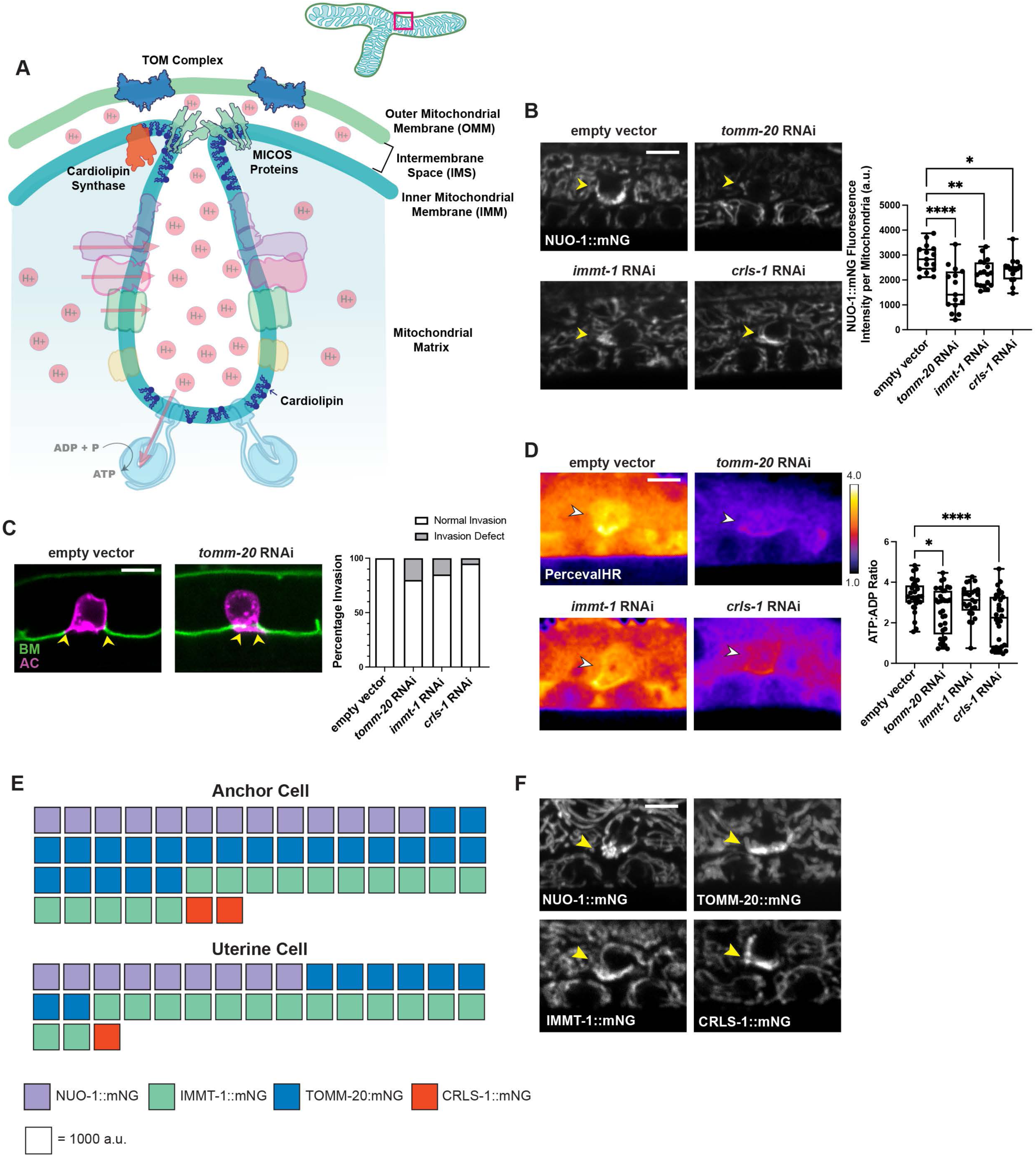
AC high-capacity mitochondria have enriched protein import machinery and dense cristae. (**A**) Schematic of cristae in the mitochondria that house the ETC complexes. Mitochondrial protein translocases (TOM complex) localize in the outer mitochondrial membrane (OMM), while MICOS proteins localize in the inner mitochondrial membrane (IMM) where they stabilize cristae formation. Cardiolipin, synthesized by cardiolipin synthase, is essential for the folding of the IMM. (**B**) (Left) NUO-1::mNG in the AC (arrowhead) at the P6.p early 2-cell stage in empty vector control and *tomm-20*, *immt-1*, and *crls-1* RNAi-treated animals. Note the loss of NUO-1::mNG enrichment in the RNAi treated animals. (Right) Quantification of fluorescence intensity NUO-1::mNG per mitochondrion in empty vector control and *tomm-20*, *immt-1*, and *crls-1* RNAi-treated animals, n ≥ 15 animals for each condition. One-way ANOVA followed by uncorrected Fisher’s LSD post hoc test, * *p* < 0.05, ** *p* < 0.01, **** *p* < 0.0001. (**C**) (Left) Empty RNAi vector control and *tomm-20* RNAi-treated animals scored for AC (magenta, *cdh-3p*::mCherry::PLCδPH) invasion through the BM (green, laminin::dendra) at the P6.p 4-cell stage. Arrowheads indicate site of BM breaching. Note smaller breach in *tomm-20* RNAi treated animals compared to control animals. Scale bar, 5 μm. (Right) Quantification of invasion defects in empty vector control and *tomm-20*, *immt-1*, and *crls-1* RNAi-treated animals, n = 20 animals per condition. (**D**) (Left) Spectral fluorescence intensity maps of the ATP:ADP ratio (*eef-1A.1p*::PercevalHR) in the AC (arrowheads) of empty vector control and *tomm-20*, *immt-1*, and *crls-1* RNAi-treated animals at the P6.p 2-cell stage. (Right) Quantification of ATP:ADP ratio in the P6.p 2-cell stage AC of vector control and *tomm-20*, *immt-1*, and *crls-1* RNAi-treated animals, n > 20 animals per condition. One-way ANOVA with Dunnett’s test for multiple comparisons, * *p* < 0.05, **** *p* < 0.0001. (**E**) Waffle plot of fluorescence intensity of IMMT-1::mNG, TOMM-20::mNG, and CRLS-1::mNG per mitochondrion in the P6.p 2-cell stage AC versus uterine cell (UC). NUO-1::mNG included as representative ETC component. Each square represents 1000 a.u., n ≥ 14 animals for each component, AC and UC measurements made in the same animal. (**F**) NUO-1::mNG, IMMT-1::mNG, TOMM-20::mNG, and CRLS-1::mNG in the AC (arrowheads) at the P6.p 2-cell stage. Scale bar, 5 μm

### High-capacity mitochondria are established early during AC invasive differentiation

We next wanted to determine if the ETC enriched mitochondria are specified by the AC pro-invasive transcriptional program^96^. EGL-43 (MECOM oncogene) is a core transcription factor of the AC transcriptional network that is crucial to specifying invasive differentiation ^43^. RNAi targeting *egl-43* dramatically reduced AC mitochondrial ETC enrichment of NUO-1::mNG and UCR-2.1::mNG, indicating high-capacity mitochondria are a component of the invasive program (Figures 4A and S4A).

**Figure 4.**
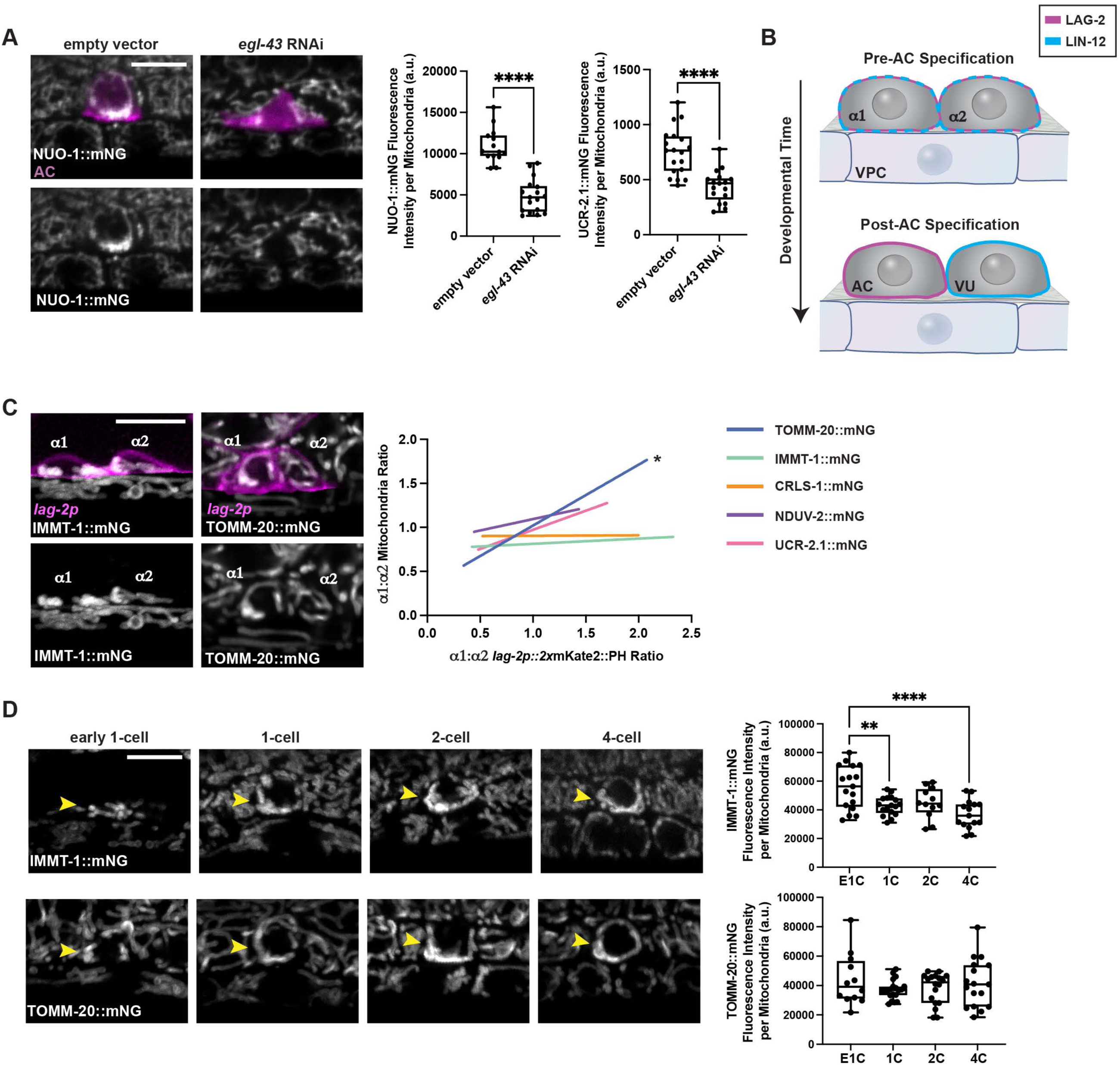
High-capacity mitochondria are established early during AC invasive differentiation. (**A**) (Left) NUO-1::mNG (grey) in P6.p early 2-cell AC (*cdh-3p*::moseinABD::mCherry, magenta) in empty vector control and *egl-43* RNAi-treated animals. (Right) Quantification of NUO-1::mNG and UCR-2.1::mNG fluorescence intensity per mitochondrion in control and *egl-43* RNAi-treated animals, n ≥ 14 animals per condition. Paired two-tailed t test, **** *p* < 0.0001. (**B**) Schematic of early AC and ventral uterine cell (VU) specification from two-proto-uterine cells (α1 and α2) via stochastic LAG-2 (Notch)/LIN-12 (Delta) signaling pre-AC specification (top) and post-AC specification (bottom). (**C**) (Left) α1 and α2 (*lag-2p*::2xmKate2::PH, magenta, top panels) and IMMT-1::mNG (grey, left panels) or TOMM-20::mNG (grey, right panels). (Right) Quantification of the *lag-2p*::2xmKate2::PH α1:α2 ratio correlated with the α1:α2 ratio of TOMM-20::mNG, IMMT-1::mNG, CRLS-1::mNG, NDUV-2::NG, or UCR-2.1::mNG, n ≥ 12 animals per strain. Simple linear regression to determine if slope is significantly non-zero, indicating a correlative relationship, * *p* < 0.05. (**D**) (Left) Developmental progression of IMMT-1::mNG (top panel) and TOMM-20::mNG (bottom panel) expression in the AC (arrowhead) from the P6.p early 1-cell (E1C) to the P6.p 4-cell (4C) stage. (Right) Quantification of IMMT-1::mNG (top graph) and TOMM-20::mNG (bottom graph) mean fluorescence intensity per AC mitochondria from the E1C to 4C stage, n ≥ 12 animals per stage for each strain. One-way ANOVA with Tukey’s post hoc test for multiple comparisons, ** *p* < 0.01, **** *p* < 0.0001. Scale bar, 5 μm.

To determine when high-capacity mitochondria are formed in the AC, we examined the timing of the molecular enrichment of ETC components, cristae components, and mitochondrial import machinery. The AC and neighboring ventral uterine cells are stochastically specified from two proto-uterine cells (α1 and α2, Figure 4B) that both have the potential to become an AC or VU (ventral uterine cell). The earliest marker of AC fate is upregulation of the LIN-12 (Notch) ligand LAG-2 in the AC (Figure 4B)^97^. We examined AC mitochondrial components that were required for ETC enrichment (TOMM-20::mNG, IMMT-1::mNG, CRLS-1::mNG), as well as ETC components (NDUV-2::mNG, and UCR-2.1::mNG) in combination with *lag-2*p::mCherry::PH (Figures 4C and S4B). We correlated the *lag-2p::*mCherry::PH signal α1:α2 ratio, with the mitochondrial component α1:α2 ratio in proto-uterine cells at the time of AC/VU specification. The non-zero slope, a measure of regression, was only significant for the relationship between TOMM-20::mNG and the *lag-2* driven signal (*i.e.* they both increased in the early specified AC at the same time). This indicates that TOMM-20 is the first component of AC mitochondria to become enriched and that this occurs at the time of AC specification. We next quantified the levels of TOMM-20::mNG, IMMT-1::mNG, CRLS-1::mNG, NDUV-2::mNG, and UCR-2.1::mNG per mitochondrion after AC specification (E1C) and leading up to (1C-2C) and after (4C) AC invasion. We found that all components at the E1C stage were elevated compared to neighboring uterine cells and that TOMM-20 remained stable across time, IMMT-1 peaked at the E1C, CRLS-1 increased over time, and both ETC proteins peaked between 1C and 2C (Figures 4D, S4C, and S4D). Transcriptional reporters of *nuo-1* and *nduv-2* (*nuo-1*p::mNG; *nduv-2*p:mNG) confirmed an increase in expression after AC specification at the E1C stage (Figure S4E). Together, these results indicate that high-capacity AC mitochondria are specified by the pro-invasive transcriptional network and begin forming during AC specification.

### Basal AC mitochondria are high-capacity and distinct from apical mitochondria

We have previously reported that mitochondria polarize at the AC invasive front ^58^. To better quantify this enrichment, we determined the volume of mitochondria in the apical and basal portions of the AC and found that ∼70% of the total mitochondria are basally localized at the 2-cell stage, while neighboring uterine cells showed equal apical/basal distribution (Figure S5A). Additionally, we found that the basal mitochondria population had higher mitochondrial membrane potential (Δ*ψ*m, TMRE staining) compared to apical mitochondria (Figure S5B). This suggested that basal AC mitochondria might be distinct from the apical mitochondria.

To determine if the composition of basal mitochondria is distinct from apical mitochondria we utilized adaptive thresholding and quantified ETC component intensity per mitochondrion of endogenously tagged ETC components (as in Figure 2) and TOMM-20, IMMT-1, and CRSL-1 (Figure 3). We found that all but one was significantly enriched (1.5-2.0-fold) in the basal mitochondria compared to apical mitochondria (Figures 5A, 5B, and S5C). We also discovered that mitochondria are initially uniformly distributed in the AC shortly after AC specification (early-1-cell stage), but segregate into two populations at the late 1-cell stage (Figure 5C).

**Figure 5.**
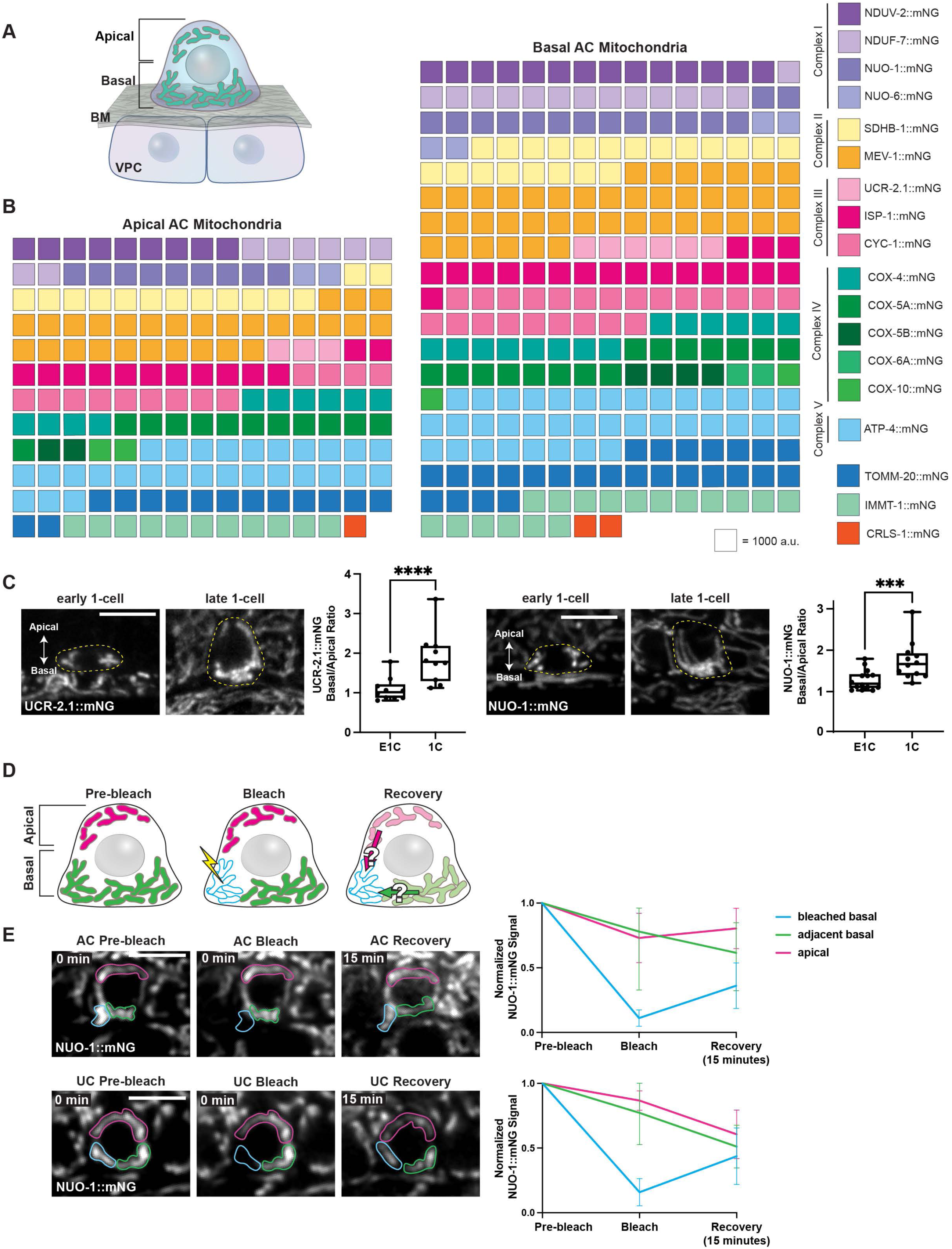
High-capacity mitochondria are basally enriched during invasion. **(A)** Schematic depicting mitochondria (teal) in both the apical and basal portions of the AC at the P6.p 2-cell stage (basement membrane, BM; vulval precursor cells, VPC). (**B**)Waffle plot of ETC components, CRLS-1, TOMM-20, and IMMT-1 fluorescence intensity per mitochondrion in the apical versus basal mitochondria in P6.p 2-cell stage AC. Each square represents 1000 a.u., n ≥ 10 animals for each protein, apical and basal measurements made in the same AC. (**C**) UCR-2.1::mNG (Left) and NUO-1::mNG (Right) expression in the P6.p early-1-cell and late-1-cell stage AC. Quantification of basal to apical UCR-2.1::mNG (Left) and NUO-1::mNG (Right) fluorescence intensity per mitochondrion in the early 1-cell and late 1-cell AC (yellow dashed outline), n ≥ 10 animals per stage. Unpaired two-tailed t-test, *** *p* < 0.001, **** *p* < 0.0001. (**D**) Schematic of FRAP experiment. (Left panel) Pre-bleach – apical mitochondria (magenta) are separate from basal mitochondria (green). (Middle panel) Bleach – 405nm laser (lightning bolt) photo-bleaches half of the basal mitochondria (blue outline). (Right panel) Recovery – determining if, and from where, the mitochondrial signal is recovered in the bleached basal region (blue outline). (**E**) (Left) NUO-1::mNG in the AC (top) and uterine cell (UC, bottom) mitochondria before, immediately after photobleaching, and 15 minutes post-photobleaching. Blue outline indicates bleached region, green outline indicates adjacent basal mitochondria, and magenta outline indicates apical mitochondria. (Right) Line graphs of normalized NUO-1::mNG fluorescence intensity pre-bleach, immediately after photobleaching, and 15 minutes after bleaching in the AC (top) and UC (bottom), n = 6 for each cell type. Scale bar, 5 μm.

To ascertain if these two populations of mitochondria intermix or remain distinct once established, we performed fluorescence recovery after photobleaching (FRAP) on mitochondria marked with endogenous NUO-1::mNG at the 2-cell stage prior to invasion. A portion of basal invasive mitochondria were photobleached, then the change in fluorescence intensity in the bleached and non-photobleached apical and neighboring basal mitochondria was measured to determine if, and from where, mitochondria moved into the bleached region (Figure 5D). We found that following photobleaching, the mitochondrial signal in the bleached basal mitochondria recovered and that the mitochondrial signal in the non-photobleached adjacent basal region decreased, while the apical signal was unchanged (Figures 5E and S5D). In contrast, similar photobleaching of neighboring non-invasive uterine cells revealed exchange between all mitochondria in the cell (Figures 5E and S5D). We conclude that the AC contains two spatially and compositionally distinct mitochondrial populations, with a specialized high-capacity basal mitochondria population with a higher membrane potential isolated from an apical population with lower ETC levels and decreased membrane potential.

### High-capacity mitochondria localize to the BM breach site and are polarized by netrin

We next wanted to examine high-capacity mitochondrial behavior during BM breaching and protrusion formation. Using endogenously-tagged type IV collagen to mark the BM (EMB-9::mRuby)^98^ with NUO-1::mNG to image high-capacity mitochondria in the AC, we performed time-lapse imaging of BM breaching and protrusion formation. In all cases we observed high-capacity mitochondria at breach site and concentrated within the emerging invasive protrusion (n = 6/6; Figures 6A, S6A, and S6B; Movie S1-3). Netrin (*C. elegans* UNC-6) secreted from the underlying 1° fated P6.p VPCs polarizes invadosomes, the invasive protrusion, and prenylation enzymes towards the invasive plasma membrane^16,45,56,58,99^. To determine if netrin signaling also polarizes high-capacity mitochondria towards the invasive front at the time of BM breaching, we examined high-capacity mitochondria (marked with NUO-1::mNG) in an *unc-6 (ev400)* null mutant. Loss of netrin disrupted high-capacity mitochondria basal enrichment and the basal enrichment of the ATP:ADP ratio (measured by PercevalHR) (Figures 6B and 6C). We conclude that netrin signaling directs high-capacity mitochondria to the invasive front and that mitochondria dynamically concentrate at the BM breach site and in the protrusion.

**Figure 6.**
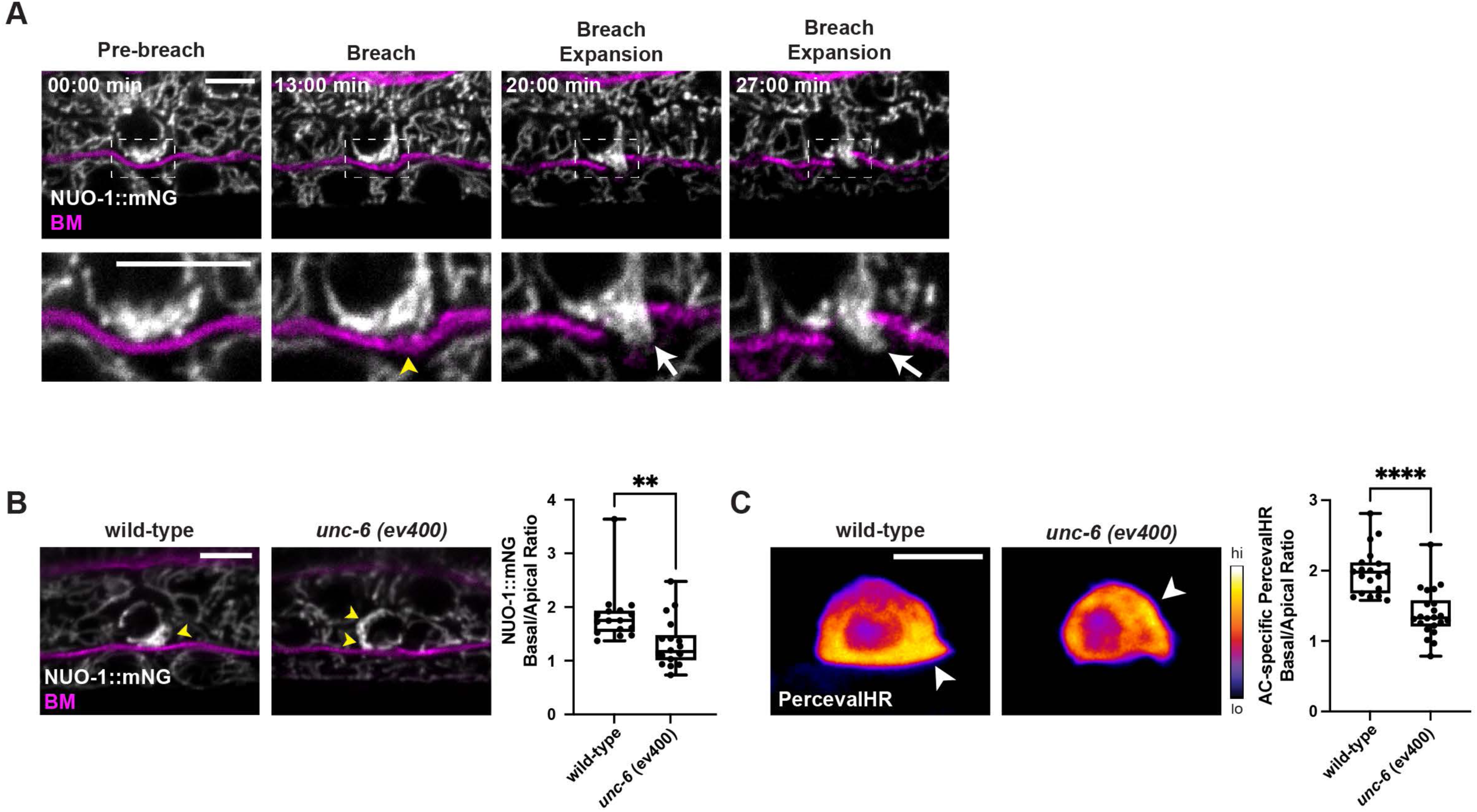
Netrin polarizes high-capacity mitochondria to the BM breach site. (**A**) Time-lapse of AC mitochondria (NUO-1::mNG, gray) prior to BM (EMB-9::mRuby2, magenta) breaching, during the initial breach, and as the breach is expanded. Animals were imaged every 1 min for 31 minutes. Bottom panels show enlarged areas corresponding to boxed regions in top panel. Yellow arrowhead indicates site of BM breaching, white arrow indicates mitochondria within the invasive protrusion. (**B**) (Left) NUO-1::mNG with BM (EMB-9::mRuby2, magenta) in wild-type and netrin mutant (*unc-6 (ev400*)) animals at the P6.p 2-cell stage. Yellow arrowheads indicate site of NUO-1::mNG enrichment. (Right) Quantification of basal to apical NUO-1::mNG fluorescence intensity per mitochondrion in wild-type and *unc-6* (*ev400*) mutant animals, n ≥ 14. Unpaired t-test, ** *p* < 0.01. (**C**) (Left) Spectral fluorescence intensity maps of the AC-specific ATP:ADP ratio (*lin-29p*::PercevalHR) of wild-type and netrin mutant (*unc-6 (ev400*)) animals at the P6.p early 2-cell stage. Arrowheads indicate site of ATP:ADP enrichment. (Right) Quantification of ATP:ADP ratio in basal versus apical regions of wild-type and *unc-6* (*ev400*) mutant P6.p 2-cell AC, n ≥ 19. Unpaired t-test, **** *p* < 0.0001. Scale bar, 5 μm.

### Netrin/Src polarizes microtubules to direct high-capacity mitochondria via metaxin adaptors to the invasive front

Microtubules transport mitochondria to areas of high ATP demand in neurons and migrating cells in culture^31,32,35,100,101^. Netrin regulates microtubule dynamics, stability, and elongation to attract and direct growing neuronal axons^102^. We have previously shown that microtubules are enriched at the invasive front^58^. To determine if netrin regulates microtubules in the AC, we visualized AC microtubules using an endogenously tagged microtubule-binding domain of ensconsin (*lin-29p*::EMTB::GFP) in wild-type animals and *unc-6* mutants and found that microtubule basal enrichment was greatly reduced (Figure 7A). Microtubules have two functionally distinct ends – the plus end with β-tubulin exposed which polymerizes/depolymerizes faster than the minus end with α-tubulin exposed^103^. To determine if microtubules are polarized in the AC, we examined the endogenously-tagged plus-end protein (EBP-2::GFP) and minus-end protein (GFP::GIP-1)^104^, and found that microtubules in the AC are oriented with the plus-end at the basal end of the cell and the minus-end at the apical (Figures 7B). These observations suggest that trafficking along polarized microtubules could direct high-capacity mitochondria to the invasive front.

**Figure 7.**
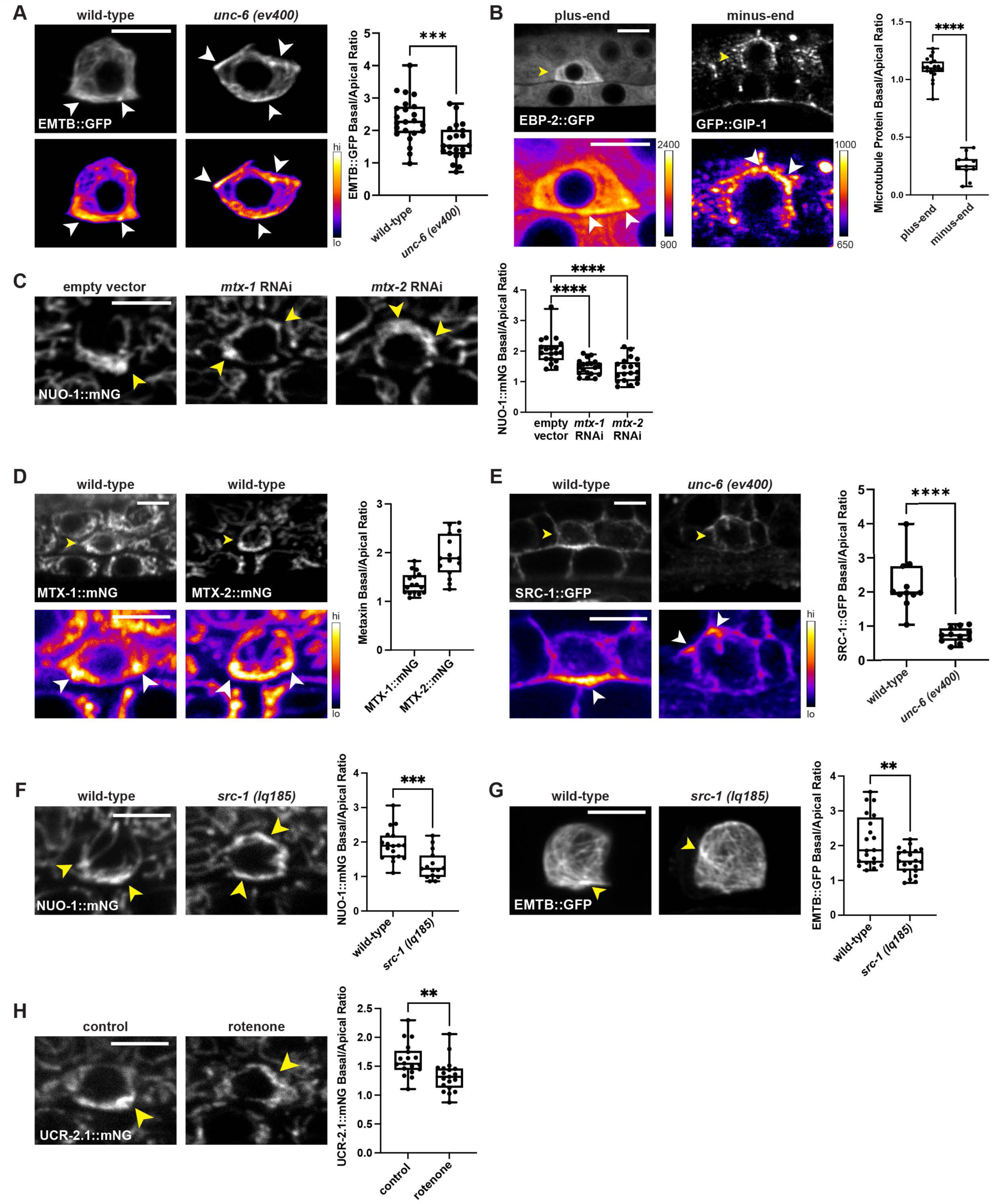
Netrin-mediated microtubule polarization directs mitochondria to the invasive front. (**A**) (Top) AC-specific microtubule marker (*lin-29p*::EMTB::GFP, top) in wild-type and netrin mutant (*unc-6 (ev400*)) animals at the P6.p 2-cell stage. (Bottom) Spectral fluorescence intensity maps applied to emphasize microtubule enriched areas (arrowheads). (Right) Quantification of basal to apical microtubule fluorescence intensity in wild-type versus *unc-6 (ev400)* mutant animals, n ≥ 20 animals. Unpaired two-tailed t-test, *** *p* <0.001. (**B**) (Top) Endogenously tagged plus-end (EBP-2::GFP) and minus-end (GFP::GIP-1) microtubule-associated proteins in the AC (yellow arrowhead) at the P6.p 2-cell stage. (Bottom) Spectral fluorescence intensity maps applied to emphasize microtubule-associated protein enriched areas (white arrowheads). (Right) Quantification of basal to apical microtubule plus-end and minus-end protein fluorescence intensity, n ≥ 11. Unpaired two-tailed t-test, **** *p* < 0.0001. (**C**) (Left) NUO-1::mNG in the AC of empty vector control and *mtx-1* and *mtx-2* RNAi-treated animals at the P6.p early 2-cell stage. Yellow arrowhead indicates NUO-1::mNG enriched mitochondria. (Right) Quantification of basal to apical NUO-1::mNG fluorescence intensity per mitochondrion in empty vector control and *mtx-1* and *mtx-2* RNAi-treated animals, n = 19 animals. One-way ANOVA with Dunnett’s multiple comparison test, **** *p* < 0.0001. (**D**) (Top) Endogenously tagged MTX-1::mNG and MTX-2::mNG in P6.p 2-cell stage ACs (yellow arrowhead). (Bottom) Spectral fluorescence intensity maps of MTX-1::mNG and MTX-2::mNG to emphasize metaxin enriched areas (white arrowheads). (Right) Quantification of basal to apical MTX-1::mNG and MTX-2::mNG fluorescence intensity per mitochondrion, n = 15 animals for each strain. (**E**) (Top) SRC-1::GFP in the AC (yellow arrowhead) in wild-type and netrin mutant (*unc-6 (ev400*)) animals at the P6.p 2-cell stage. (Bottom) Spectral fluorescence intensity maps to emphasize SRC-1::GFP enriched areas (white arrowheads). (Right) Quantification of basal to apical SRC-1::GFP fluorescence intensity in wild-type versus *unc-6 (ev400)* mutant animals, n = 11 animals. Unpaired two-tailed t-test, **** *p* < 0.0001. (**F**) (Left) NUO-1::mNG in P6.p 2-cell ACs in wild-type and *src-1 (lq185)* mutant animals. Yellow arrowhead indicates NUO-1::mNG enriched mitochondria. (Right) Quantification of basal to apical NUO-1::mNG fluorescence intensity per mitochondrion in wild-type versus *src-1 (lq185)* mutant animals, n ≥ 15 animals. Unpaired two-tailed t-test, *** *p* < 0.001. (**G**) (Left) AC-specific microtubule marker (*lin-29p*::EMTB::GFP) in P6.p 2-cell ACs in wild-type and *src-1 (lq185)* mutant animals. Yellow arrowhead indicates microtubule enriched areas. (Right) Quantification of basal to apical microtubule fluorescence intensity in wild-type versus *src-1 (lq185)* mutant animals, n = 20 animals. Unpaired two-tailed t-test, ** *p* < 0.01. (**H**) (Left) UCR-2.1::mNG in P6.p 2-cell ACs in control and rotenone-treated animals. Yellow arrowhead indicates UCR-2.1::mNG enriched mitochondria. (Right) Quantification of basal to apical UCR-2.1::mNG fluorescence intensity per mitochondrion in control and rotenone-treated animals, n = 18 animals. Unpaired two-tailed t-test, ** *p* < 0.01. Scale bar, 5 μm.

Microtubules serve as tracks for motor proteins and their adaptors to transport cargo. Notably, genes encoding mitochondrial trafficking proteins were among the upregulated MitoCarta pathways in the AC transcriptome (Table S2). Of these mitochondrial associated proteins, we were particularly interested in MTX-1, a metaxin, which functions as an OMM mitochondrial adaptor for plus ended directed mitochondrial trafficking^105,106^. The two *C. elegans* metaxin homologs, MTX-1 and MTX-2, have functionally distinct roles in trafficking. MTX-2 works with the known microtubule adaptor, MIRO-1, as part of the core adaptor complex for both microtubule plus-end and minus-end transport, whereas MTX-1 only serves as an adaptor for plus-end transport^106^. RNAi-mediated loss of *mtx-1* and *mtx-2*, significantly reduced the basal polarity of high-capacity mitochondria (Figures 7C and S7A). Furthermore, we endogenously-tagged both *mtx-1* and *mtx-2* with mNG and found MTX-1::mNG and MTX-2::mNG are basally enriched in the AC and thus enriched in the high-capacity mitochondria (Figure 7D). We also found that metaxin localization was dependent on netrin signaling and loss of netrin (*unc-6* mutant) resulted in a loss of basal metaxin enrichment (Figure S7B). We conclude that high-capacity mitochondrial enrichment at the site of invasion is netrin dependent and that basally localized mitochondria are trafficked on plus-end microtubules using the metaxin adaptor complex.

We next wanted to determine what links the netrin (*unc-6*) directional cue to polarized microtubule transport. Evidence suggests that Src family kinases couple netrin signaling to microtubule dynamics through phosphorylation of β-tubulin, which regulates microtubule dynamics^102,107^. *C. elegans* harbor two *src* genes*, src-1* and *src-2*^108^ and *src-1* is expressed at high levels in the AC transcriptome^52^. We examined endogenously-tagged SRC-1::GFP and found basal AC enrichment in an *unc-6* dependent manner (Figure 7E). Further, the basal enrichment of both mitochondria and microtubules was reduced in a *src-1(lq185)* mutant background (Figures 7F and 7G), indicating that SRC-1 polarizes microtubules and mitochondria to the basal invasive front.

AMPK, an energy sensor and metabolic regulator, has been shown in ovarian cancer cells to direct mitochondria to the leading edge of lamellipodia^31^. RNAi knockdown of four AMPK components (*aak-1*, *aak-2*, *aakb-1*, and *aakb-2)*, however, did not disrupt mitochondrial localization (Figure S4D). Interestingly, previous studies in neurons and several cancer cell lines have shown that loss of ATP production reduces mitochondrial trafficking along microtubules^31,35,109–111^. suggesting that ATP output could be used as a mechanism that preferentially traffics high-capacity AC mitochondria along polarized microtubules to the invasive front. Consistent with this notion, both pharmacological (rotenone treatment) and genetic (*nuo-1* RNAi) perturbation of ETC function resulted in a loss in the basal enrichment of high-capacity mitochondria (Figures 7H and S7C). Taken together, our results suggest a model where netrin (UNC-6) signaling through SRC-1 polarizes microtubules towards the invasive front. This facilitates trafficking of high-capacity mitochondria harboring the metaxin adaptor complex to supply high levels of ATP for BM breaching (Figure S7E).

## Discussion

There is an emerging notion that mitochondria specialize to meet the functional needs of different cells and tissues^38,112^. Tissue level biochemical studies provided the first evidence of specialized mitochondria and showed distinct mitochondrial proteomes in the mouse brain, liver, heart, and kidney^113^. This was followed by findings that identified distinct mitochondria within different cell types, including mitochondria in oocytes lacking assembled ETC complex I to mitigate damaging ROS production^114^, mitochondria within fibroblasts that lack ATP synthase and are specialized to produce proline and ornithine^115^, and mitochondria with decreased matrix proteins and mitochondrial DNA and an upregulated Ca^2+^ uniporter that regulate filipodia length in osteosarcoma cells^40^.

Here we show that the *C. elegans* AC has specialized mitochondria that produce high levels of ATP to fuel invasion through BM. Transcriptomic analysis revealed broad enrichment of ETC components and examination of 15 endogenous mNeonGreen (mNG) tagged ETC components identified a distinct subpopulation of mitochondria localized to the invasive front of the AC. In these high-capacity mitochondria, increased TOM complex transport proteins and dense cristae facilitate import and housing of higher levels of ETC components that produce an elevated mitochondrial membrane potential driving high ATP production at the site of BM breaching. Notably, these mitochondria form shortly after AC specification, and this coincides with increased ATP levels a full ∼5 hours before AC invasion. The elevated ATP prior to invasion likely supports energy demanding cellular processes necessary for invasion, such as increased translation of pro-invasive proteins and de novo lipid synthesis^16–18,52^. Further, we found mitochondrially produced ATP peaked at the time of BM breaching when energy consuming invasive protrusion dynamics occur^58^. Notably, there was also heterogeneity in the upregulation of different ETC components within the high-capacity mitochondria, both within ETC complexes and between complexes. The difference in stoichiometry between components may indicate further specialization of ETC function and highlights an unknown complexity of the ETC.

The AC high-capacity mitochondria appear to be built for heightened ATP generation, which modeling experiments suggest is the rate limiting step for supplying cytosolic ATP^116^. ETC-enriched mitochondria might be a common strategy for cells to support energetically demanding processes. For example, proteomic studies of mitochondria in the heart and synapse, which both are energy intensive, have shown mitochondria contain higher levels of some ETC components^117,118^. Further, imaging studies at the synapse, have revealed these mitochondria contain dense cristae and elevated cytochrome C (complex III)^119^. ^120^. Adipocyte mitochondria associated with lipid droplets also have dense cristae, elevated levels of the ETC complex IV component Cox4 and elevated respiratory capacity, which is thought to support ATP-dependent triacylglyceride synthesis^121^. In addition, high levels of ETC might be a common feature of invasive cells, as studies in aggressive ovarian, breast, pancreatic, and lymphoma cancers have revealed upregulation of various ETC components and complexes^122–129^.

Mitochondria traffic on microtubules to areas of high energy demand, including to the leading edge of invasive and migrating cells and to neuronal synapses and dendrites^31,36,37,100,110,130,131^. However, the cues and mechanisms that direct trafficking are largely unknown. Our studies indicate that netrin signaling is a key component of high-capacity mitochondria trafficking to the site of BM invasion through polarization of microtubules to the invasive front. It’s been proposed that netrin signaling polarizes microtubules to redirect microtubules towards sources of netrin by balancing both the promotion of microtubule dynamics and stabilization^132,133^. Evidence suggests that netrin might stabilize microtubules through Src family kinase mediated phosphorylation of tubulin^133^. Consistent with Src acting downstream of netrin to polarize microtubules, we discovered that SRC-1 is upregulated in the AC and localized to the invasive front in a netrin dependent manner. Further, loss of *src-1* disrupted microtubule and high-capacity mitochondria polarization.

We also show that high-capacity mitochondrial trafficking is dependent on metaxin adaptors. Metaxins were first identified as mitochondrial proteins in the sorting and assembly machinery (SAM) complex and have been separately found to be essential in TNF-induced apoptosis^134,135^. Evidence from *C. elegans*, *Drosophila*, and human neurons suggests that metaxin-2 is also a core component of a mitochondrial transport adaptor complex and in combination with either metaxin-1 or TRAK-1 directs plus end or minus end mitochondrial movements, respectively^105,106^. Our work extends the function of metaxins beyond neurons to invasive and migratory cells. In the AC we found that the plus-ends of microtubules are oriented towards the site of BM invasion. Consistent with studies in neurons, loss of *mtx-1* (plus-end adaptor) and *mtx-2* (core adaptor) dramatically perturbed basal (plus-end directed) enrichment of high-capacity mitochondria. Further, we previously found that loss of TRAK-1 alone does not alter basal mitochondria enrichment^58^, suggesting a minor or absent function for minus end directed movement in high-capacity mitochondria positioning.

An open question is how high-capacity mitochondria preferentially localize to the invasive front over mitochondria that have reduced ETC components and ATP production and remain apically in the AC. AMPK, an energy sensor, has been shown to direct mitochondria to leading edge of lamellopodia in ovarian cancer cells, although the molecular targets of this kinase for directing trafficking are not known^31^. However, we did not observe defects in high-capacity mitochondrial trafficking after knockdown of four AMPK components, suggesting AMPK does not regulate mitochondrial trafficking in the AC. Reduction of the ETC component NUO-1 and inhibition of OXPHOS with rotenone, which both reduced ATP production, however, perturbed high-capacity mitochondrial enrichment to the invasive front. Thus, when high-capacity mitochondria no longer produce elevated ATP, they appear not to be basally trafficked. As ATP diffusion is limited in cells^136^ and trafficking on microtubules is energy intensive^31,137^, one possible explanation is a competitive self-organizing system, where high-capacity mitochondria with MTX-1 (plus ended directed adaptor) and increased ATP output are more competitive in moving along microtubules, and thus preferentially traffic to the ends of microtubules at the invasive front. Consistent with this model, disruption of ATP production in mitochondria limits their trafficking in cancer cells and neurons^35,109–111^. Molecular competition of shared limited resources is known to govern a number of biological processes, such as preference for mRNA competition to be translated by ribosomes, sigma factor competition to bind with RNA polymerase to dictate gene transcription, and competition between RNA binding proteins that drives localized phase separation in *C. elegans* embryos^137–139^. As metaxins are also enriched on high-capacity mitochondria, this could bestow the high-capacity mitochondria an additional competitive advantage in loading on to microtubules.

Mitochondria are responsible for numerous cellular processes beyond ATP production, including fatty acid synthesis, stress response, cell-cell signaling, and calcium homeostasis^140,141^. Thus, specialized mitochondria might be a common, but hidden feature of cells. A key impediment in assessing mitochondrial diversity is the challenge of detecting differences in individual mitochondrial molecular composition within and between cells. By endogenously editing 20 genes with mNG tags whose encoded proteins are components of mitochondria—15 ETC, two cristae, two metaxins and a TOMM complex component—our studies have greatly expanded the toolkit of live cell reagents to examine mitochondrial diversity. Our endogenously tagged strains have revealed ETC-enriched high-capacity mitochondria in the AC and led to insights into single cell mitochondrial heterogeneity, specialization, specification, and trafficking. As the lineages and differentiation programs for *C. elegans* cells are known in detail^142^, this emerging toolkit can be used and expanded to investigate mitochondrial specialization in other cell types, how mitochondria are regulated in cell division, apoptosis, and oogenesis, how mitochondria respond to environmental changes, and how mitochondria are altered during aging. Our Mitocarta-based curation indicate approximately 1000 known proteins associated with *C. elegans* mitochondria. Ultimately, it should be possible to endogenously tag most of these mitochondrial components, which offers to reveal the breadth of mitochondrial diversity and lead to new insights into unknown mitochondrial functions.

### Limitations of this study

Our studies indicate that the AC has ETC enriched mitochondria that generate a higher membrane potential and ATP to drive cell invasion. The ETC is composed of over 100 components that are subdivided into five complexes, each with unique functions^143^. Testing the function of the entire upregulated ETC complex was not possible. Instead, we relied on a diverse approach and targeted individual ETC components, chemically inhibit complex I, reduce mitochondrial membrane potential, and decrease ATP release from mitochondria. While our data demonstrate that the ETC is upregulated to generate more ATP, we cannot rule out upregulation enhances or alters other functions of the ETC, such as ROS signaling, metabolite transport, Ca^2+^ regulation, and regeneration of electron carriers^140,141^. A related limitation is that we could weaken ETC upregulation but not eliminate it. Thus, while we were able to target the enriched ETC by knocking down individual components, reduce the membrane potential with the uncoupler UCP-4, and target the ATP transporter *ant-1.1*, we could not completely reduce the upregulated ETC and thus cannot assess its full contribution to breaching the BM and cell invasion.

## Methods

### Experimental Model Details

*Caenorhabditis elegans* (*C. elegans*) were maintained at 20°C on nematode growth media (NGM) plates and fed *Escherichia coli* strain OP50. In text and figures, wild-type refers to N2 Bristol strain *C. elegans,* and standard *C. elegans* nomenclature is used to convey genotypes of strains. Promoter-driven transgenes are denoted by the italicized gene name followed by “p”, “::” signifies an adjoining protein, and italicized mutant alleles are enclosed within parentheses following the gene name. As previously described, anchor cell (AC) invasion was scored in L3 hermaphrodites and staged in reference to the number of vulval precursor cells (VPCs) (Figure 1)(Sherwood and Sternberg, 2006). Synchronization of L1 animals was performed using standard hypochlorite treatment for all RNAi and rotenone experiments^144^. All endogenously tagged strains in this study were generated by injecting into the gonads of wild-type *C. elegans*. Endogenous strains that were homozygous infertile were either maintained by picking animals with the fluorescent signal of interest or crossed with a balancer strain (detailed in Key Resources Table). Measurements of fluorescence intensity in all strains were done in homozygous animals (See Table S3 for details on the health of endogenous tagged strains). All animals were well-fed (3+ generations without starvation) prior to any experimentation or imaging to control for nutrient-dependent phenotypes. All genotypes of all strains were verified by genotyping PCR and/or plate-level phenotype, such as “unc” – uncoordinated animals that are unable to crawl. All strains used in this study are listed in the key resources table.

### Construction of endogenously-tagged strains

CRISPR-Cas9 mediated genome editing with a self-excising hygromycin selection cassette (SEC) was used to generate endogenously tagged strains^145^. In brief, optimal location to insert the fluorescent protein was identified based on the tertiary and quaternary structure of the protein and any protein cleavage sites were accounted for (such as mitochondrial localization sequences that get cleaved). To generate the SEC repair plasmids, N2 gDNA was used as a template to amplify ∼2KB homology arms upstream and downstream of the PAM site. To ensure the Cas9 doesn’t subsequently cut the PAM site after the initial insertion cut, the mNeonGreen (mNG) was inserted in between the guide sequence and PAM or silent mutations to the guide sequence were made in the homology arms. The homology arms were inserted into the vector mNG-C1^SEC^bothlink SEC repair template^146^ and confirmed correct assembly by colony PCR and sequencing. PureLink™ HiPure Plasmid Miniprep (Invitrogen #K210002) was used to isolate high purity sgRNA plasmid DNA and SEC repair plasmid DNA for injection. https://crispor.gi.ucsc.edu/ was used to identify short guide sequences where the fluorescent protein should be inserted and PAM sites where the Cas9 protein cleaves. The short guide RNA (sgRNA) plasmid was generated by cutting the Cas9 guide plasmid, pDD122 (Addgene, #47550), with Nhel and EcoRV, then using HiFI assembly (New England Biolabs, #E2621L) to insert the respective sgRNA sequences into the plasmid. For each strain the germline of 10-30 young adult N2 hermaphrodites were injected with a mixture containing 100ng/μL of the SEC repair plasmid DNA, 50ng/μL of the sgRNA plasmid DNA, and 2.5 ng/ul of co-injection markers (pCFJ90 *myo-*2p::mCherry, pCFJ104 *myo-3*p::mCherry). After 3-4 days at 20°C, F1 progeny of singled-out injected animals were treated with 500μl of 2mg/ml Hygromycin B (Sigma-Aldrich #H3274). Candidate knockin animals exhibited rolling (due to *sqt-1(e1350)* within the SEC), survived Hygromycin B treatment, and lacked the red fluorescent co-injection markers. F2 rolling (*sqt-1(e1350)*+) animals were singled, confirmed homozygous knockin through roller phenotypes and presence of consistent fluorescence signals. To excise the SEC, we heat shocked about 6 L3/L4 homozygous rollers at 34°C in a water bath for 4 hours. After 3-4 days, adult non-rolling animals were singled to check for homozygous excision by confirming loss of rolling in all progeny. Successful genome editing was verified by visualizing fluorescence and PCR genotyping. See Table S5 for list of oligonucleotides used in strain generation and genotyping and See Key Resources Table for all strains.

### Construction of promoter-driven transgenic strains

Promoter-driven transgenic strains were generated through the Mos single copy insertion (MosSCI) on Chromosome I or Chromosome II or extrachromosomal array integration. Promoters and fusion proteins were either amplified from N2 gDNA (*nuo-1* promoter, *nduv-2* promoter, *ucp-4*, *tomm-20*), another plasmid (*eef-1A.1* promoter, *lin-29* promoter, *GFP, mNeonGreen, mKate2, PercevalHR, HYlight, HYlight-RA*), or a *C. elegans* codon-optimized gene block (*iATPSnFR1.0*). The Cas9 guide plasmid, pCFJ352, which is targeted near the ttTi4348 Mos insertion site, was used to generate transgenes via the Mos single copy insertion (MosSCI) on chromosome I. pCFJ352 was a gift from Erik Jorgensen (Addgene plasmid # 30539; http://n2t.net/addgene:30539; RRID:Addgene_30539). The Cas9 guide plasmid, pDD122, which is targeted near the ttTi5605 Mos insertion site, was used to generate transgenes via the Mos single copy insertion (MosSCI) on chromosome II^147^.The SEC repair plasmids for MosSCI chromosome I (pAP088) and MosSCI chromosome II (pAP087) were cut with restriction enzymes, NheI and NotI, then the respective promoter and fusion protein were cloned into the cut SEC repair plasmid backbone using NEBuilder HiFi DNA Assembly master mix (NEB, #E2621L). Injection, selection, excision, and genotyping were performed as described above. The following strains were generated by MosSCI: The biosensors (iATPSnFR1.0 and HYlight) and their controls (cytosolicGFP and HYlight-RA) were driven by the *eef-1A.1* ubiquitous promoter. Ubiquitous mitochondrially-localized mKate2 to visualize all mitochondria independent of endogenous fluorescence intensity (driven by *rpl-28* ubiquitous promoter) and AC-specific overexpression of UCP-4::SL2::mKate2 (driven by the AC promoter *lin-29*). See Key Resources Table for all strains.

To generate strains using extrachromosomal array integration, *unc-119(ed4)* hermaphrodites were injected with promoter-driven fusion protein plasmids, 50 ng/ml *unc-119+* rescue DNA, 50ng/ml pBsSK, and 25 ng/ml EcoRI cut salmon sperm DNA. Stable extrachromosomal lines were established and then integrated using gamma radiation^148^. To reduce the possibility of background mutations resulting from radiation, integrated lines were backcrossed three times with N2 animals. See Key Resources Table for all strains.

### Feeding RNAi

RNA interference (RNAi) was performed by feeding animals *Escherichia coli* strain HT115 containing the L4440 RNAi vector targeting genes of interest or the empty L4440 vector. RNAi clones were sourced from either the Vidal or Ahringer feeding RNAi libraries, then streaked out on LB plates containing tetracycline and ampicillin. A single RNAi colony was inoculated in LB with 100 μL/mL Ampicillin (Sigma-Aldrich #A0166) and grown overnight (12-16 hours) at 37°C in an incubator with a shaker. To induce double-stranded RNA expression, 1 μL/mL isopropyl β-d-1-thiogalactopyranoside (IPGT, Sigma-Aldrich #I6758) was added to RNAi cultures and placed back in the 37°C shaker for 1 h. After initial dsRNA induction, RNAi cultures were seeded on NGM plates containing topically applied 1mM IPTG and 100-mg/ml ampicillin and allowed to dry at room temperature overnight for further induction.

All RNAi clones were sequenced to confirm correct target gene. Synchronized L1 animals were plated on RNAi NGM plates, fed for 39 hours for imaging P6.p 2-cell stage or 42 hours to score invasion at the P6.p late 4-cell stage. For every RNAi experiment, a negative control (empty L4440 RNAi vector) and either knockdown efficiency measurement (Table S6) or positive control (*fos-1a* RNAi that causes a penetrant invasion defect) was used to ascertain RNAi activity.

### Rotenone Treatment

Rotenone (EMD Millipore #557368) was reconstituted in DMSO to make a 20mM stock solution and stored at −20°C. Synchronized L1 animals were plated on NGM plates seeded with *E. coli* OP50 and allowed to grow for 34 hours at 20°. Early L3 larvae were washed off from NGM plates with M9 solution and collected into 1.5 mL canonical tubes. 20mM rotenone (final working concentration 40 μM) were added to the collecting tube and placed on a rocker or rotator (to avoid hypoxia) for 2-hours at room temperature. The same volume of DMSO in M9 was used on control animals for the equivalent amount of time. Before imaging, animals were recovered from the incubation onto fresh OP50 NGM plates.

### Seahorse Extracellular Flux Analyzer-based measurements of mitochondrial respiration

All experiments were performed with bleach synchronized L3 *C. elegans* from wild-type N2 lines and 8 endogenously tagged ETC component lines (*nuo-1::mNG*, *nduv-2::mNG*, *sdhb-1::mNG*, *mev-1::mNG*, *ucr-2.1::mNG*, *cox-4::mNG*, *cox-6A::mNG*, *cox-10::mNG*). L3 *C. elegans* were washed from OP50 plates and transferred into each well of a 24-well Seahorse plate. The Seahorse Extracellular Flux Bioanalyzer was used to quantify oxygen consumption rate (OCR) as described in previous studies^149^. The number of individual animals per well were counted using confocal imaging of the plate to normalize the OCR measurements per individual worm. For each of the 9 lines, 5-10 L3 animals per strain were used and four technical replicates were done for each strain.

### Assessment of AC invasion

Assessment of anchor cell (AC) invasion was performed as previously described^42^. In brief, synchronized L3 animals at the VPC P6.p late 4-cell stage were mounted onto 5% agar with 1% sodium azide to anesthetize the worms and imaged on a Ziess upright compound microscope equipped with 488 and 561 nm filters and a Nomarski prism for differential interference contrast (DIC). The assessment of AC invasion was based on two indicators: loss of the DIC phase dense BM line and absence of the fluorescence BM marker underneath the AC. Complete removal of the BM resulting in a gap of the width of anchor cell nucleus is scored as normal invasion. Incomplete or blocked invasion is scored if the size of the BM gap is less than the width of the AC nucleus or fully intact respectively. The sensitized strain referred to in Table S1 is null for matrix metalloproteases (MMPs) *zmp-1, 3, 4, 5, 6*. Loss of MMPs delays invasion and requires more ATP to breach the BM^49^.

### Mitochondria import and cristae screen

Candidate genes for the mitochondrial import and cristae screen were compiled by finding *C.elegans* orthologs of mammalian genes annotated as having a role in mitochondrial protein import and cristae formation^80^ for which RNAi clones were available and sequenced correctly. RNAi plates were made according to the RNAi protocol described above. Screening was conducted with the strain (NK2657) with co-labeled endogenous NUO-1::mNG (mNeonGreen) and AC-specific plasma membrane marker (mCherry). Animals were synchronized by hypochlorite treatment, plated on RNAi or L4440 control, and grown until the VPC P6.p 2-cell stage. Screening was performed on a Ziess upright compound microscope equipped with 488 and 561 nm filters. We scored animals for the loss of NUO-1::mNG fluorescence enrichment in the AC compared to the uterine cell within the same animal. Systematic effects of the RNAi were evaluated by comparing signal of the promoter driven AC-specific mCherry to the control. RNAi treatment that led to a decrease in the mCherry signal would suggest a nonspecific effect. All genes that decreased the enrichment of NUO-1::mNG in the AC subsequent to knockdown were reported as hit (Table S) and each hit was repeated three times to validate the effect.

### AC-specific UCP-4 Overexpression

The plasmid containing the AC-specific overexpression of UCP-4 also contained a membrane-localized mKate2 (mKate2::PLC∂P). When imaging the effects of UCP-4 on the ATP:ADP ratio, we confirmed expression of the mKate2.

### Microscopy and image acquisition

Confocal images were acquired on a Ziess Axioimager microscope equipped with either a Yokogawa CSU-10 or CSU-W1 spinning disk confocal controlled by Micromanager Software vv1.4.23 or v2.0.1 using a Zeiss 100x Plan-Apochromat 1.4NA oil immersion objective and with either a Hamamatsu ORCA-Fusion sCMOS camera, Hamamatsu ORCA-Quest qCMOS camera, or ImageEM EMCCD camera. Animals were mounted on 5% noble agar pads and anesthetized with either 0.01M sodium azide (Sigma-Aldrich #S2002) or 5mM Levamisole (Millipore Sigma #L9756). Time-lapse imaging of AC invasion was performed as previously described on a Ziess Axioimager microscope with a Yokogawa CSU-W1 spinning disk and Hamamatsu ORCA-Fusion sCMOS camera ^150^. Fluorescence recovery after photobleaching (FRAP) was done on a Ziess Axioimager microscope equipped with an iLas targeted laser system from BioVision using an Omicron Lux 60mW 405nm laser, Yokogawa CSU-W1 spinning disk confocal controlled by Metamorph imaging software, a Zeiss 100x Plan-Apochromat 1.4NA oil immersion objective and Hamamatsu ORCA-Fusion sCMOS camera.

All quantitative mitochondrial protein measurements of endogenously tagged strains (Figures 2D and 3E) were imaged using identical acquisition settings (488nm laser at 0.6 laser power, 150ms exposure, 128eGain, relative z-stack from −3.5µmto 3.5um, z-step size 0.37um) along with a control strain to confirm reproducibility across multiple imaging sessions. PercevalHR was imaged as previously described^57^. Briefly, optimal acquisition settings for both the ATP (488nm excitation and 525nm emission) and ADP (405nm excitation and 525nm emission) channels were determined to be 488nm laser at 2.5 laser power with 500ms exposure and 405nm laser at 3.0 laser power with 1000ms exposure. To prevent photobleaching from 405nm laser from effecting 488nm excitation, multidimensional acquisition parameters were set to acquire in order of channel then z-slice, meaning all z-slices of the ATP channel were acquired before the ADP channel. HYlight was imaged as previously described^67^. Using the published physiological ratio for the HYlight-RA control (FBP-bound/FBP-unbound ratio of ∼0.4)^67^, appropriate acquisition settings were determined as 488nm laser at 2.5 laser power with 800ms exposure and 405nm laser at 3.0 laser power with 800ms exposure. For all experiments measuring glycolysis, both HYlight and HYlight-RA were imaged and with the same acquisition settings. Due to the photosensitivity of the sensor HYlight(-RA) one z-plane was imaged per animal. Quantitative imaging of iATPSnFr1.0, cytosolic GFP, and ETC promoter driven fluorescent expression, was carefully acquired with standardized acquisition settings to assure intensiometric measurements were reproducible across developmental time and experimental condition.

### Staining with mitochondrial dyes

Tetramethylrhodamine, Ethyl Ester, Perchlorate (TMRE, ThermoFisher Scientific #T669) was reconstituted in DMSO and diluted to 1μM in M9 buffer. A mixed population of animals were added to fresh OP50 NGM plates with 400μL of 1μM TMRE. Animals were stained and allowed to continue to grow overnight in the dark at room temperature before imaging. Nonyl Acridine Orange (NAO, ThermoFisher Scientific #A1372) was reconstituted in M9 then diluted to 5μM in M9 buffer. As with the TMRE protocol, 100μL of 5μM NAO was added to OP50 NGM plates with a mixed population of animals. The NAO treated animals were grown overnight at room temperature before imaging. To account for variability in the uptake of dyes, AC mitochondrial measurements of TMRE and NAO were internally controlled by either measuring neighboring uterine cells or measuring both apical and basal mitochondria within the same cell.

### Assembling *C. elegans* MitoCarta

MitoCarta is a compendium of human and mouse mitochondrial genes coding for proteins in the 149 annotated mitochondrial pathways^80^. The current version of MitoCarta3.0 includes 1136 human mitochondrial genes. Using OrthoList2.0, a comparative orthologs prediction tool between *C. elegans* and human genome^151^, we identified 824 *C. elegans* orthologs to the human MitoCarta3.0 genes. Out of the 312 human genes that Ortholist2.0 did not annotate, orthologs for an additional 131 genes were retrieved manually from the Alliance of Genome Resources^152^. In total, 84% of the Human MitoCarta genes had orthologous genes in *C. elegans* (955 out of 1136 genes) and 16% of the Human genes did not have known *C. elegans* orthology (181 out of 1136). Ortholist2.0 and Alliance of Genome Resources cross reference 6 andd 9 different gene ortholog databases, respectively, accounting for the discrepancies in ortholog annotations. The number of databases confirming orthologs for each gene is reported in Table S2. The number of databases confirming the orthology is reported in the Table to represent the source, either from Alliance or from Ortholist2.0. A total of 1094 (including duplicates of any *C. elegans* orthologs) *C. elegans* mitochondrial genes were compiled (Table S2).

### Transcriptome analysis of the *C.elegans* MitoCarta

The *C. elegans* MitoCarta (CMC) genes were cross-referenced with a recently generated AC transcriptome^52^ to determine the transcriptional enrichment of each gene in the AC compared to the whole animal (as measured by Log2-fold change). Gene Set Enrichment Analysis (GSEA) was conducted based on the data set of the transcriptional enrichment (log2-fold change) of each CMC genes and the gene set of 149 mitochondria pathways^153–155^. The GSEA algorithm filters the gene set size and removes all mitochondrial pathways where the number of genes is less than 2^155^. A list of 145 gene sets/mitochondrial pathways were analyzed and sorted based on the normalized enrichment score (NES). The significance of the enrichment was interpreted with a false discovery rate (FDR) Q-value that is less than 0.05. Plotting for the GSEA (Figure S2) was conducted in R studio (version 4.4.0).

## Quantification and Statistical Analysis

### Quantitative Mitochondrial Protein Measurements

All acquired images of endogenously-tagged mitochondrial proteins were processed using ImageJ/Fiji. Mitochondria signal within the cell of interest (either AC or UC) was analyzed using 3-slice sum projections (0.2 μm z-step size) confocal z-stacks (with background subtraction, 6.7 pixel rolling ball radius). A mask for each mitochondria in the cell of interest was generated using the adaptive thresholding plugin^156^. This mask was used as a region of interest and the mean fluorescence intensity was measured to determine the “total” fluorescence per mitochondria for all mitochondria in the cell. The cell was then manually subdivided into apical and basal regions and the mean fluorescence intensity per mitochondria was measured in each region. Polarity was then calculated using the following ratio:

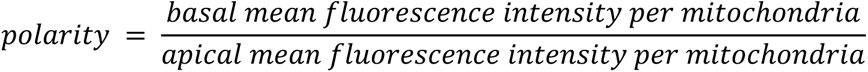

### Mitochondrial volume quantification

Using a mitochondrially localized mKate (*rpl-28p::tomm-20::mKate*), a confocal z-stack was acquired through the entire uterus (51 slices, 0.2 µm step size) and the images were background subtracted using 6.7 pixel rolling ball radius. To generate an accurate projection of all mitochondria in each analyzed cell (both AC and uterine cell), regions of interest around the boundary of each cell were manually drawn and any signal outside of this ROI was removed for each z-slice. The new compiled stack for each cell of interest was subsequently analyzed using Imaris Version 9.9.1 to generate a 3D isosurface rendering. 3D isosurface renderings relied on adaptive thresholding as described above and were used to determine the volume of the mitochondria within each cell.

### Mitochondrial Morphology Assessment

Z-stacks of all the mitochondria within the cell of interest were generated as described above. Using the ImageJ plugin “Mitochondrial Analyzer^156^”, each z-stack was thresholded and mitochondrial sphericity and branch number were measured.

### HMG-5 puncta

Using animals with endogenously tagged HMG-5 (*hmg-5::GFP*)^157^, a confocal z-stack was collected through the entire uterus (51 slices, 0.20 µm step size). 4 central slices of the AC or UC were sum z-projected and the number of puncta were manually counted.

### Transmission Electron Microscopy cristae analysis

Using transmission electron microscopy (TEM) images from a depositary of images from previously published work^44^, boundaries around individual mitochondria were manually drawn and annotated as either AC or UC. Those images were then assigned random numbers to blind the analysis. For each blinded image the freehand selection tool was used to circle and measure both the outer perimeter of the mitochondria (OMM) and the inner membrane (IMM) folds/cristae. The total area of the IMM and OMM were individually measured. The ratio of IMM to OMM were obtained by dividing the total area of IMM by the total area of the OMM.

### Analysis of ratiometric biosensors

Ratiometric biosensors PercevalHR and HYlight(-RA) were used to determine the ATD:ADP ratio and FBP-bound:FBP-unbound ratio, respectively. As described above (See Microscopy and Image Acquisition), the imaging data for these biosensors were acquired as two-channel images. To transform the two-channel images into ratiometric images, the “Imaging Calculator > Divide” function in Fiji was used to divide the signal of the 488nm-excitation channel (ATP for PercevalHR and FBP-bound for HYlight(-RA)) by the 405nm-excitation channel (ADP for PercevalHR and FBP-unbound for HYlight(-RA)). The divided ratiometric images were used for quantification. Representative images for figures were displayed as the spectral intensity map using the “Fire” look-up table in Fiji (LUT).

### Spectral representation of fluorescence intensity

Spectral intensity maps were used to emphasize differences in fluorescent intensity. Using the “Fire” look-up table (LUT) in Fiji, the spectral intensity map and corresponding calibration bar were applied to representative images shown in figures. In instances where all representative images within the panel were constrained to the same minimum and maximum pixel intensity, exact values were displayed at the bottom and top of the calibration bar, respectively. Calibration bars were annotated with “hi” and “lo” (represent high and low intensity pixel values) for representative images that were displayed with different minimum and maximum pixel intensity either due to variation in the image acquisition settings or normalization of ATP:ADP ratios.

### Statistical Analysis

For all experiments, the sample size (*n*) of animals, cells, or mitochondria measured was listed in the figure legend or table, along with the statistical tests used and the p-values. All statistical analyses and graph generation were done in GraphPad Prism (Version 10). Normality of each experimental data set was evaluated using the Shapiro-Wilk test. Comparisons across multiple time-points were performed using a one-way ANOVA with Tukey’s post hoc test for multiple comparisons. Comparisons of fluorescence intensity between two cells in the same animal (AC vs US)or between two mitochondrial populations within the same cell (apical vs basal) were conducted using paired t-test or mixed-effects ANOVA with Geisser-Greenhouse correction and uncorrected Fisher’s LSD to allow for comparison between designated pairs. Comparisons of fluorescence intensity between a control and treatment were made using a two-tailed unpaired t-test with Welch’s correction if the variance was unequal between the two groups. Comparisons of fluorescence intensity between a single control and two or more treatments were performed by one-way ANOVA with Dunnett’s test for multiple comparisons. When assessing AC invasion defects based on invasion scoring, a Fisher’s exact 2×2 test was used to compare the treatment verses the control. When determining if there was a statistically significant relationship between the lag2 ratio and the mitochondrial component fluorescence intensity per mitochondria ratio between the α1 and α2 cells, measure of non-zero slope was used to interpret the results of the simple linear regression analysis.

## Acknowledgements

We wish to thank Jake Leyhr for assistance with MitoCarta data curation and presentation, Siddharthan Balachandar Thendral for movie processing, Meghan Morrissey for generation of the TEM image repository, and Delaney Graham for help with RNAi screening. Additionally, we would like to thank Laura Kelley for helpful discussions throughout the project. We gratefully thank the Pablo Lara-Gonzalez lab for the SRC-1::GFP strain and the Colón-Ramos lab for supplying the HYlight and HYlightRA plasmids. The Caenorhabditis Genetics Center, supported by the National Institutes of Health Office of Research Infrastructure Programs (P40 OD01044), provided some of the *C. elegans* strains. I.W.K., L.P.B., Q.C., L.W., and D.R.S. were supported by R35GM118049 and a Department of Biology Hargitt fellowship. J.M., and K.M. were supported by P42ES010356 and T32ES021432.

## Author contributions

Conceptualization: I.W.K. and D.R.S.; Data curation: I.W.K.; Formal analysis: I.W.K., L.P.B., K.M., L.W.; Funding acquisition: D.R.S.; Investigation: I.W.K., L.P.B, L.W., K.M., C.S.; Methodology: I.W.K., L.P.B., Q.C., K.M.; Project administration: I.W.K., D.R.S.; Resources: J.N.M. and D.R.S.; Supervision: D.R.S.; Visualization: I.W.K., L.P.B., L.W., K.M.; Writing – original draft: I.W.K., L.P.B., D.R.S.; Writing – review & editing: I.W.K., L.P.B., D.R.S.

---

**Figure S1.**
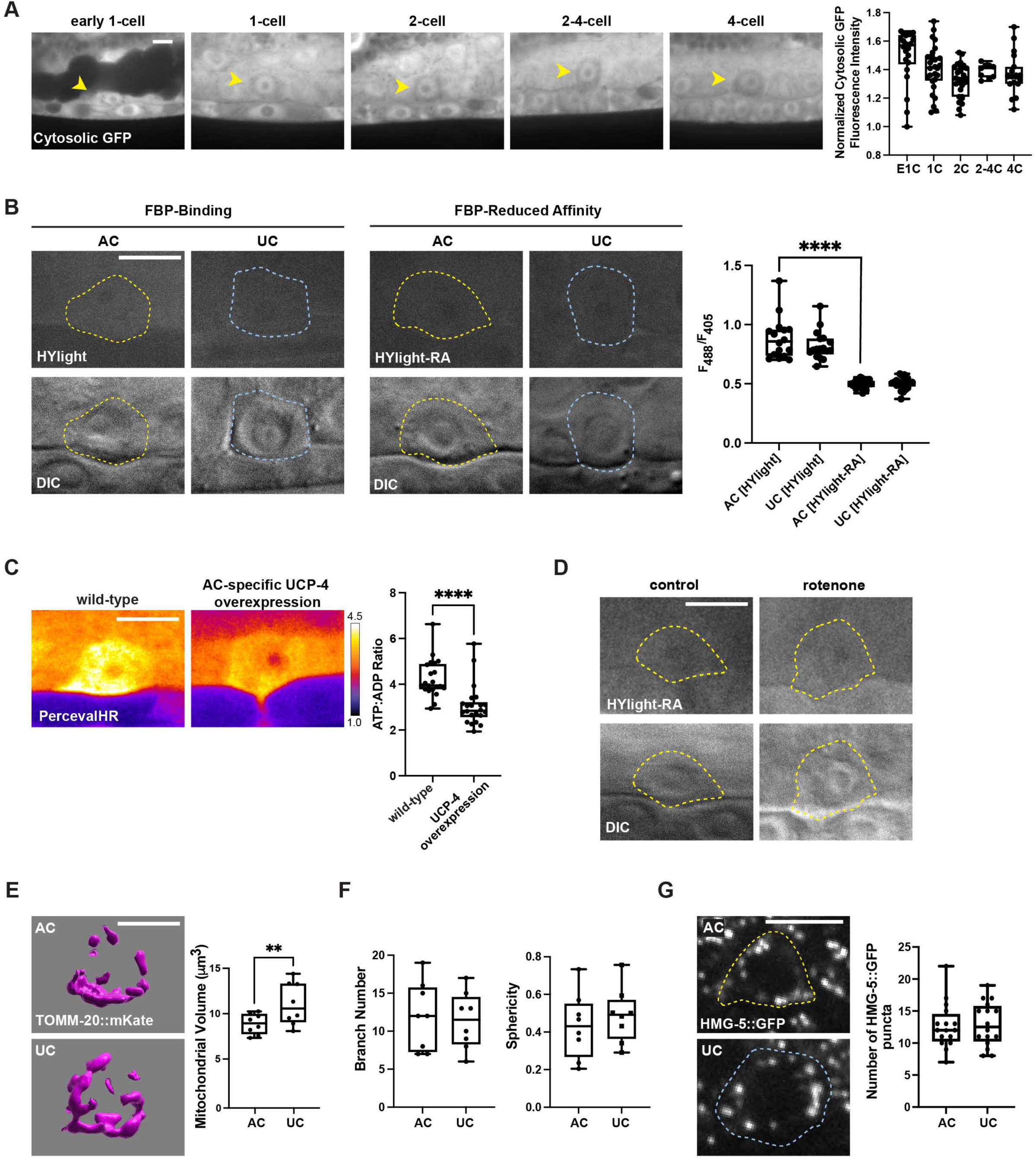
ATP increase in the AC is not a result of glycolysis or mitochondrial abundance, related to Figure 1. (**A**) (Left) Promoter-driven cytosolic GFP levels as visualized by *eef-1A.1p*::GFP as a control for *eef-1A.1p*::iATPSnFR1.0. Representative images of GFP in the AC (arrowheads) before (P6.p early 1-cell (E1C) and 1-cell (1C) stage) and during (P6.p 2-cell (2C), 2-4 cell (2-4C)) and after (4-cell (4C) stages) BM breaching. (Right) Quantification of normalized cytosolic GFP levels over developmental time, n ≥ 10 animals per time point. One-way ANOVA with Tukey’s post hoc test for multiple comparisons. Note there is no significant differences in GFP levels between adjacent stages. (**B**) (Left) HYlight ratiometric glycolytic sensor (*eef-1A.1p*::HYlight) and (Middle) HYlight-RA control (*eef-1A.1p*::HYlight-RA) and differential interference contrast (DIC) images of P6.p 2-cell AC (yellow outline) and uterine cell (UC, blue outline). The HYlight-RA control has reduced affinity for FBP and does not respond to changes in FBP concentration. (Right) Quantification of HYlight and HYlight-RA ratiometric levels in the P6.p 2-cell AC and UC, n ≥ 16 animals per condition. One-way ANOVA with Tukey’s post hoc test for multiple comparisons, **** *p* < 0.0001. (**C**) (Left) Spectral fluorescence intensity maps of the ATP:ADP ratio (*eef-1A.1p*::PercevalHR) in wild-type and UCP-4 overexpression worms (*lin-29p::UCP-4::SL2::mKate2::PH*).(Right) Quantification of ATP:ADP ratio in wildtype and UCP-4 overexpression P6.p 2-cell AC, n = 10 animals. Unpaired two-tailed t-test **** *p* < 0.0001. (**D**) HYlight-RA (*eef-1A.1p*::HYlight-RA) glycolytic sensor control, in the AC of control and rotenone-treated animals that correspond to the data presented in Figure 1G. Note no difference in HYlight-RA in control and rotenone-treated animals. (**E-F**) Mitochondrial morphology characterization in AC versus UC at the P6.p 2-cell stage. (**E**) (Left) Isosurface renderings of mitochondrial volume (visualized by *rpl-28p*::TOMM-20::mKate2, magenta) using Imaris. (Right) Mitochondrial volume quantification determined from isosurface renderings, n = 8 animals. Paired two-tailed t-test, ** *p* < 0.01. (**F**) Quantification of mitochondrial branch number and sphericity, n = 8 animals. Paired two-tailed t-test. Note no significant difference in branch number or sphericity between AC and UC. (**G**) (Left) Mitochondrial DNA transcription factor, HMG-5, as visualized by HMG-5::GFP in AC (yellow outline) and UC (blue outline) at the P6.p 2-cell stage. (Right) Quantification of the number of HMG-5::GFP puncta in the AC and neighboring UC, n = 16 animals. Paired two-tailed t-test. Note no significant difference in HMG-5 puncta between AC and UC. Scale bar, 5 μm.

**Figure S2.**
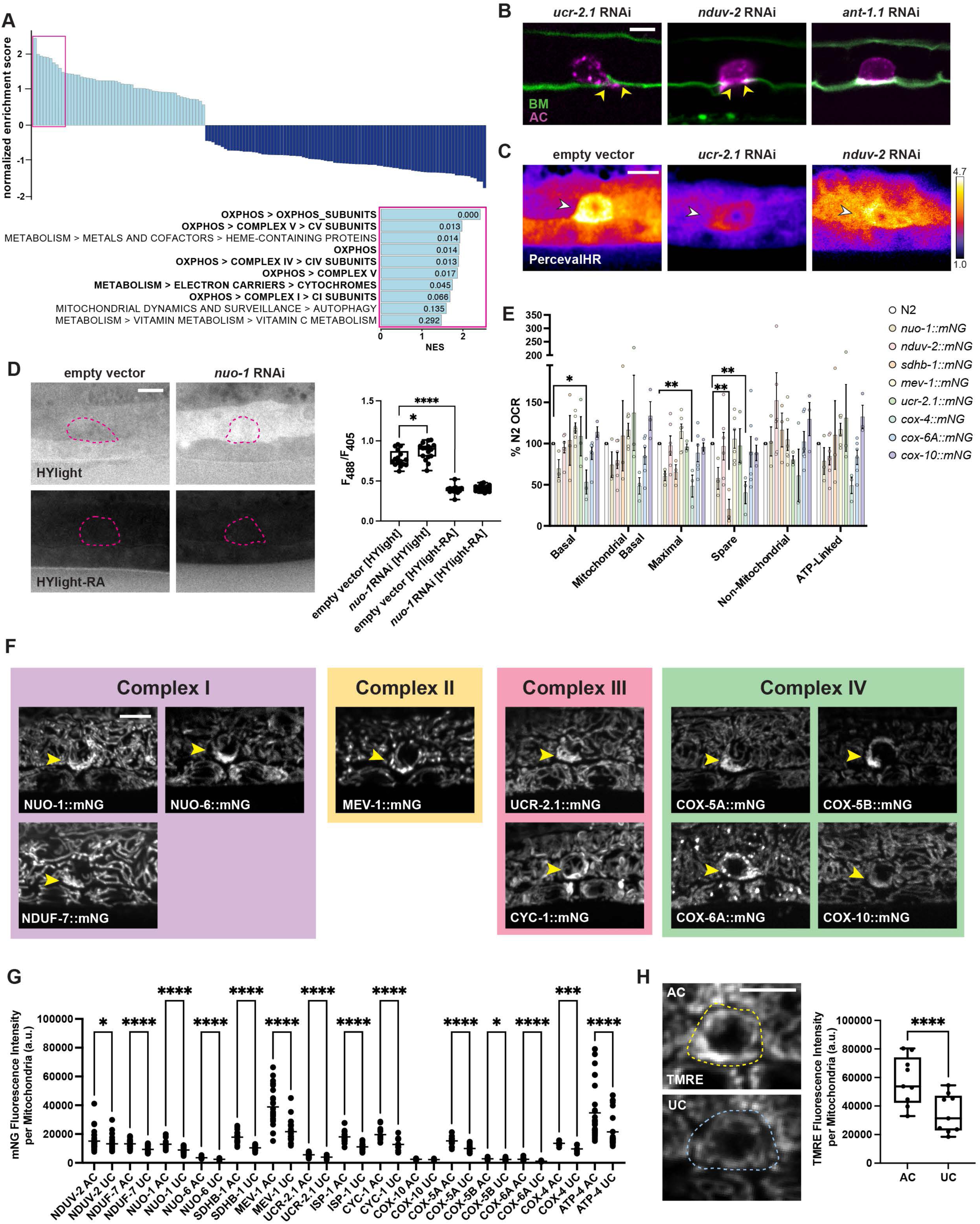
AC mitochondria are enriched in ETC components as quantified via transcriptomics and visualized by endogenous tags, related to Figure 2. (**A**) MitoCarta^78^ Pathways gene-set enrichment analysis performed on transcriptome^50^ comparing the AC and the whole body. Two-sided bar plot visualizes the normalized enrichment score (NES) for MitoCarta pathways. Pink box indicates the top ten enriched pathways in the AC. Note 7 of the 10 pathways (bolded) encode proteins involved in the ETC and OXPHOS. (**B**) AC invasion defects in *ucr-2.1*, *nduv-2* and *ant-1.1* RNAi-treated animals. Arrowheads indicate sites of incomplete BM (green, laminin::dendra) breaching by the AC (magenta, *cdh-3p*::mCherry::PLCδPH) at the P6.p 4-cell stage, corresponding to the data quantified in Figure 2B. Note example of blocked invasion from *ant-1.1* RNAi treatment. (**C**) Spectral fluorescence intensity map of the ATP:ADP ratio (*eef-1A.1p*::PercevalHR) in the AC (arrowhead) of empty vector control, *ucr-2.1,* and *nduv-2* RNAi-treated animals at the P6.p early 2-cell stage, corresponding to the data quantified in Figure 2C. (**D**) (Left) HYlight (*eef-1A.1p*::HYlight) and HYlight-RA control (*eef-1A.1p*::HYlight-RA) in the AC (pink outline) in empty vector control and *nuo-1* RNAi-treated animals. (Right) Quantification of HYlight and HYlight-RA ratiometric levels in the AC of empty vector control and *nuo-1* RNAi-treated animals, n = 16 animals per condition. One-way ANOVA with Tukey’s post hoc test for multiple comparisons, **** *p* ≤ 0.0001, * *p* < 0.05. (**E**) Seahorse measurements of mitochondrial oxygen consumption rate (OCR) parameters for 8 endogenously tagged ETC. Tagged strains compared to wild-type (N2) OCR. One-way ANOVA with Dunnett’s multiple comparisons test, * *p* < 0.05, ** *p* < 0.01. (**F**) Endogenously tagged ETC components within each complex in the P6.p 2-cell stage AC (arrowhead). (**G**) Quantification of fluorescence intensity of mNG tagged ETC components per mitochondrion in the AC versus the uterine cell (UC), n ≥ 10 animals for each tagged component, AC and UC measurements made in the same animal. Mixed-effects ANOVA with Geisser-Greenhouse correction, followed by uncorrected Fisher’s LSD post hoc test, * *p* < 0.05, *** *p* < 0.001, **** *p* < 0.0001. (**H**) (Left) TMRE staining for mitochondrial membrane potential in the AC (top, yellow outline) versus UC (bottom, blue outline) of P6.p 2-cell stage wild-type (N2) animals. (Right) Quantification of TMRE mean fluorescence intensity per mitochondrion in the AC versus UC, n = 9 animals. Paired two-tailed t-test, **** *p* < 0.0001. Scale bar, 5 μm.

**Figure S3.**
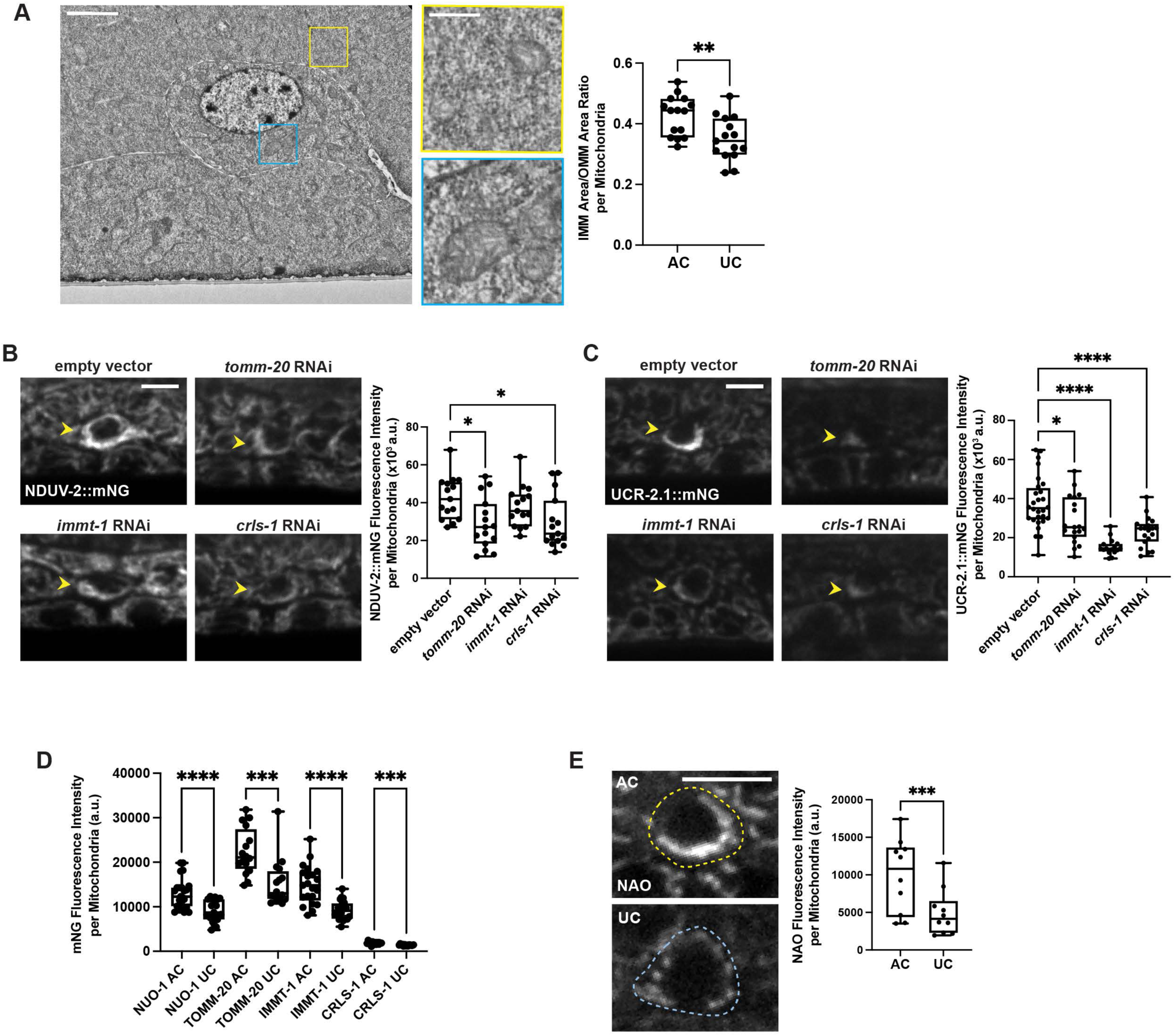
Cristae and import machinery in the AC, related to Figure 3. (**A**) (Left) Transmission electron microscopy (TEM) images of P6.p 2-cell stage AC (white outline), scale bar, 2 μm. Right panels show enlarged areas of AC mitochondria (blue box) and uterine cell (UC) mitochondria (yellow box) corresponding to boxed regions in left panel, scale bar 500 nm. (Right) Quantification of the ratio between inner membrane (IMM) area and outer membrane (OMM) area per mitochondrion in AC and UC. n = 15 mitochondria per cell type. Unpaired two-tailed t-test, ** *p* < 0.01. (**B**) (Left) NDUV-2::mNG in the AC (arrowhead) at the P6.p early 2-cell stage in empty vector control and *tomm-20*, *immt-1*, and *crls-1* RNAi-treated animals. (Right) Quantification of NDUV-2::mNG fluorescence intensity per mitochondrion in empty vector control and *tomm-20*, *immt-1*, and *crls-1* RNAi-treated animals, n = 15 animals for each condition. One-way ANOVA with Dunnett’s test for multiple comparisons, * *p* < 0.05. (**C**) (Left) UCR-2.1::mNG in the AC (arrowhead) at the P6.p early 2-cell stage in empty vector control and *tomm-20*, *immt-1*, and *crls-1* RNAi-treated animals. (Right) Quantification of UCR-2.1::mNG fluorescence intensity per mitochondrion in empty vector control and *tomm-20*, *immt-1*, and *crls-1* RNAi-treated animals, n ≥ 16 animals for each condition. One-Way ANOVA with Dunnett’s test for multiple comparisons, * *p* < 0.05, **** *p* < 0.0001. (**D**) Quantification of IMMT-1::mNG, TOMM-20::mNG, CRLS-1::mNG, and NUO-1::mNG fluorescence intensity per mitochondrion in the AC versus the uterine cell (UC), n ≥ 14 animals for each tagged component, AC and UC measurements made in the same animal. Mixed-effects ANOVA with Geisser-Greenhouse correction, followed by uncorrected Fisher’s LSD post hoc test, *** *p* < 0.001, **** *p* < 0.0001. (**E**) (Left) NAO staining of cardiolipin in the AC (top, yellow outline) and UC (bottom, blue outline) of P6.p 2-cell stage wild-type (N2) animals. (Right) Quantification of NAO fluorescence intensity per mitochondrion in the AC versus UC, n = 10 animals. Paired two-tailed t test, *** *p* < 0.001. Unless noted otherwise, scale bar, 5 μm.

**Figure S4.**
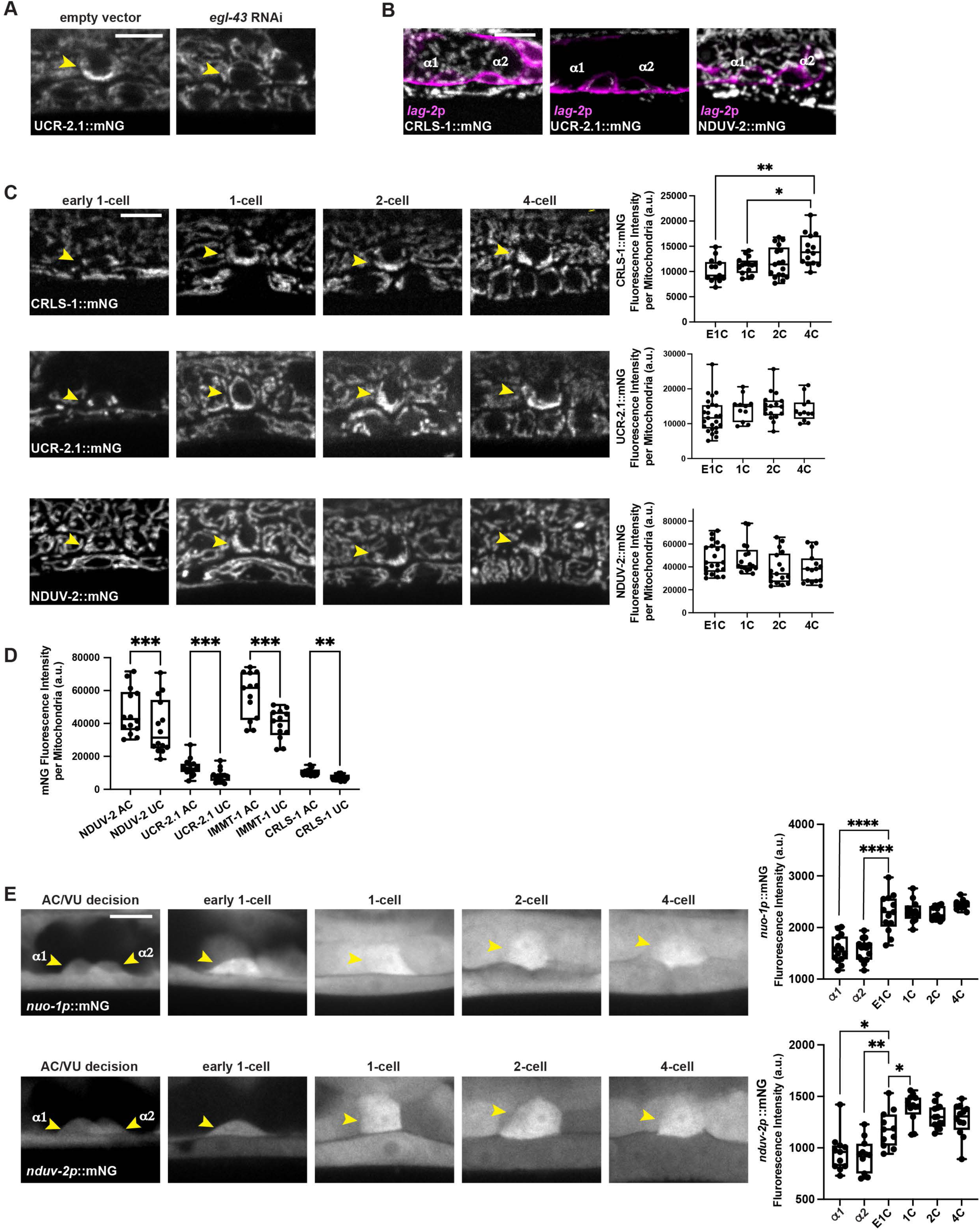
High-capacity mitochondria are specialized early during AC differentiation, related to Figure 4. (**A**) UCR-2.1::mNG in the AC (arrowhead) in empty vector control and *egl-43* RNAi-treated animals, corresponding to the quantification reported in Figure 4A. (**B**) CRLS-1::mNG (gray, left panel), UCR-2.1::mNG (gray, middle panel) and NDUV-2::mNG (grey, right panel) with *lag-2p*::2xmKate2::PH (magenta) in α1 and α2 during AC specification. (**C**) (Left) Developmental progression of CRLS-1::mNG (top panel), UCR-2.1::mNG (middle panel) and NDUV-2::mNG (bottom panel) expression in the AC (arrowhead) from the P6.p early 1-cell (E1C) to the P6.p 4-cell (4C) stage. (Right) Quantification of CRLS-1::mNG (top), UCR-2.1::mNG (middle) and NDUV-2::mNG (bottom) mean fluorescence intensity per AC mitochondria from the E1C to 4C stage, n ≥ 11 animals per stage for each strain. One-way ANOVA with Tukey’s post hoc test for multiple comparisons, * *p* < 0.05, ** *p* < 0.01. (**D**) Quantification of TOMM-20::mNG, IMMT-1::mNG, CRLS-1::mNG, NDUV-2::mNG, and UCR-2.1::mNG fluorescence intensity per mitochondrion in the AC versus uterine cell (UC) at the P6.p early 1-cell stage, n ≥ 12 animals per strain. Mixed-effects ANOVA with Geisser-Greenhouse correction, followed by uncorrected Fisher’s LSD post hoc test, ** *p* < 0.01, *** *p* < 0.001. (**E**) (Left) *nuo-1p*::mNG (top) and *nduv-2p*::mNG (bottom) transcriptional reporter in the AC (arrowhead) over developmental time from P6.p AC/VU decision to 4-cell stage. (Right) Quantification of *nuo-1p*::mNG (top) and *nduv-2p*::mNG (bottom) expression in the AC from the P6.p AC/VU decision to 4-cell stage, n ≥ 11 animals per strain per stage. At the AC/VU decision, both proto-uterine cells (α1 and α2) were measured because their cell fates are undetermined. One-way ANOVA with Tukey’s post hoc test for multiple comparisons, * *p* < 0.05, ** *p* < 0.01, **** *p* < 0.0001. Scale bar, 5 μm.

**Figure S5.**
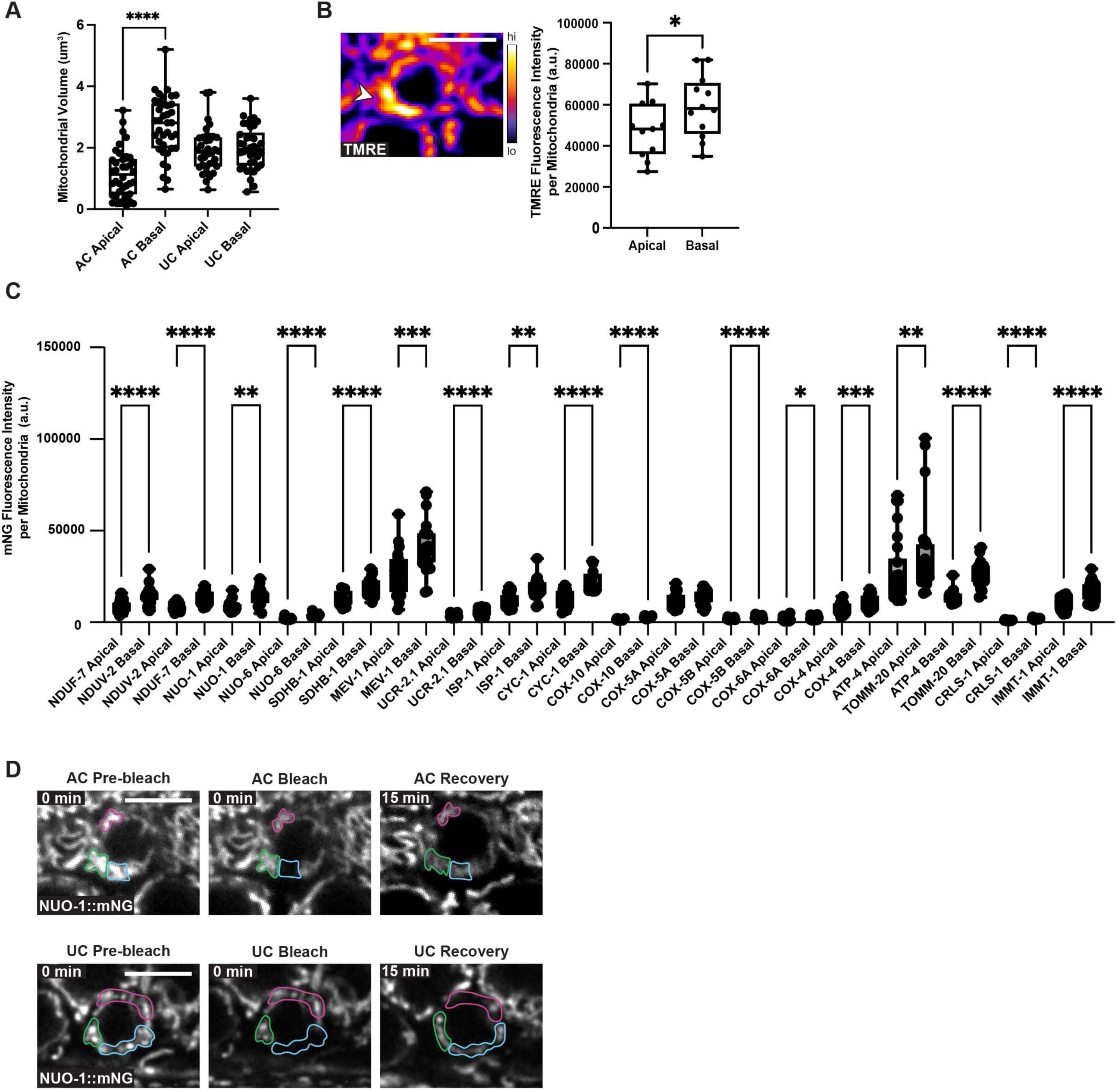
High-capacity mitochondria localize basally and remain separate from apical mitochondria, related to Figure 5. (**A**) Quantification of apical and basal mitochondrial volume in the AC and uterine cell (UC), n = 30. RM one-way ANOVA, with the Geisser-Greenhouse correction, followed by Sidák’s multiple comparisons test, with individual variances computed for each comparison, **** *p* > 0.0001. (**B**) (Left) Spectral fluorescence intensity maps of TMRE staining in the AC of P6.p 2-cell stage animals. Note the basal enrichment of TMRE (arrowhead). (Right) Quantification of TMRE fluorescence intensity per mitochondrion in the apical versus basal mitochondria in the AC, n = 12 animals. Paired two-tailed t test, * *p* < 0.05. (**C**) Quantification of fluorescence intensity of mNG tagged ETC components, CRLS-1::mNG, TOMM-20::mNG, and IMMT-1::mNG per mitochondrion in the apical versus basal mitochondria in P6.p 2-cell stage AC, n > 10 animals for each tagged component. Mixed-effects ANOVA with Geisser-Greenhouse correction, followed by uncorrected Fisher’s LSD post hoc test, * *p* < 0.05, ** *p* < 0.01, *** *p* < 0.001, **** *p* < 0.0001. (**D**) (Left) Additional examples of FRAP of NUO-1::mNG in the AC (top) and uterine cell (UC, bottom) mitochondria before, immediately after photobleaching, and 15 minutes post-photobleaching. Blue outline indicates bleached region, green outline indicates adjacent basal mitochondria, and magenta outline indicates apical mitochondria. Scale bar, 5 μm

**Figure S6.**
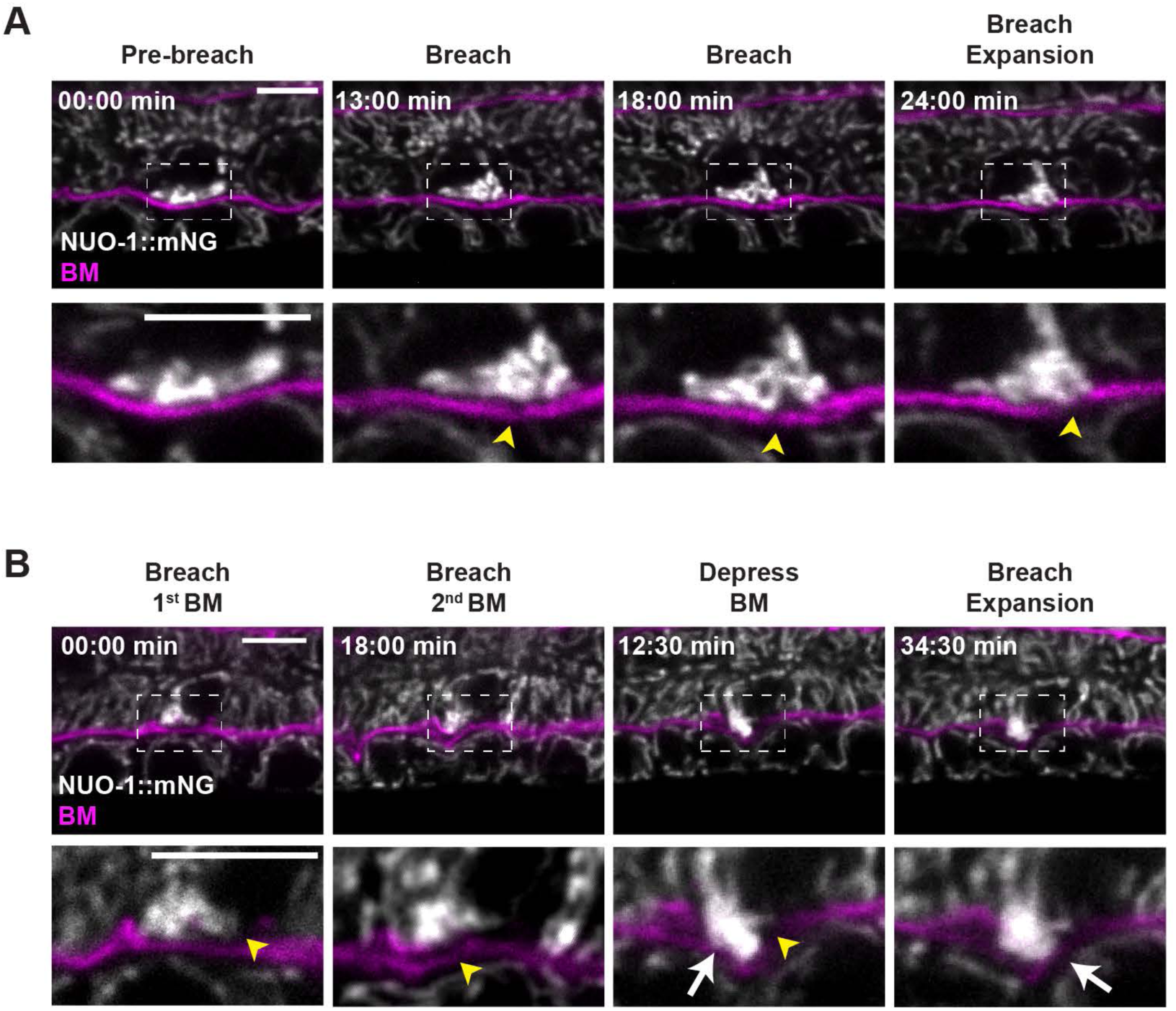
High-capacity mitochondria localize to the BM breach during invasion, related to Figure 6. (**A**) Time-lapse of AC mitochondria (NUO-1::mNG, gray) prior to BM (EMB-9::mRuby2, magenta) breaching, during the initial breach, and as the breach is expanding. Animals were imaged every 1 minute for 25 minutes. Bottom panels show enlarged areas corresponding to boxed regions in top panel. Yellow arrowheads indicate site of BM breaching. (**B**) Time-lapse of AC mitochondria (NUO-1::mNG, gray) during the BM (EMB-9::mRuby2, magenta) breaching, and as the breach is expanding. The BM underlying the AC is composed of two BM networks that are linked together during AC invasion^44^. Occasionally, the two previously linked BMs visibly separate. Animals were imaged every 1.5 minutes for 51 min. Bottom panels show enlarged areas corresponding to boxed regions in top panel. Yellow arrowheads indicate site of BM breaching. White arrow indicates mitochondria within the invasive protrusion. Scale bars, 5 μm.

**Figure S7.**
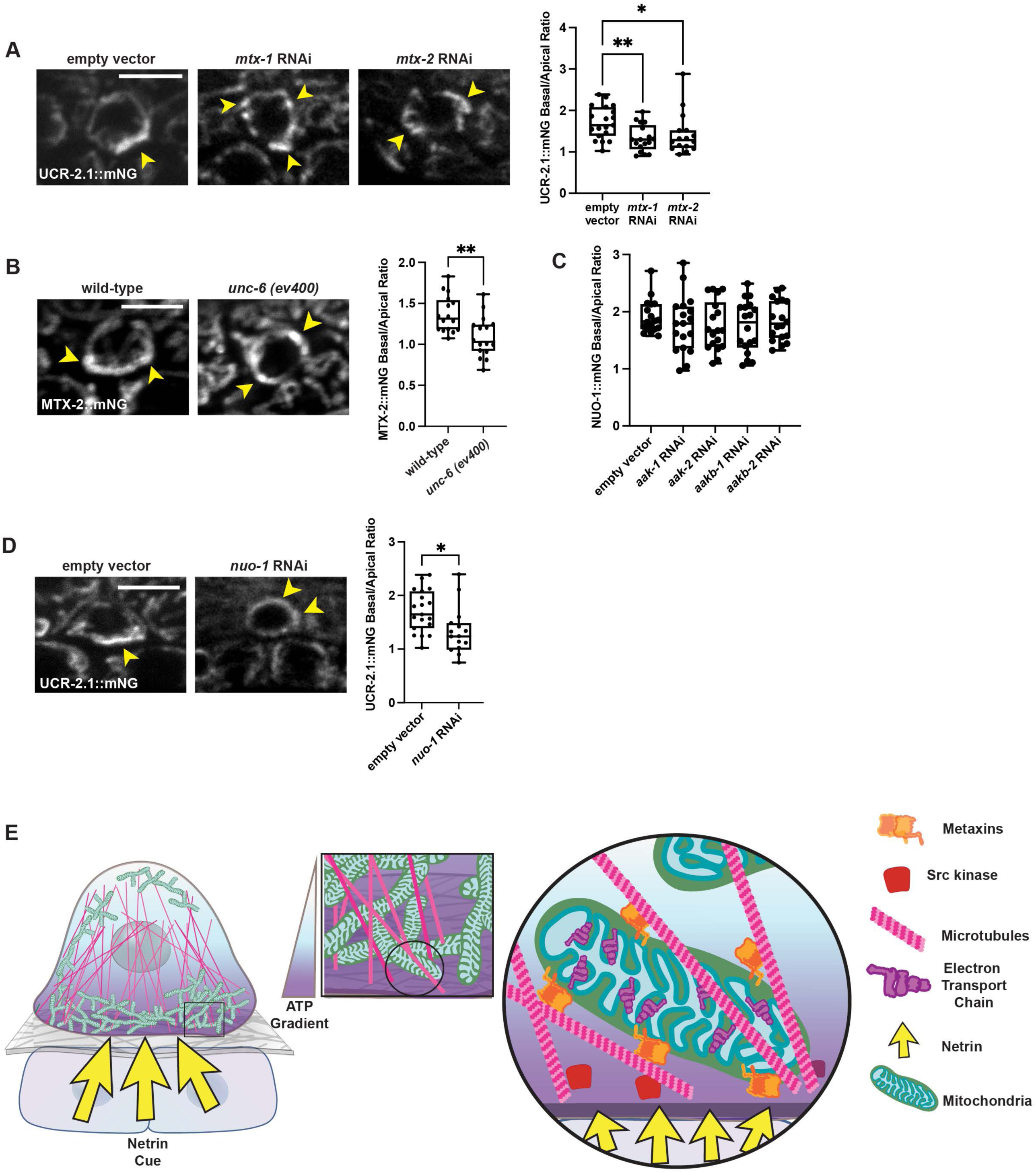
High-capacity mitochondria trafficking is dependent on metaxin adaptor complex, related to Figure 7. (**A**) (Left) UCR-2.1::mNG in the AC of empty vector control and *mtx-1* and *mtx-2* RNAi-treated animals at the P6.p early 2-cell stage. Yellow arrowhead indicates UCR-2.1::mNG enriched mitochondria. (Right) Quantification of basal to apical UCR-2.1::mNG fluorescence intensity per mitochondrion in empty vector control and *mtx-1* and *mtx-2* RNAi-treated animals, n = 19 animals. One-way ANOVA with Dunnett’s multiple comparison test, ** *p* < 0.01, * *p* < 0.05. (**B**) (Left) MTX-2::mNG in the AC of wild-type and netrin mutant (*unc-6 (ev400*)) animals at the P6.p 2-cell stage. Yellow arrowhead indicates site of MTX-2::mNG enrichment. (Right) Quantification of basal to apical MTX-2::mNG fluorescence intensity in wild-type versus *unc-6 (ev400)* mutant animals, n ≥ 14. Unpaired two-tailed t-test,** *p* < 0.01. (**C**) Quantification of basal to apical NUO-1::mNG fluorescence intensity per mitochondrion in empty vector control and AMPK subunit (*aak-1*, *aak-2*, *aakb-1*, and *aakb-2*) RNAi-treated animals, n =18. One-way ANOVA with Dunnett’s multiple comparison test found no significant differences between the basal/apical ratio of the control and AMPK subunit RNAi-treated animals. (**D**) (Left) UCR-2.1::mNG in the AC of empty vector and *nuo-1* RNAi-treated animals at the P6.p 2-cell stage. Yellow arrowhead indicates UCR-2.1::mNG enriched mitochondria. (Right) Quantification of basal to apical UCR-2.1::mNG fluorescence intensity in empty vector and *nuo-1* RNAi-treated animals, n ≥ 15 animals. Unpaired two-tailed t-test, * *p* < 0.05. (**E**) Schematic of AC mitochondrial localization dependent on the netrin directional cue, microtubules, metaxins, and Src kinase. Basal localization of mitochondria results high levels of ATP at the invasive front. Scale bar, 5 μm.

**Movie S1.** Time-lapse of mitochondria (NUO-1::mNG, gray) prior to and during AC invasion through the BM (EMB-9::mRuby2, magenta) and in the emerging protrusion. Animals were imaged every 1 min for 31 min.

**Movie S2.** Time-lapse of mitochondria (NUO-1::mNG, gray) prior to and during AC invasion through the BM (EMB-9::mRuby2, magenta) and in the emerging protrusion. Animals were imaged every 1 min for 25 min.

**Movie S3** Mitochondria (NUO-1::mNG, gray) during AC invasion through the BM (EMB-9::mRuby2, magenta) and in the emerging protrusion. Animals were imaged every 1:30 min for 51 min.

Scale bar, 5 µm.

## Key Resources Table

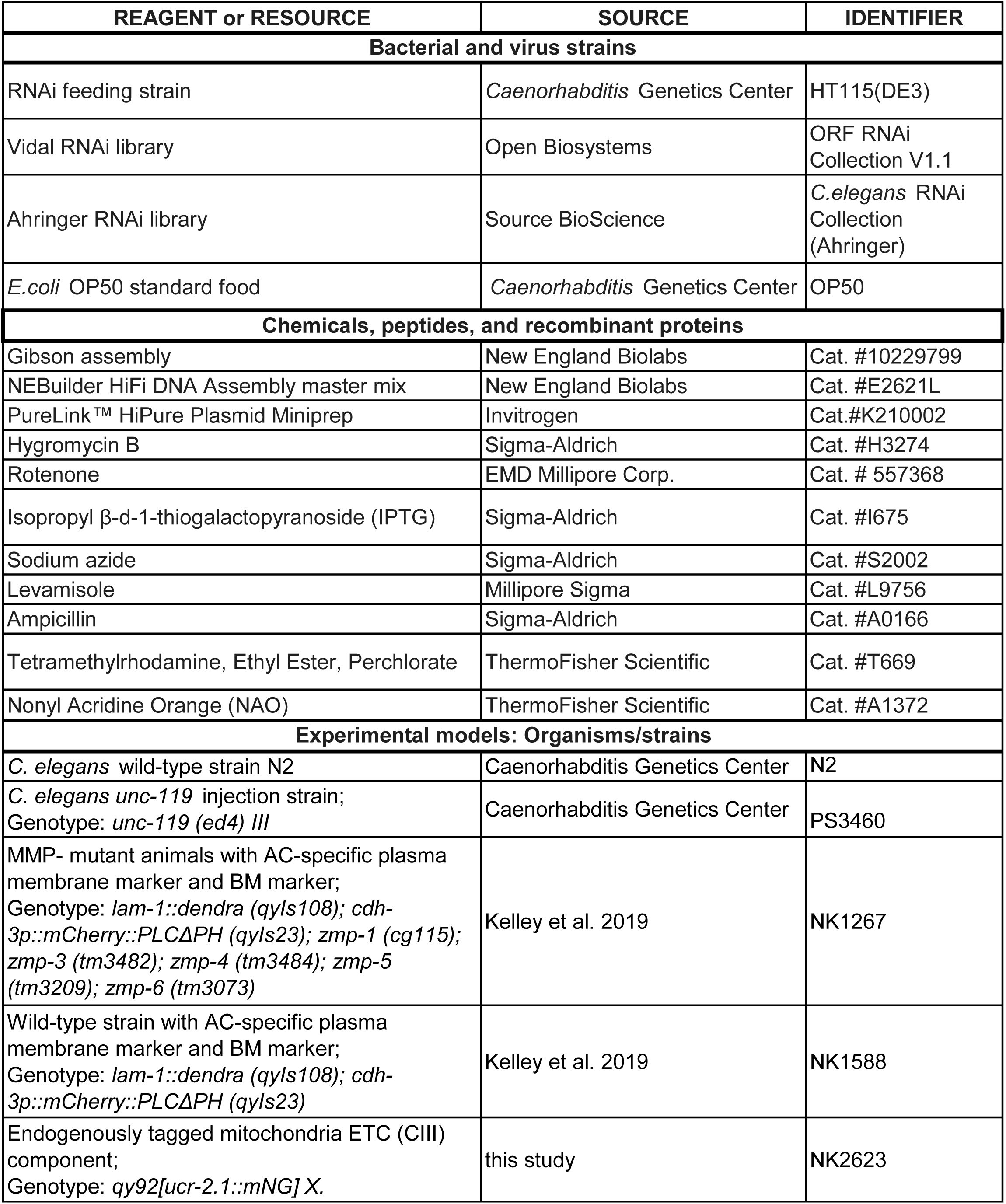
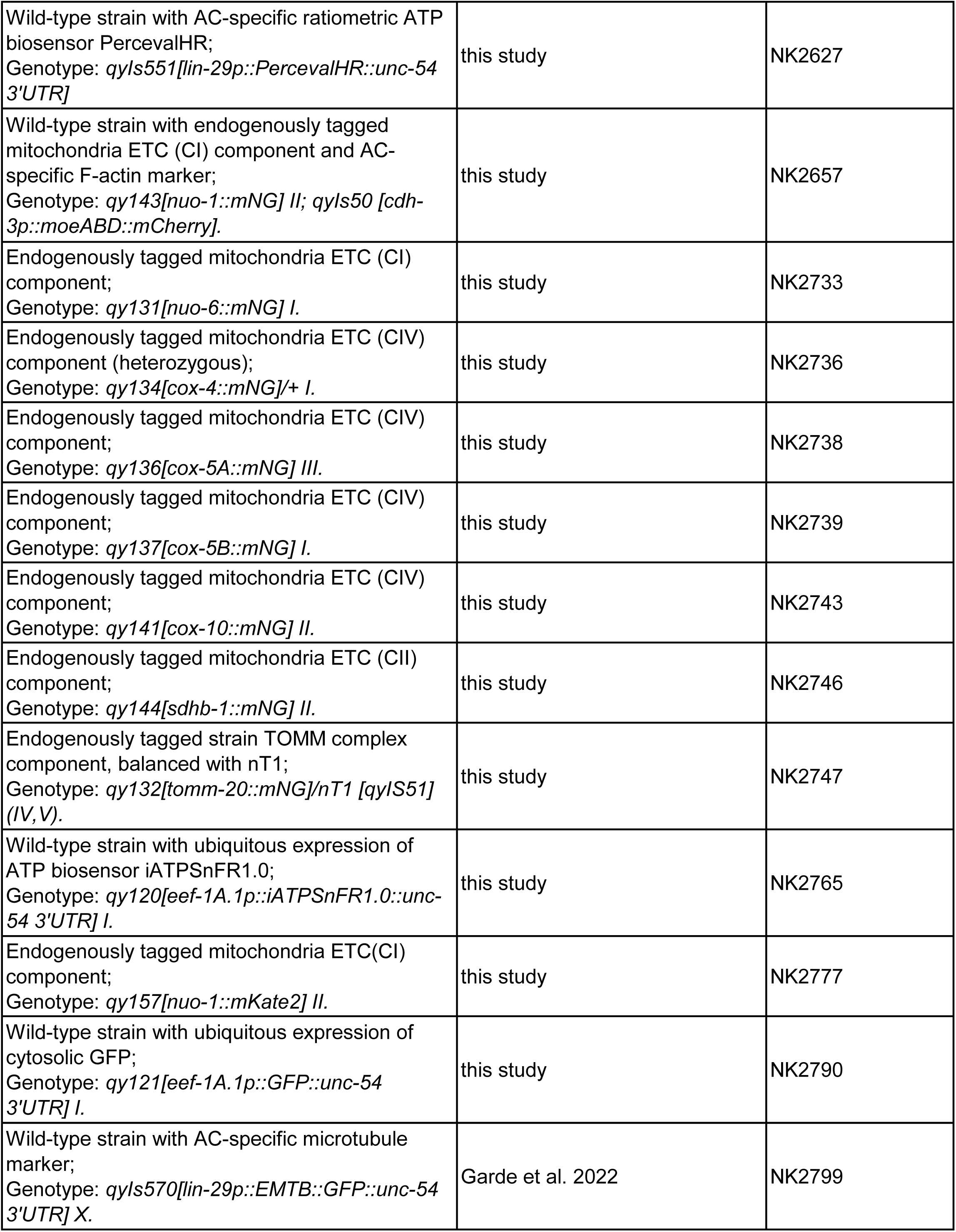
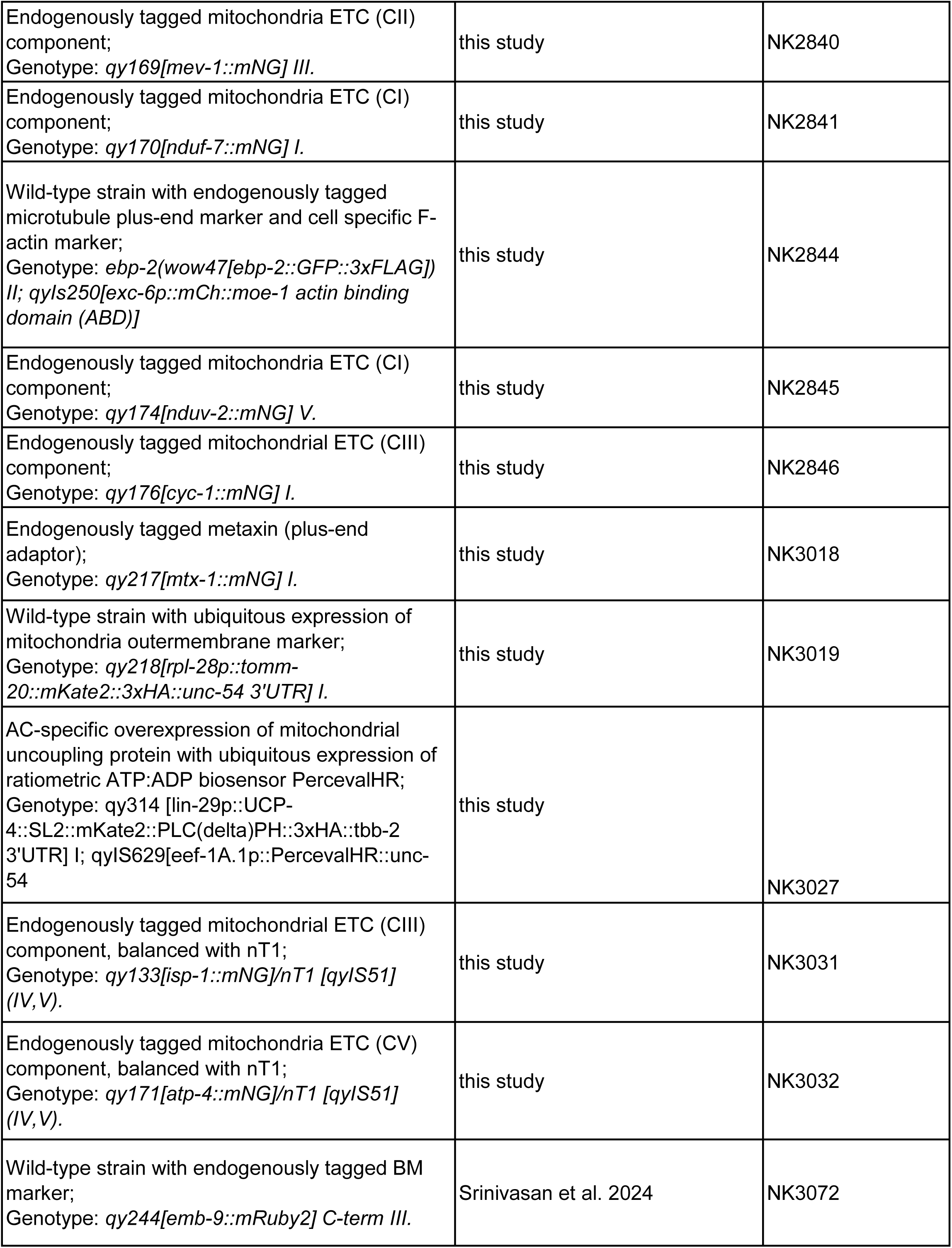
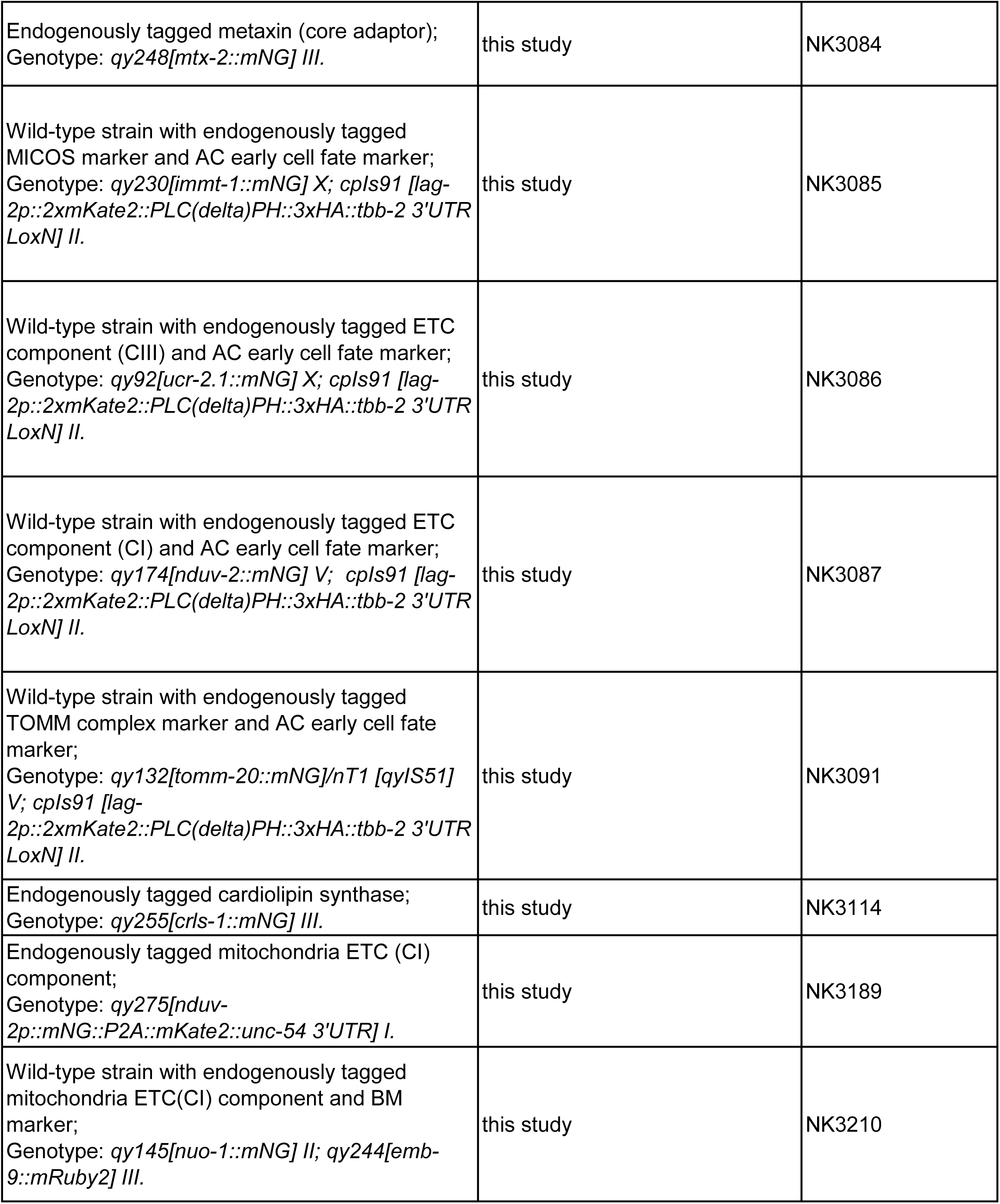
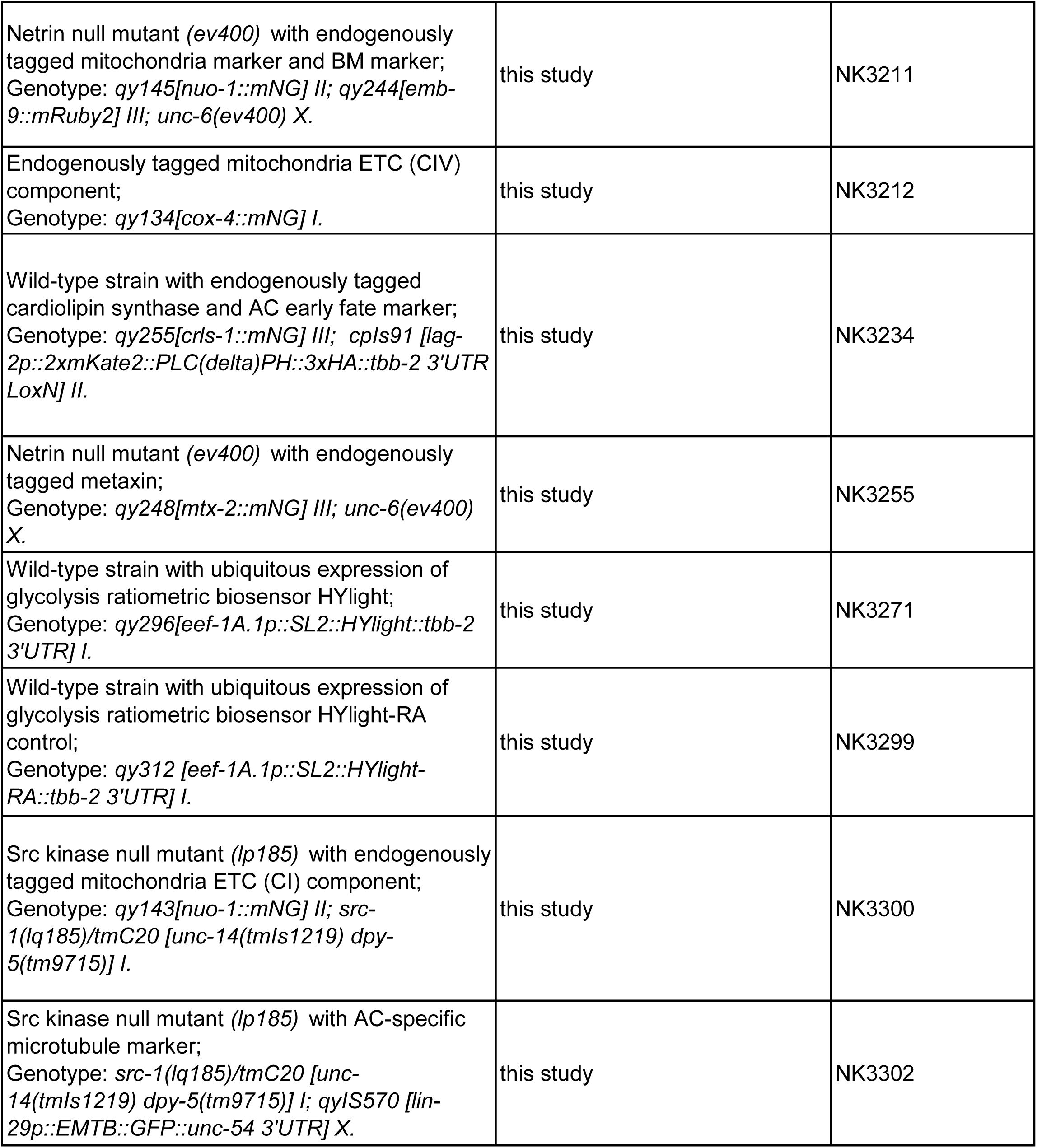
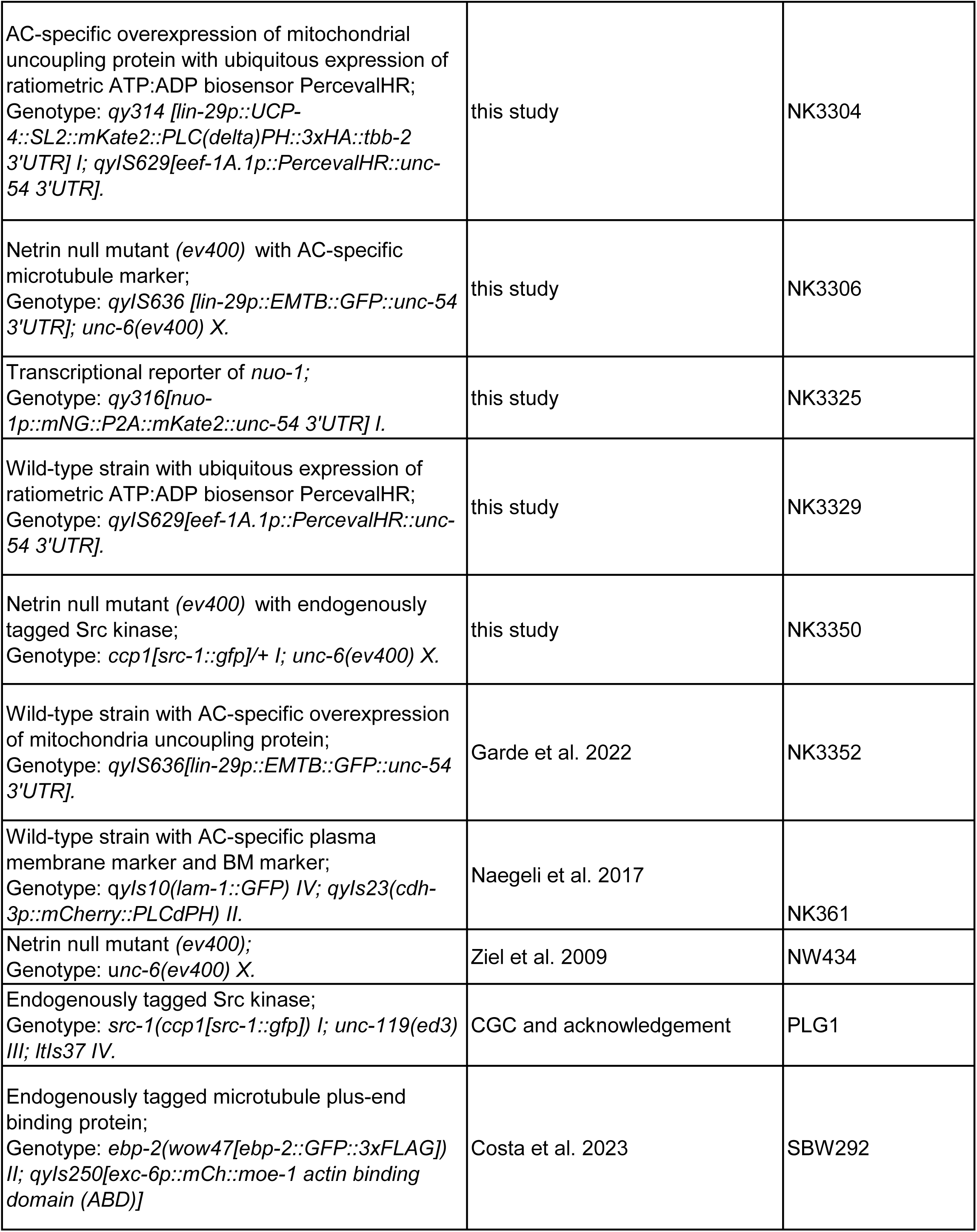
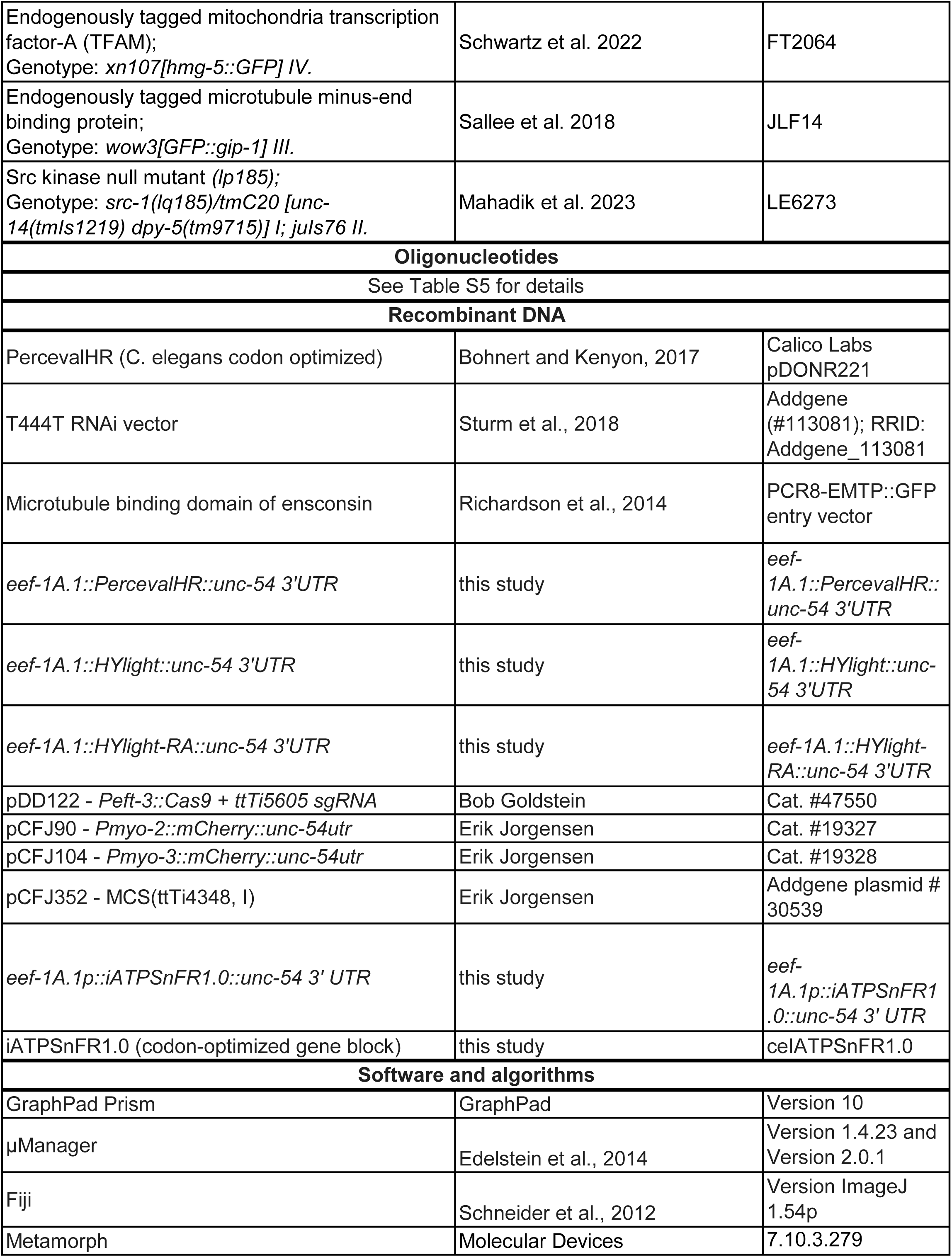
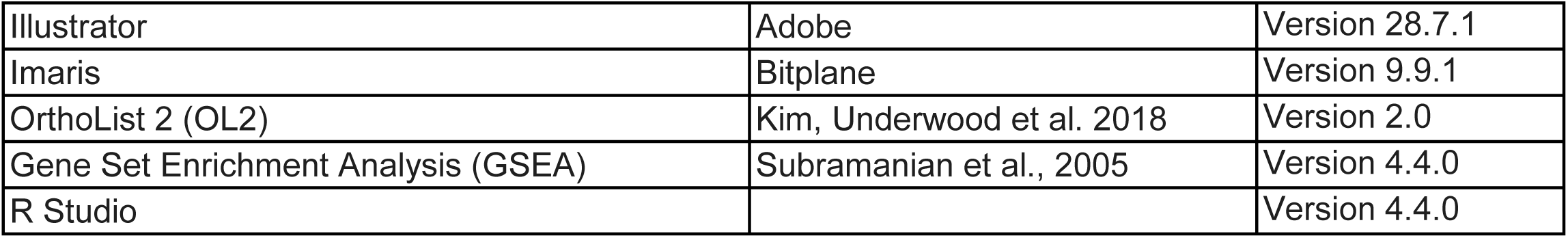

---

**Table S1.**
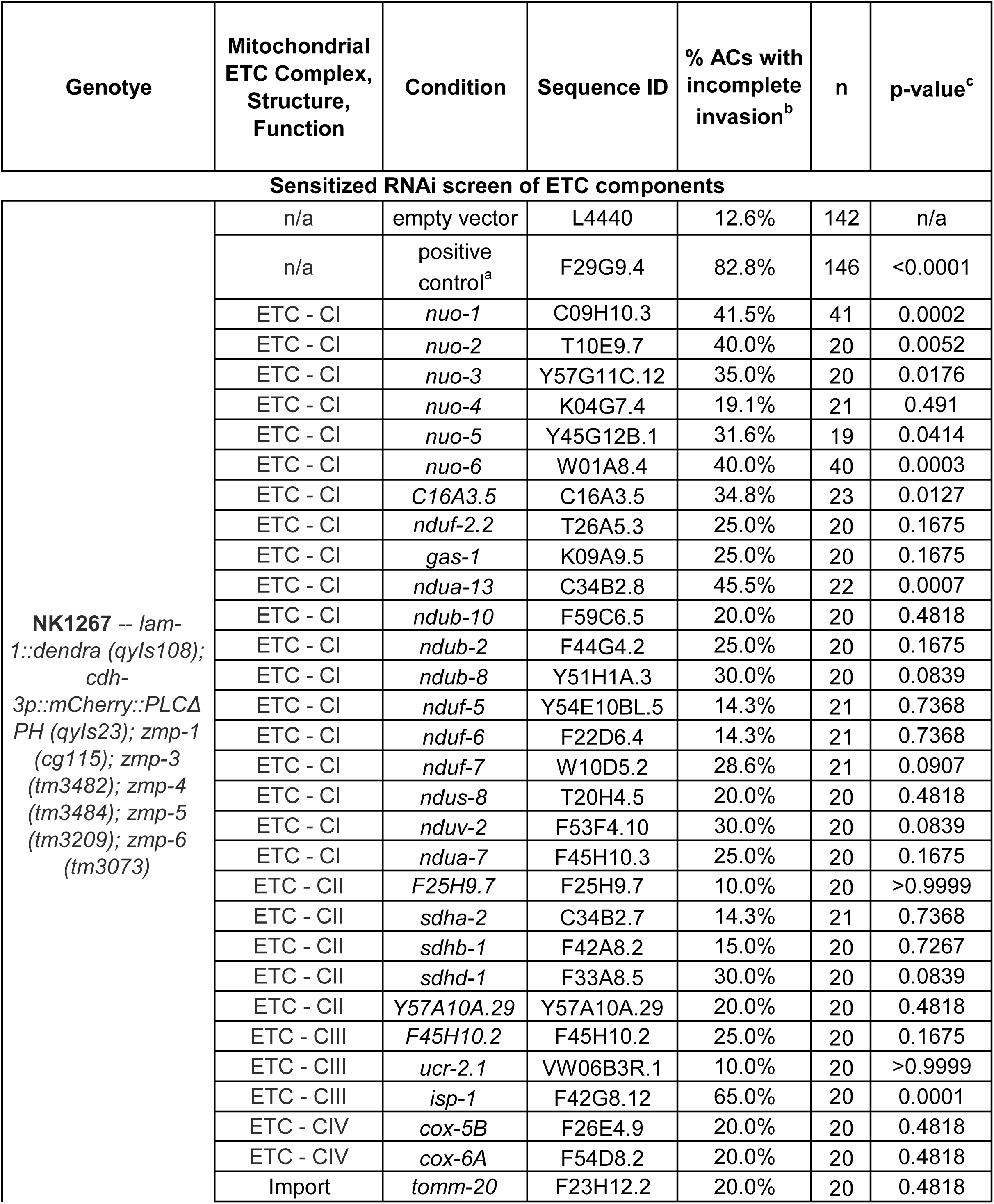
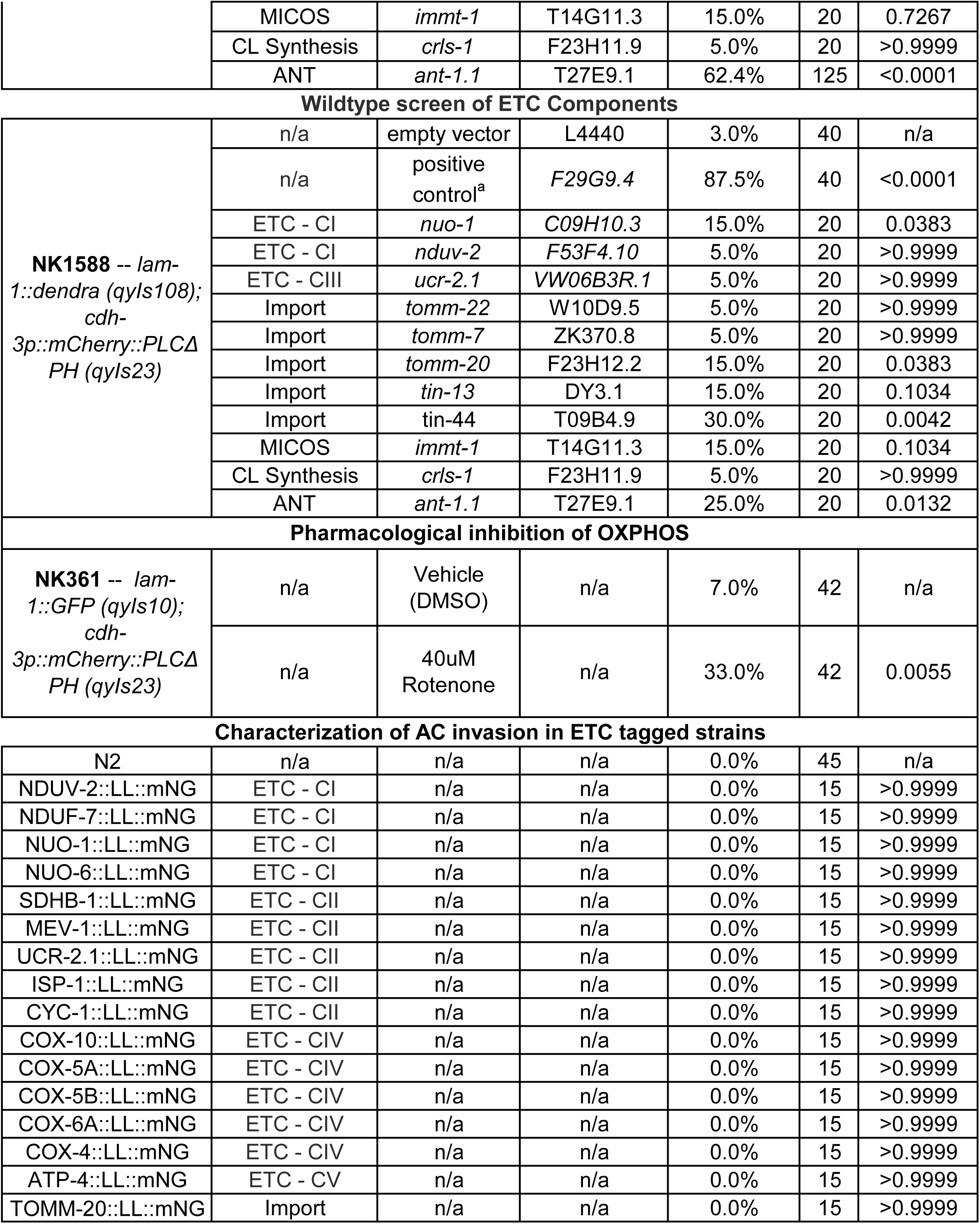
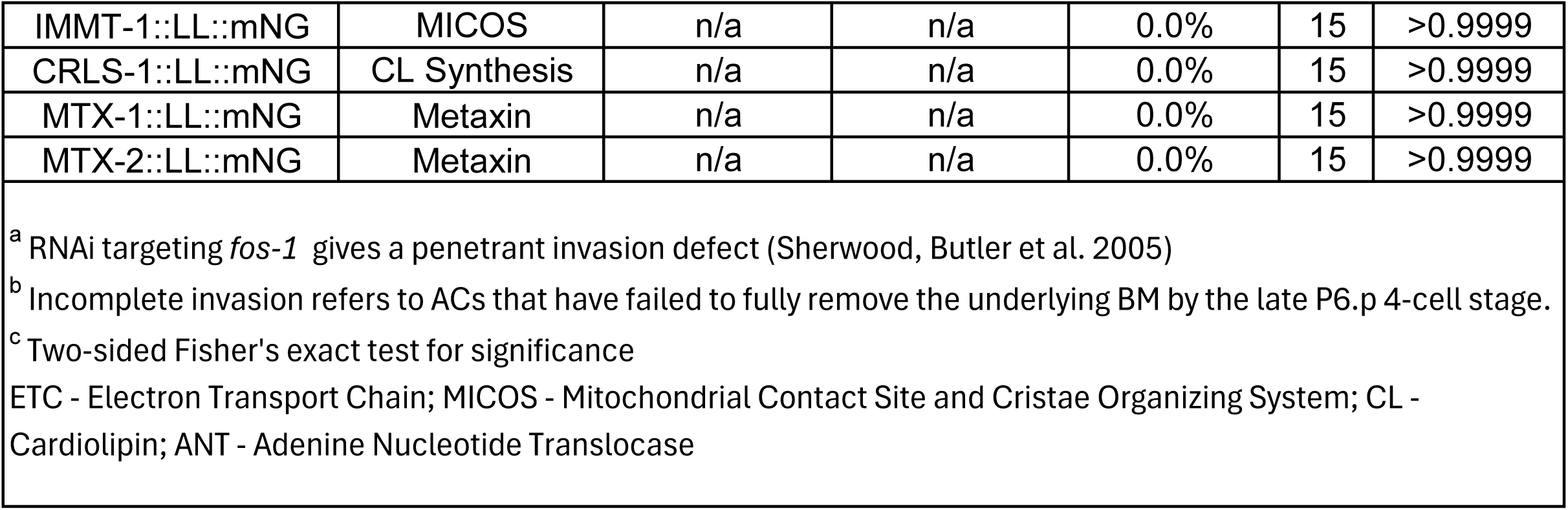
AC invasion scoring.

| Genotype | Mitochondrial ETC Complex, Structure, Function | Condition | Sequence ID | % ACs with incomplete invasion<sup>b</sup> | n | p-value<sup>c</sup> |
| --- | --- | --- | --- | --- | --- | --- |
| **Sensitized RNAi screen of ETC components** |  |  |  |  |  |  |
| **NK1267** -- *lam-1::dendra (qyls108); cdh-3p::mCherry::PLCΔ PH (qyls23); zmp-1 (cg115); zmp-3 (tm3482); zmp-4 (tm3484); zmp-5 (tm3209); zmp-6 (tm3073)* | n/a | empty vector | L4440 | 12.6% | 142 | n/a |
|  | n/a | positive control<sup>a</sup> | F29G9.4 | 82.8% | 146 | <0.0001 |
|  | ETC - CI | *nuo-1* | C09H10.3 | 41.5% | 41 | 0.0002 |
|  | ETC - CI | *nuo-2* | T10E9.7 | 40.0% | 20 | 0.0052 |
|  | ETC - CI | *nuo-3* | Y57G11C.12 | 35.0% | 20 | 0.0176 |
|  | ETC - CI | *nuo-4* | K04G7.4 | 19.1% | 21 | 0.491 |
|  | ETC - CI | *nuo-5* | Y45G12B.1 | 31.6% | 19 | 0.0414 |
|  | ETC - CI | *nuo-6* | W01A8.4 | 40.0% | 40 | 0.0003 |
|  | ETC - CI | *C16A3.5* | C16A3.5 | 34.8% | 23 | 0.0127 |
|  | ETC - CI | *nduf-2.2* | T26A5.3 | 25.0% | 20 | 0.1675 |
|  | ETC - CI | *gas-1* | K09A9.5 | 25.0% | 20 | 0.1675 |
|  | ETC - CI | *ndua-13* | C34B2.8 | 45.5% | 22 | 0.0007 |
|  | ETC - CI | *ndub-10* | F59C6.5 | 20.0% | 20 | 0.4818 |
|  | ETC - CI | *ndub-2* | F44G4.2 | 25.0% | 20 | 0.1675 |
|  | ETC - CI | *ndub-8* | Y51H1A.3 | 30.0% | 20 | 0.0839 |
|  | ETC - CI | *nduf-5* | Y54E10BL.5 | 14.3% | 21 | 0.7368 |
|  | ETC - CI | *nduf-6* | F22D6.4 | 14.3% | 21 | 0.7368 |
|  | ETC - CI | *nduf-7* | W10D5.2 | 28.6% | 21 | 0.0907 |
|  | ETC - CI | *ndus-8* | T20H4.5 | 20.0% | 20 | 0.4818 |
|  | ETC - CI | *nduv-2* | F53F4.10 | 30.0% | 20 | 0.0839 |
|  | ETC - CI | *ndua-7* | F45H10.3 | 25.0% | 20 | 0.1675 |
|  | ETC - CII | *F25H9.7* | F25H9.7 | 10.0% | 20 | >0.9999 |
|  | ETC - CII | *sdha-2* | C34B2.7 | 14.3% | 21 | 0.7368 |
|  | ETC - CII | *sdhb-1* | F42A8.2 | 15.0% | 20 | 0.7267 |
|  | ETC - CII | *sdhd-1* | F33A8.5 | 30.0% | 20 | 0.0839 |
|  | ETC - CII | *Y57A10A.29* | Y57A10A.29 | 20.0% | 20 | 0.4818 |
|  | ETC - CIII | *F45H10.2* | F45H10.2 | 25.0% | 20 | 0.1675 |
|  | ETC - CIII | *ucr-2.1* | VW06B3R.1 | 10.0% | 20 | >0.9999 |
|  | ETC - CIII | *isp-1* | F42G8.12 | 65.0% | 20 | 0.0001 |
|  | ETC - CIV | *cox-5B* | F26E4.9 | 20.0% | 20 | 0.4818 |
|  | ETC - CIV | *cox-6A* | F54D8.2 | 20.0% | 20 | 0.4818 |
|  | Import | *tomm-20* | F23H12.2 | 20.0% | 20 | 0.4818 |
|  | MICOS | *immt-1* | T14G11.3 | 15.0% | 20 | 0.7267 |
|  | CL Synthesis | *crls-1* | F23H11.9 | 5.0% | 20 | >0.9999 |
|  | ANT | *ant-1.1* | T27E9.1 | 62.4% | 125 | <0.0001 |
| Wildtype screen of ETC Components |  |  |  |  |  |  |
| **NK1588** -- *lam-1::dendra (qyls108); cdh-3p::mCherry::PLCΔ PH (qyls23)* | n/a | empty vector | L4440 | 3.0% | 40 | n/a |
|  | n/a | positive control<sup>a</sup> | *F29G9.4* | 87.5% | 40 | <0.0001 |
|  | ETC - CI | *nuo-1* | *C09H10.3* | 15.0% | 20 | 0.0383 |
|  | ETC - CI | *nduv-2* | *F53F4.10* | 5.0% | 20 | >0.9999 |
|  | ETC - CIII | *ucr-2.1* | *VW06B3R.1* | 5.0% | 20 | >0.9999 |
|  | Import | *tomm-22* | W10D9.5 | 5.0% | 20 | >0.9999 |
|  | Import | *tomm-7* | ZK370.8 | 5.0% | 20 | >0.9999 |
|  | Import | *tomm-20* | F23H12.2 | 15.0% | 20 | 0.0383 |
|  | Import | *tin-13* | DY3.1 | 15.0% | 20 | 0.1034 |
|  | Import | *tin-44* | T09B4.9 | 30.0% | 20 | 0.0042 |
|  | MICOS | *immt-1* | T14G11.3 | 15.0% | 20 | 0.1034 |
|  | CL Synthesis | *crls-1* | F23H11.9 | 5.0% | 20 | >0.9999 |
|  | ANT | *ant-1.1* | T27E9.1 | 25.0% | 20 | 0.0132 |
| Pharmacological inhibition of OXPHOS |  |  |  |  |  |  |
| **NK361** -- *lam-1::GFP (qyls10); cdh-3p::mCherry::PLCΔ PH (qyls23)* | n/a | Vehicle (DMSO) | n/a | 7.0% | 42 | n/a |
|  | n/a | 40uM Rotenone | n/a | 33.0% | 42 | 0.0055 |
| Characterization of AC invasion in ETC tagged strains |  |  |  |  |  |  |
| N2 | n/a | n/a | n/a | 0.0% | 45 | n/a |
| NDUV-2::LL::mNG | ETC - CI | n/a | n/a | 0.0% | 15 | >0.9999 |
| NDUF-7::LL::mNG | ETC - CI | n/a | n/a | 0.0% | 15 | >0.9999 |
| NUO-1::LL::mNG | ETC - CI | n/a | n/a | 0.0% | 15 | >0.9999 |
| NUO-6::LL::mNG | ETC - CI | n/a | n/a | 0.0% | 15 | >0.9999 |
| SDHB-1::LL::mNG | ETC - CII | n/a | n/a | 0.0% | 15 | >0.9999 |
| MEV-1::LL::mNG | ETC - CII | n/a | n/a | 0.0% | 15 | >0.9999 |
| UCR-2.1::LL::mNG | ETC - CII | n/a | n/a | 0.0% | 15 | >0.9999 |
| ISP-1::LL::mNG | ETC - CII | n/a | n/a | 0.0% | 15 | >0.9999 |
| CYC-1::LL::mNG | ETC - CII | n/a | n/a | 0.0% | 15 | >0.9999 |
| COX-10::LL::mNG | ETC - CIV | n/a | n/a | 0.0% | 15 | >0.9999 |
| COX-5A::LL::mNG | ETC - CIV | n/a | n/a | 0.0% | 15 | >0.9999 |
| COX-5B::LL::mNG | ETC - CIV | n/a | n/a | 0.0% | 15 | >0.9999 |
| COX-6A::LL::mNG | ETC - CIV | n/a | n/a | 0.0% | 15 | >0.9999 |
| COX-4::LL::mNG | ETC - CIV | n/a | n/a | 0.0% | 15 | >0.9999 |
| ATP-4::LL::mNG | ETC - CV | n/a | n/a | 0.0% | 15 | >0.9999 |
| TOMM-20::LL::mNG | Import | n/a | n/a | 0.0% | 15 | >0.9999 |
| IMMT-1::LL::mNG | MICOS | n/a | n/a | 0.0% | 15 | >0.9999 |
| CRLS-1::LL::mNG | CL Synthesis | n/a | n/a | 0.0% | 15 | >0.9999 |
| MTX-1::LL::mNG | Metaxin | n/a | n/a | 0.0% | 15 | >0.9999 |
| MTX-2::LL::mNG | Metaxin | n/a | n/a | 0.0% | 15 | >0.9999 |

<sup>a</sup> RNAi targeting *fos-1* gives a penetrant invasion defect (Sherwood, Butler et al. 2005)
<sup>b</sup> Incomplete invasion refers to ACs that have failed to fully remove the underlying BM by the late P6.p 4-cell stage.
<sup>c</sup> Two-sided Fisher's exact test for significance
ETC - Electron Transport Chain; MICOS - Mitochondrial Contact Site and Cristae Organizing System; CL - Cardiolipin; ANT - Adenine Nucleotide Translocase

**Table S2.**
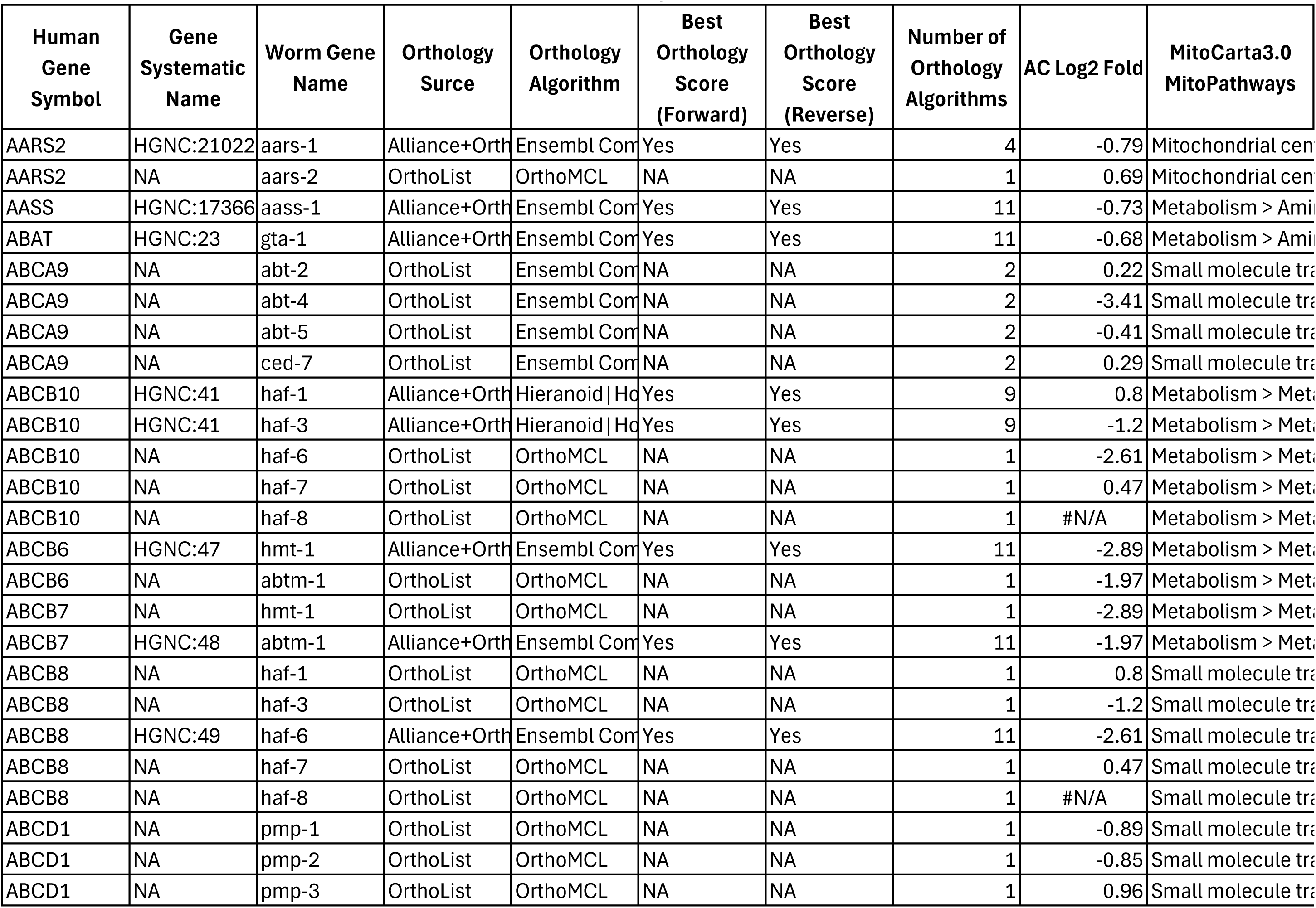
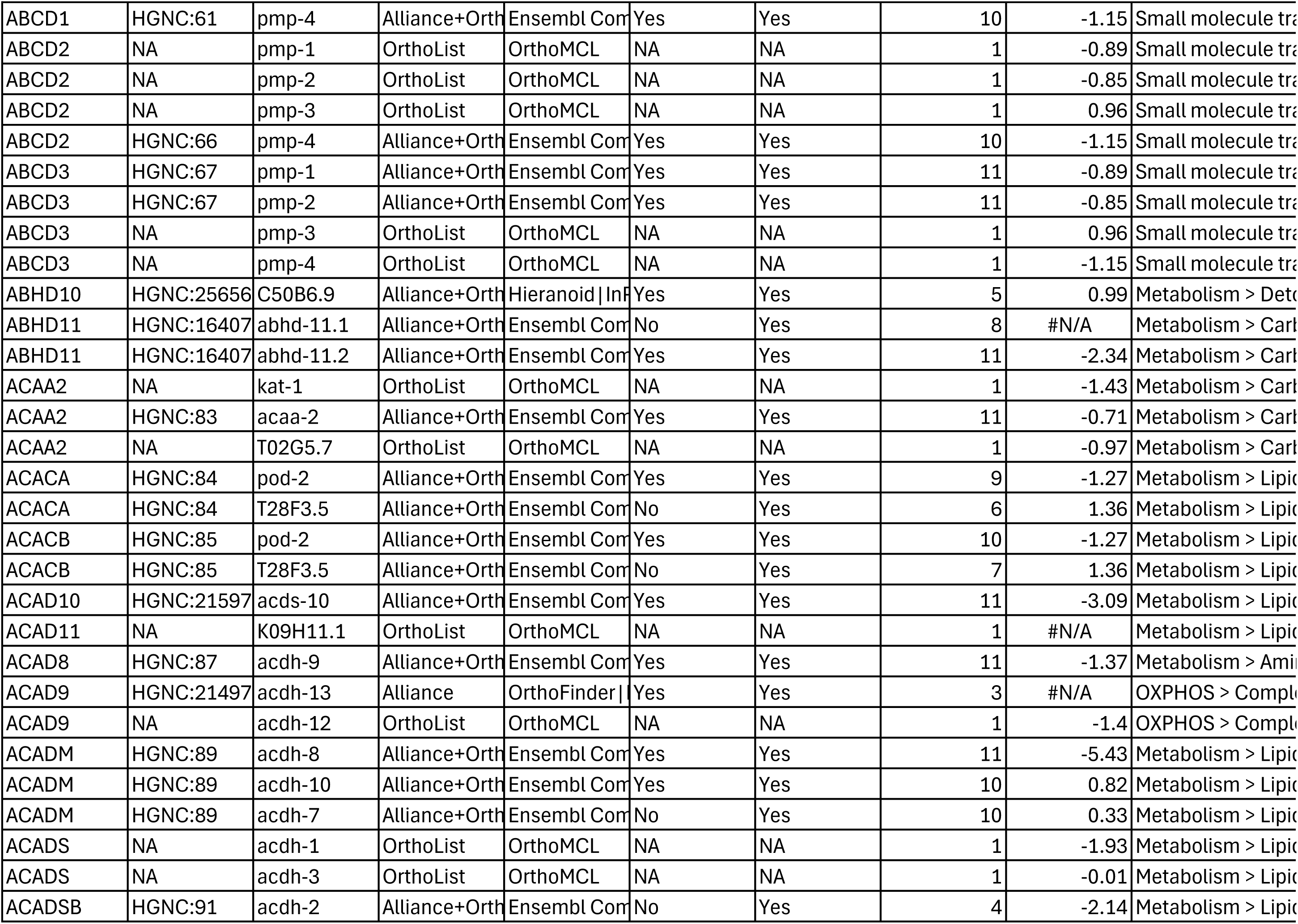
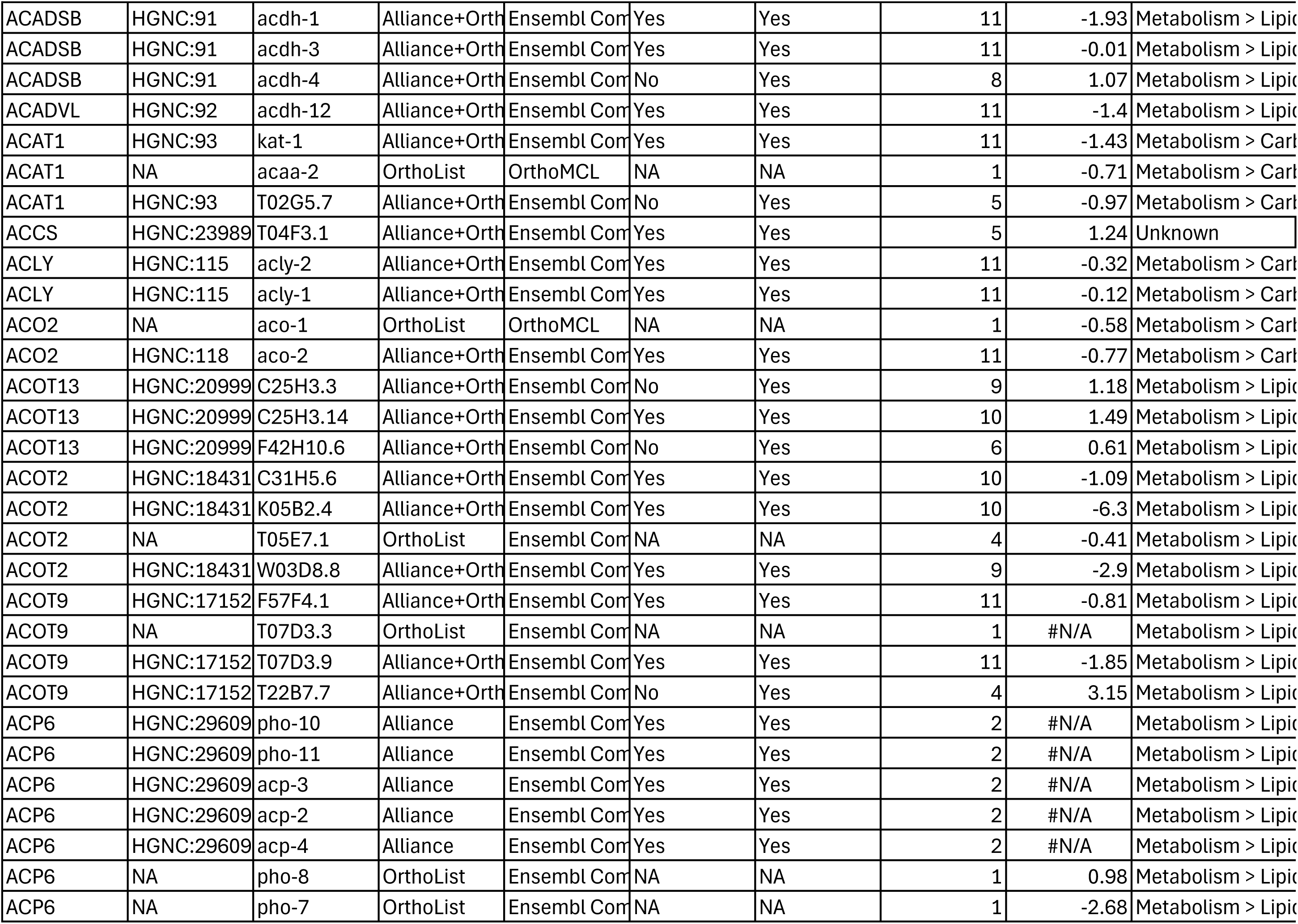
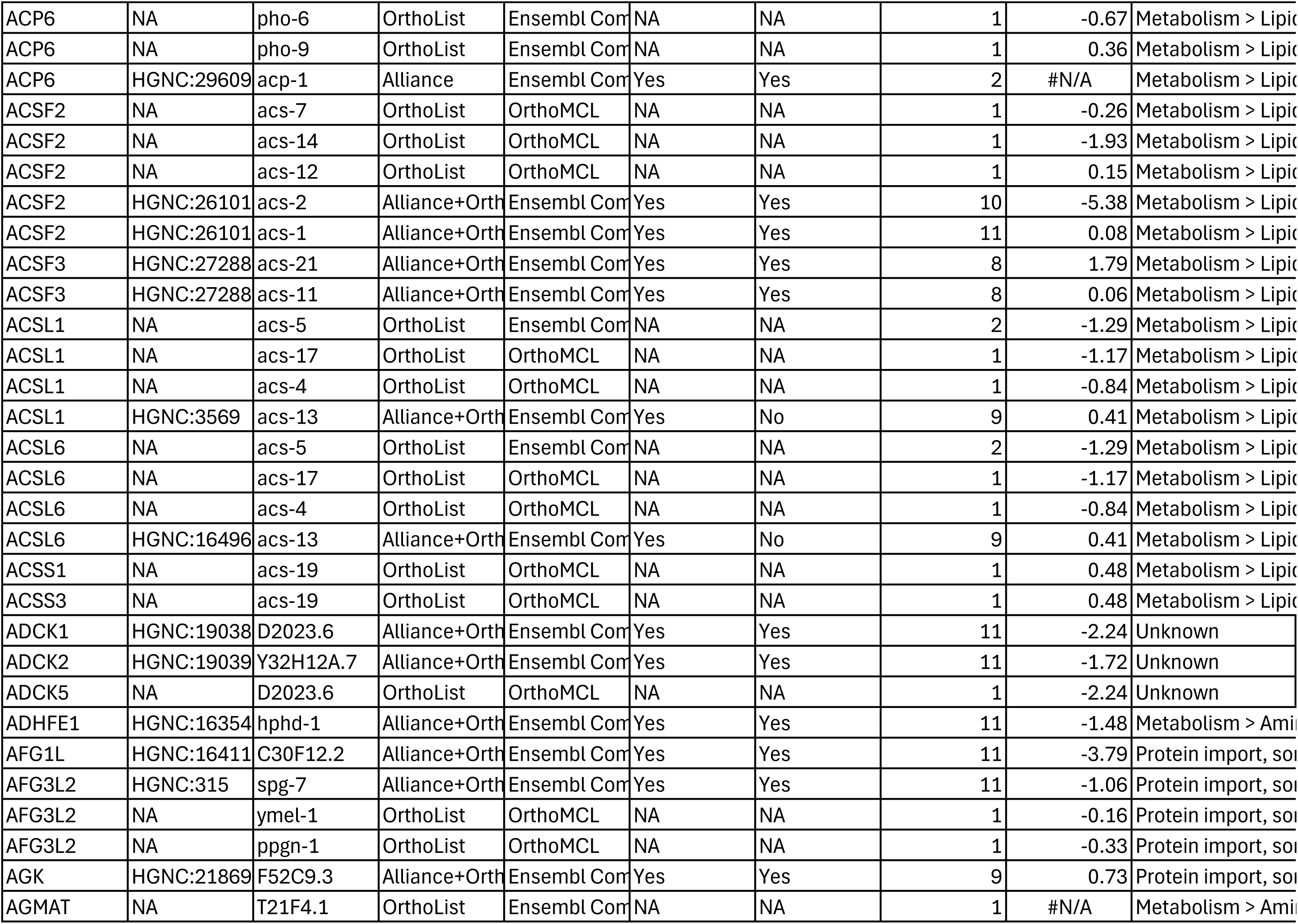
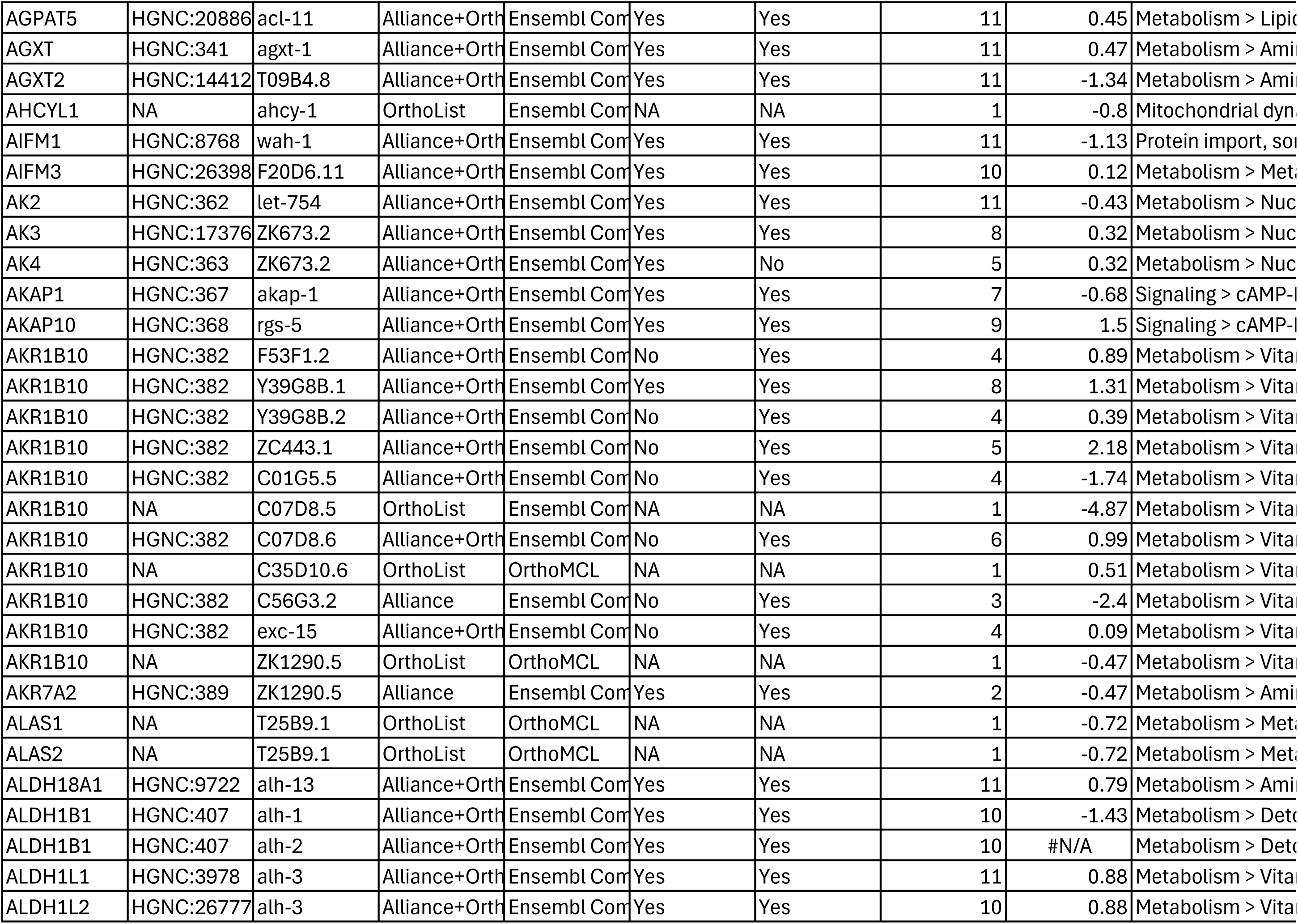
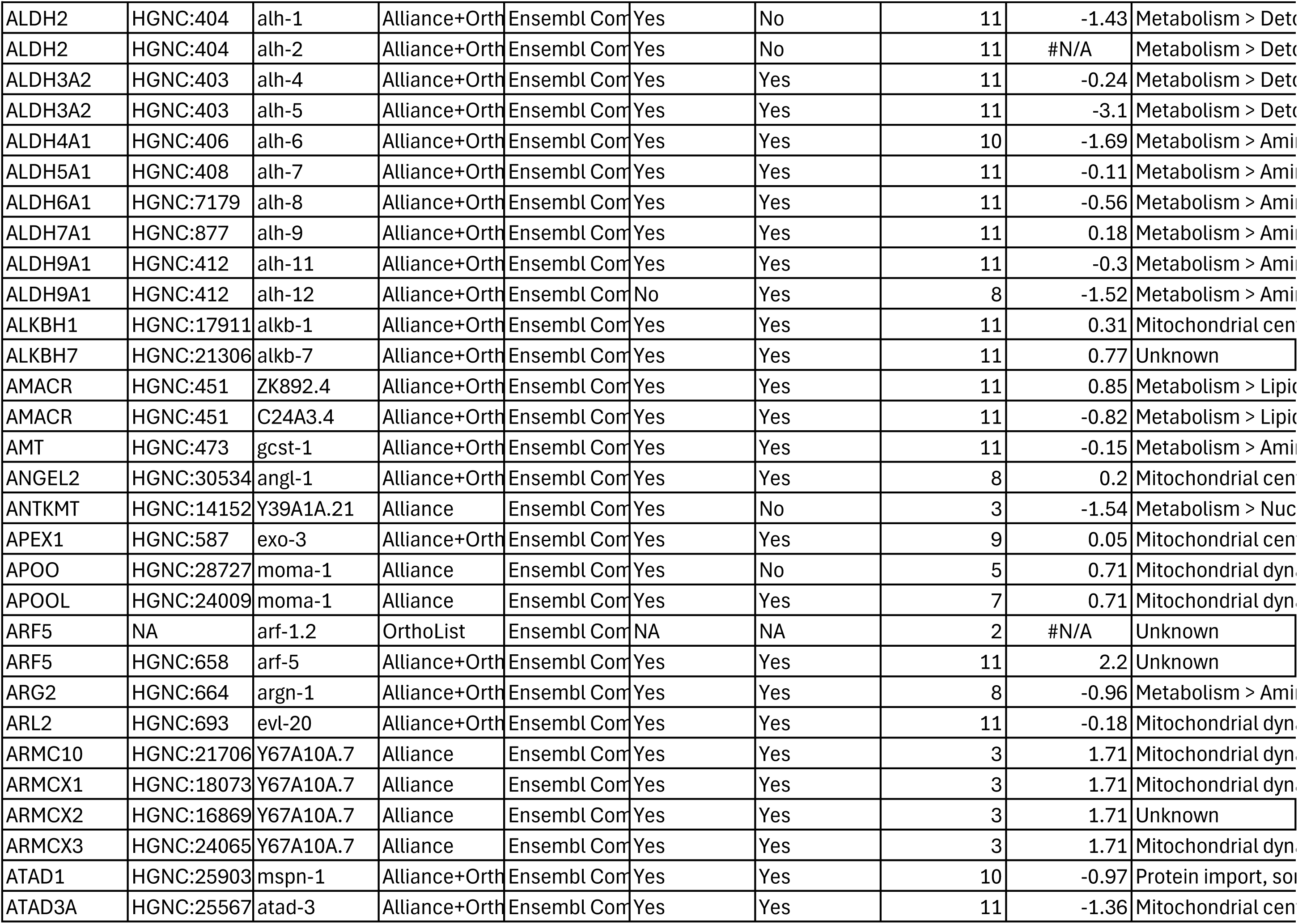
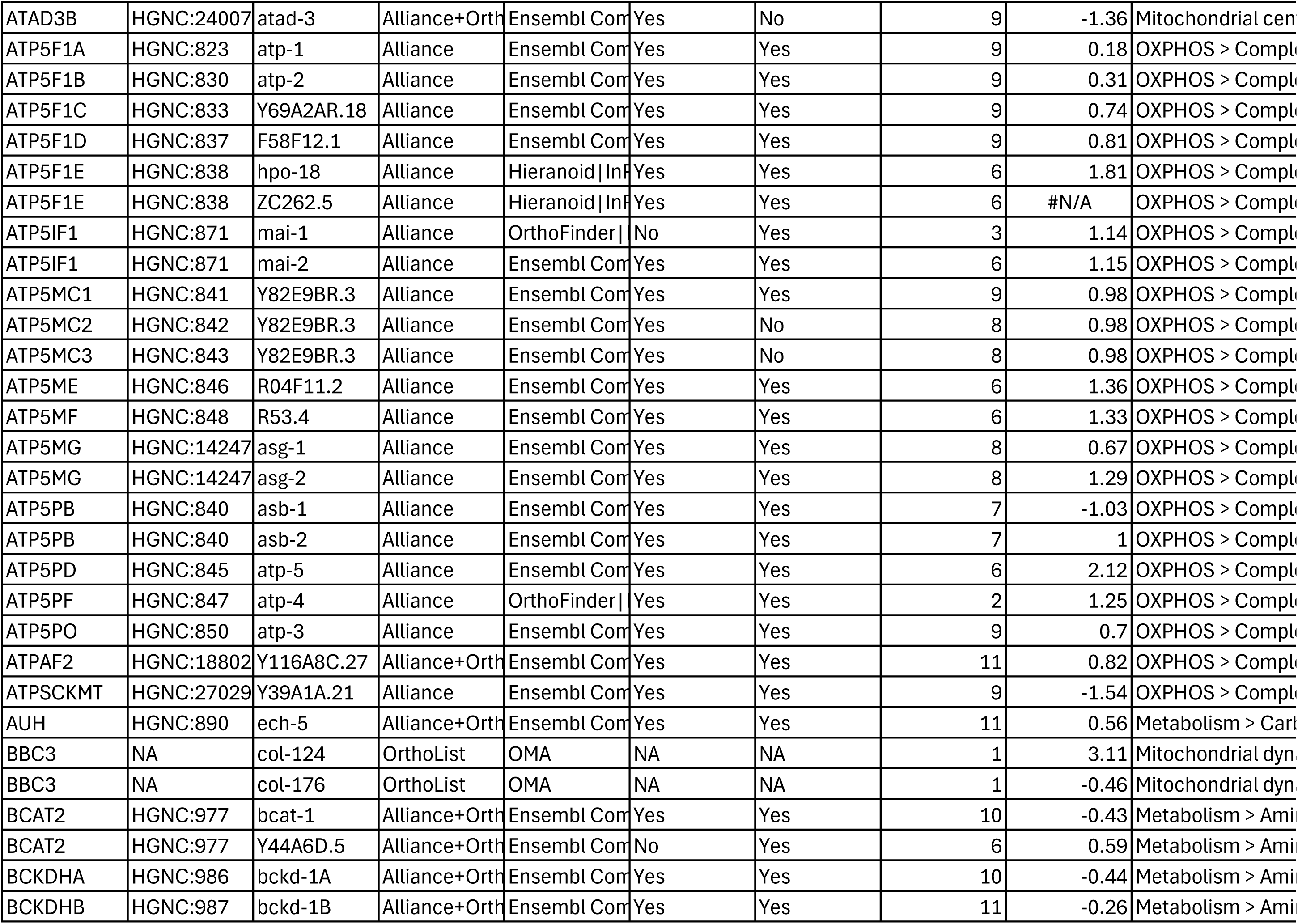
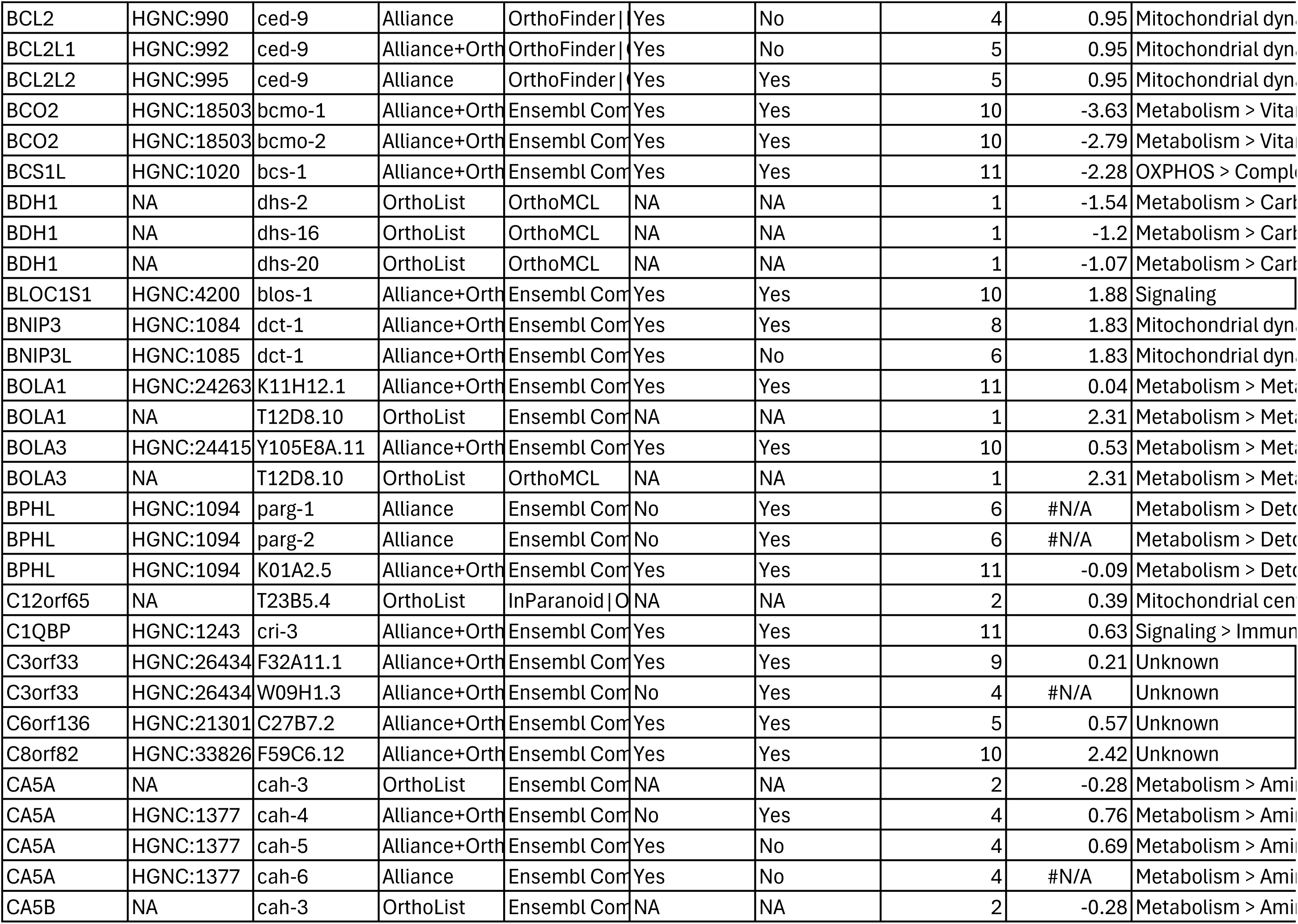
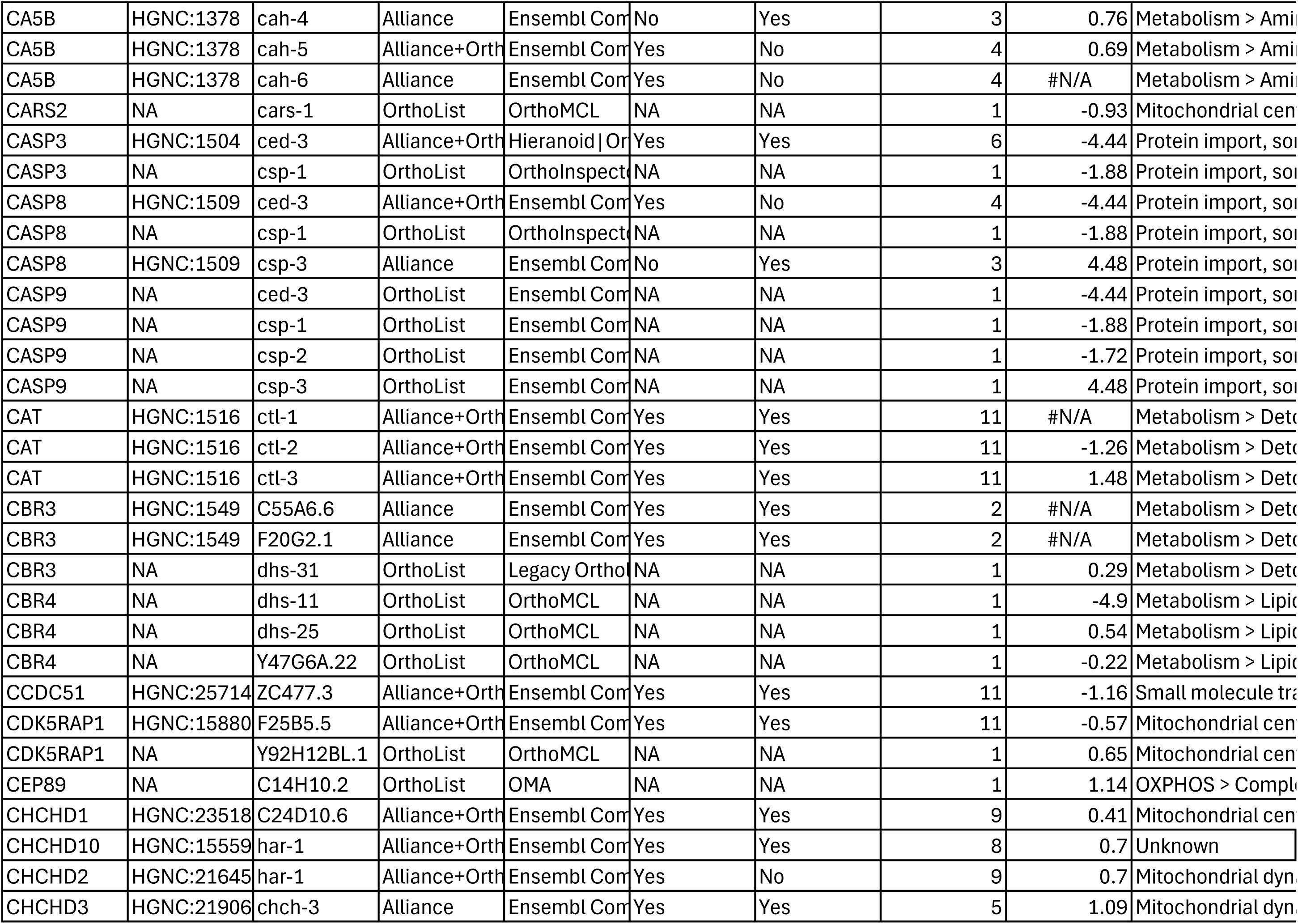
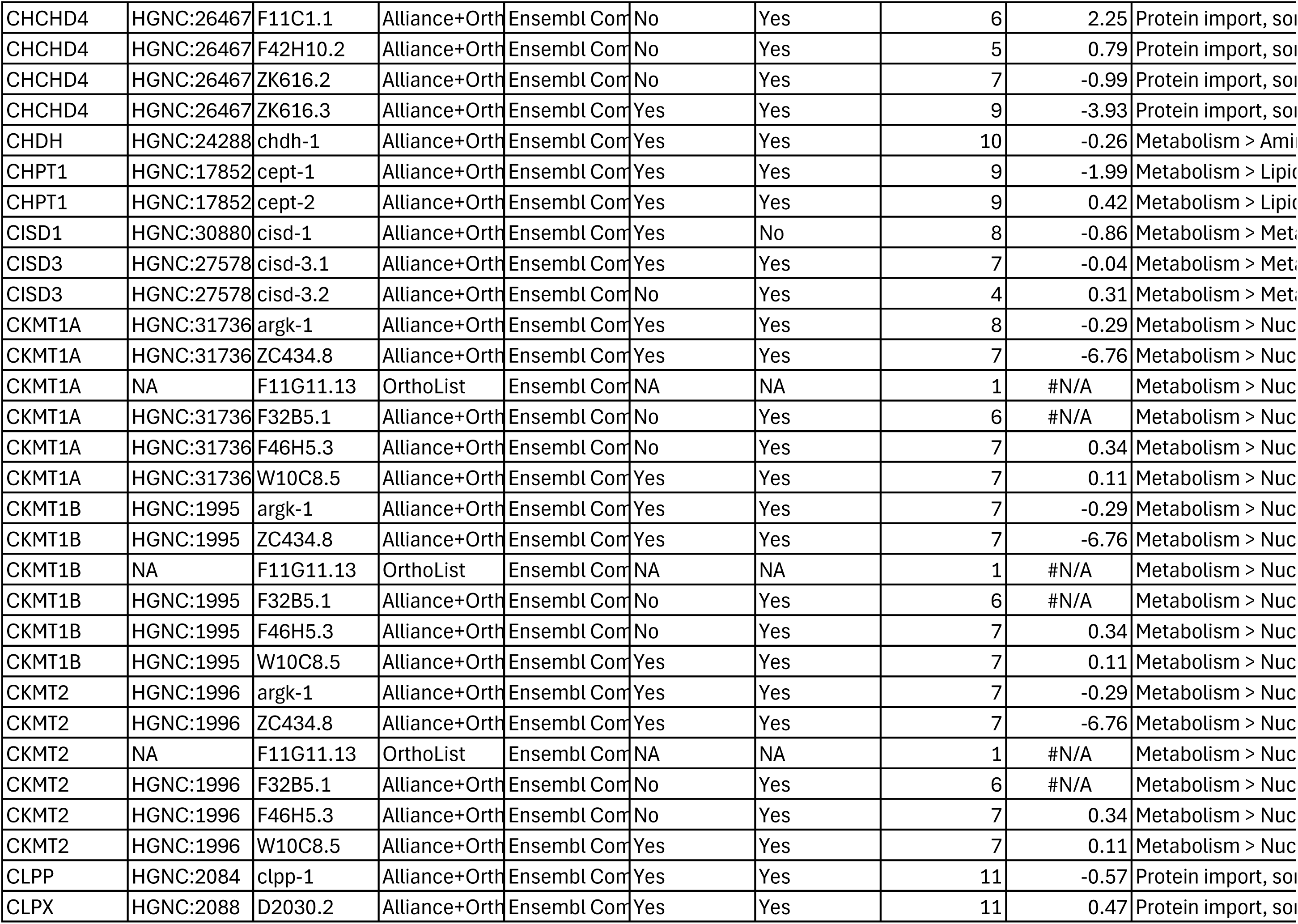
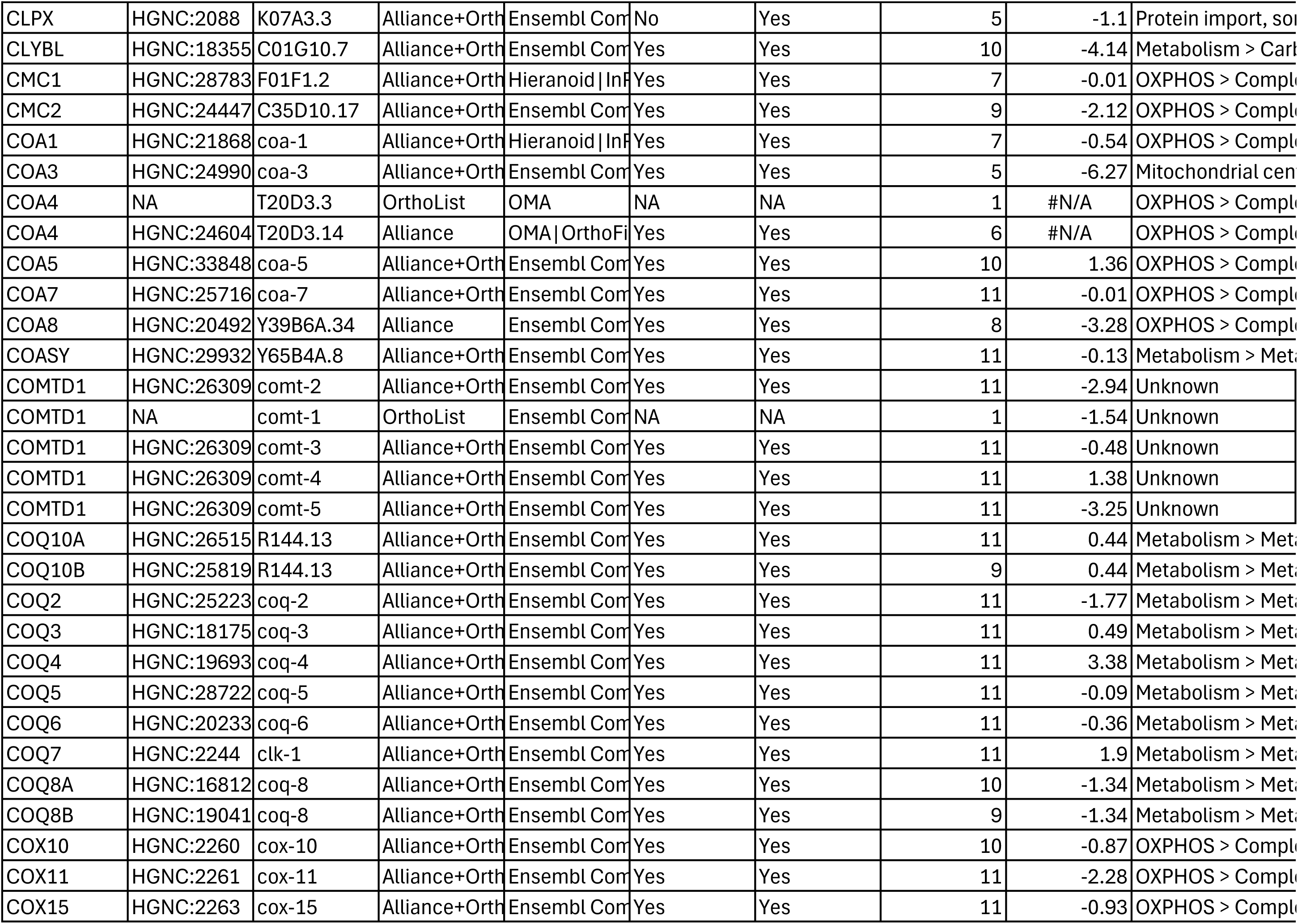
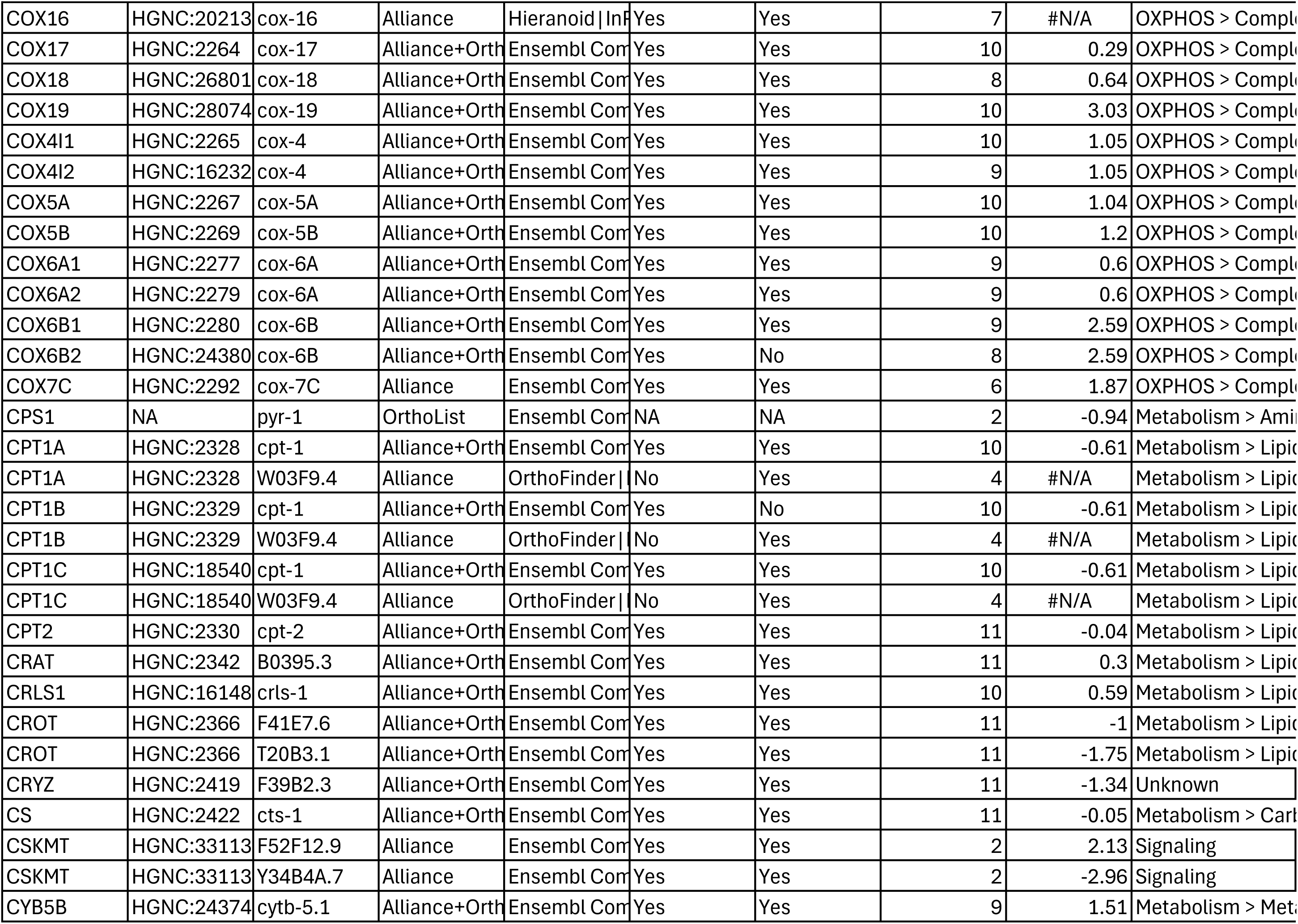
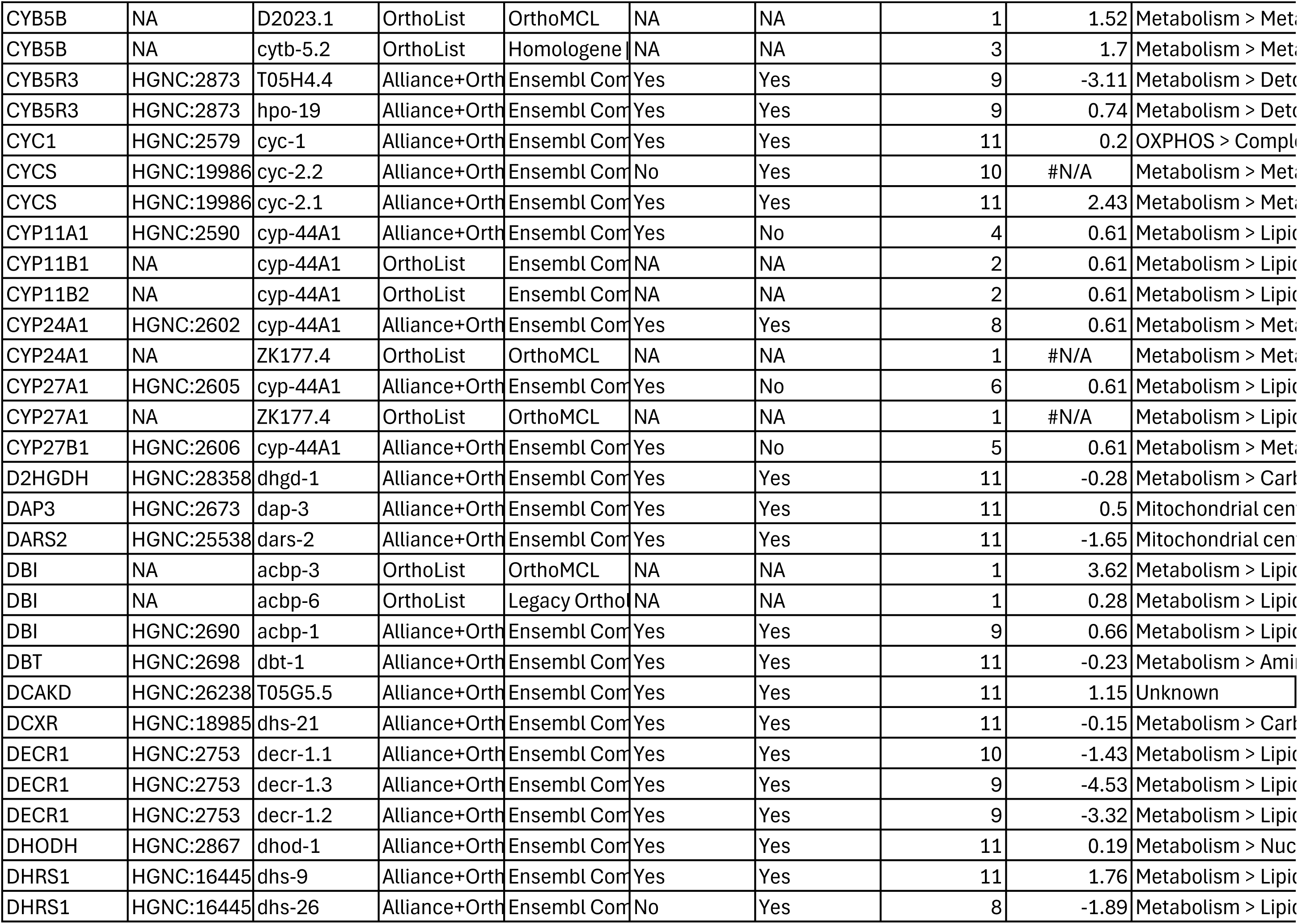
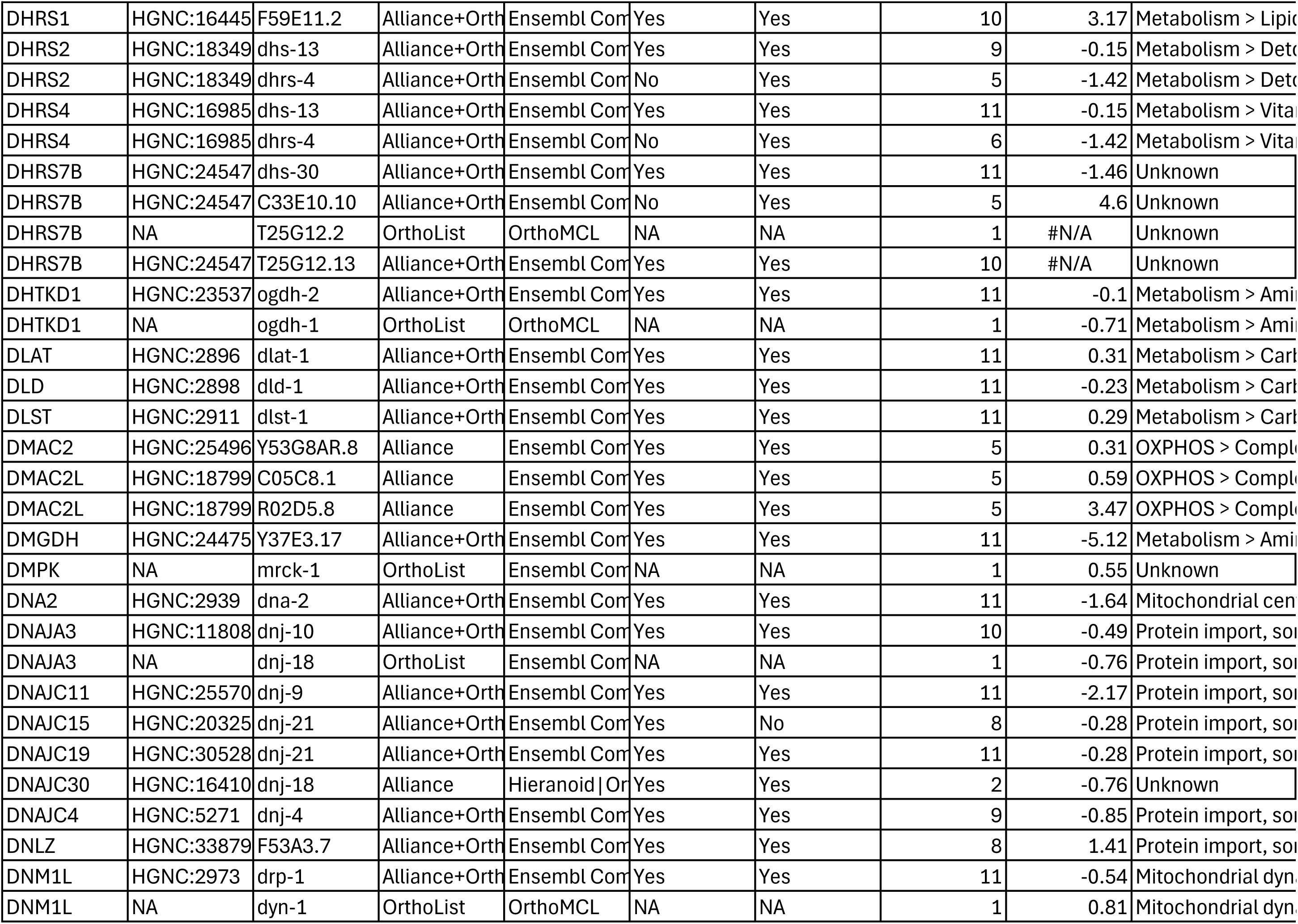
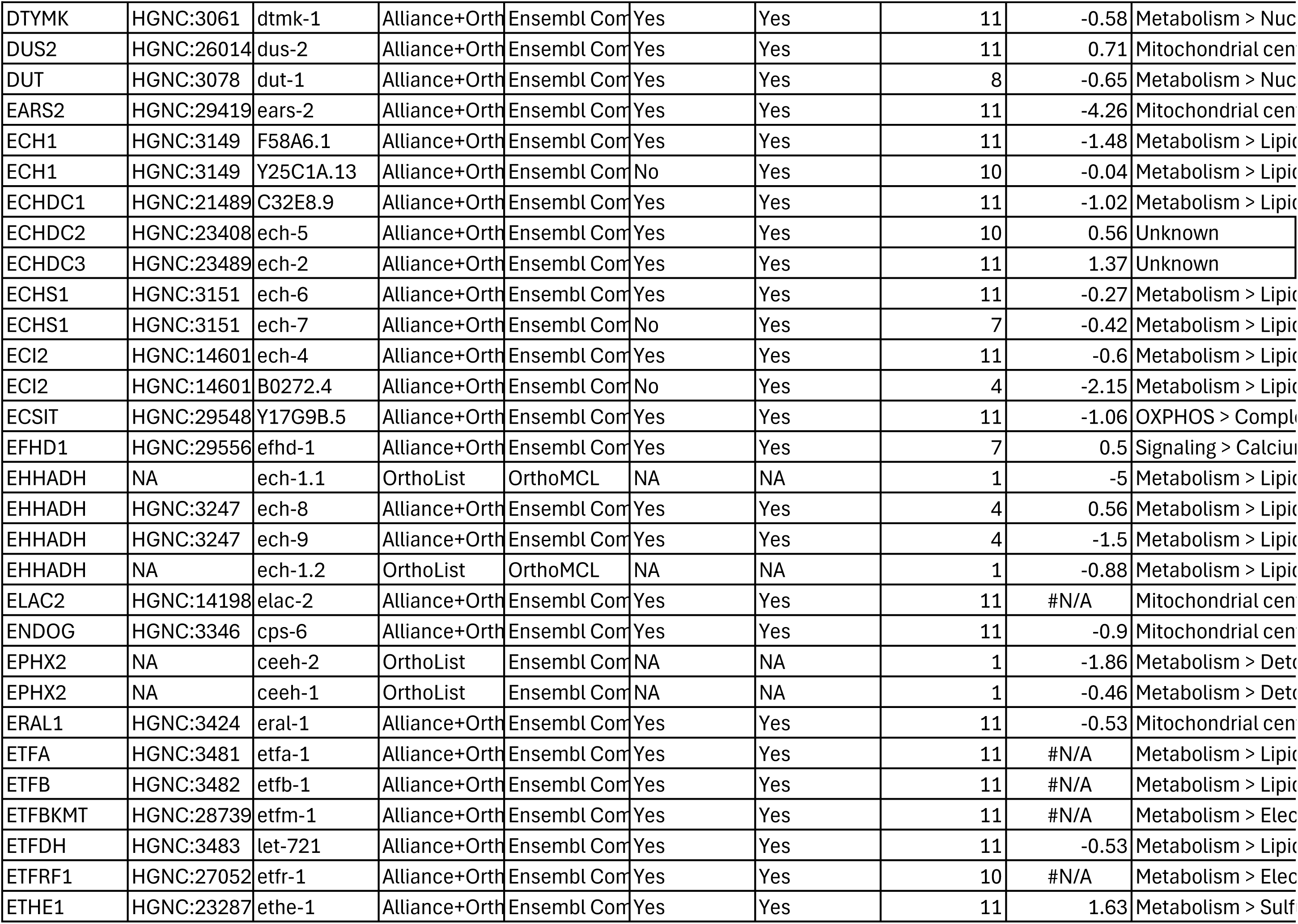
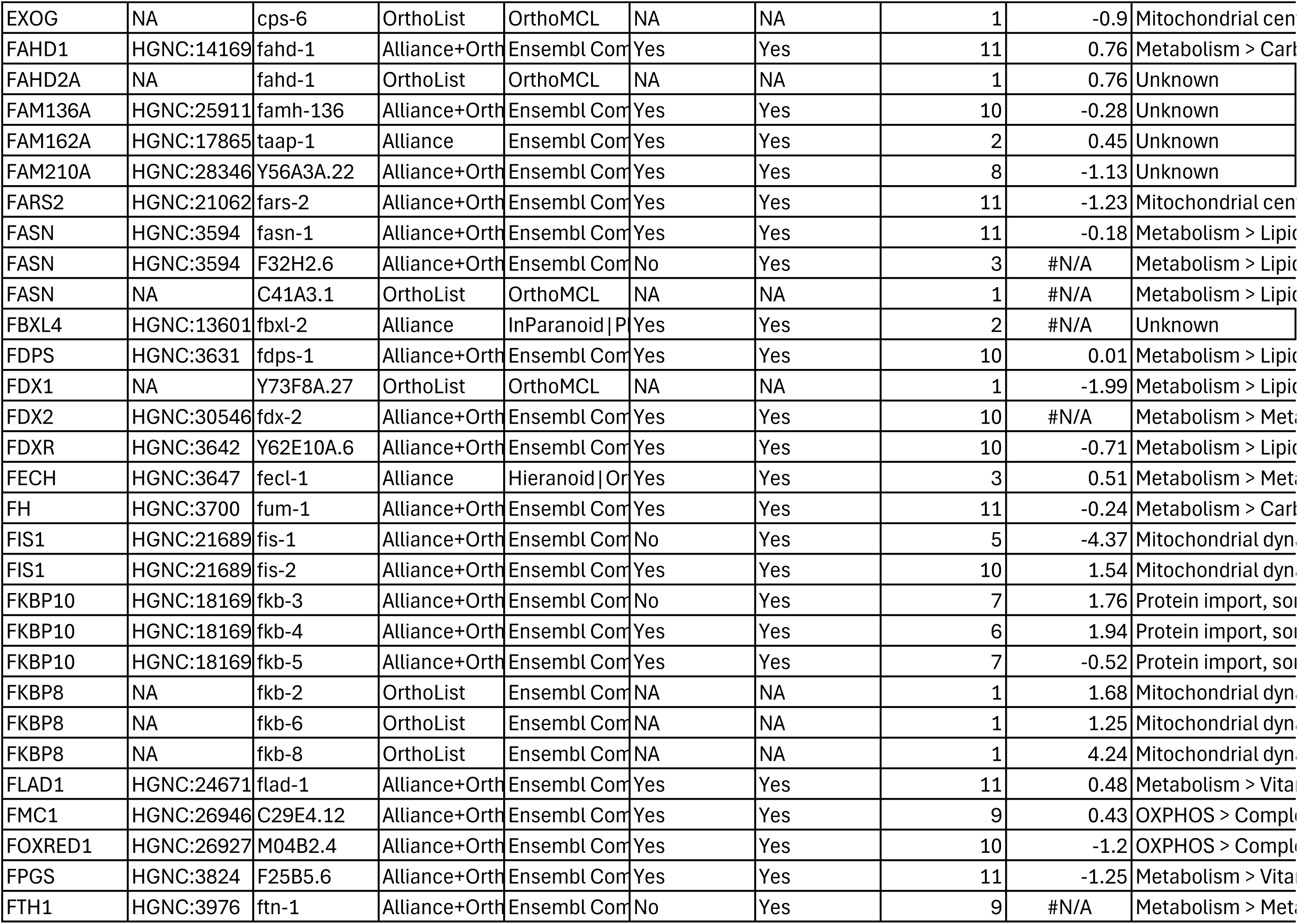
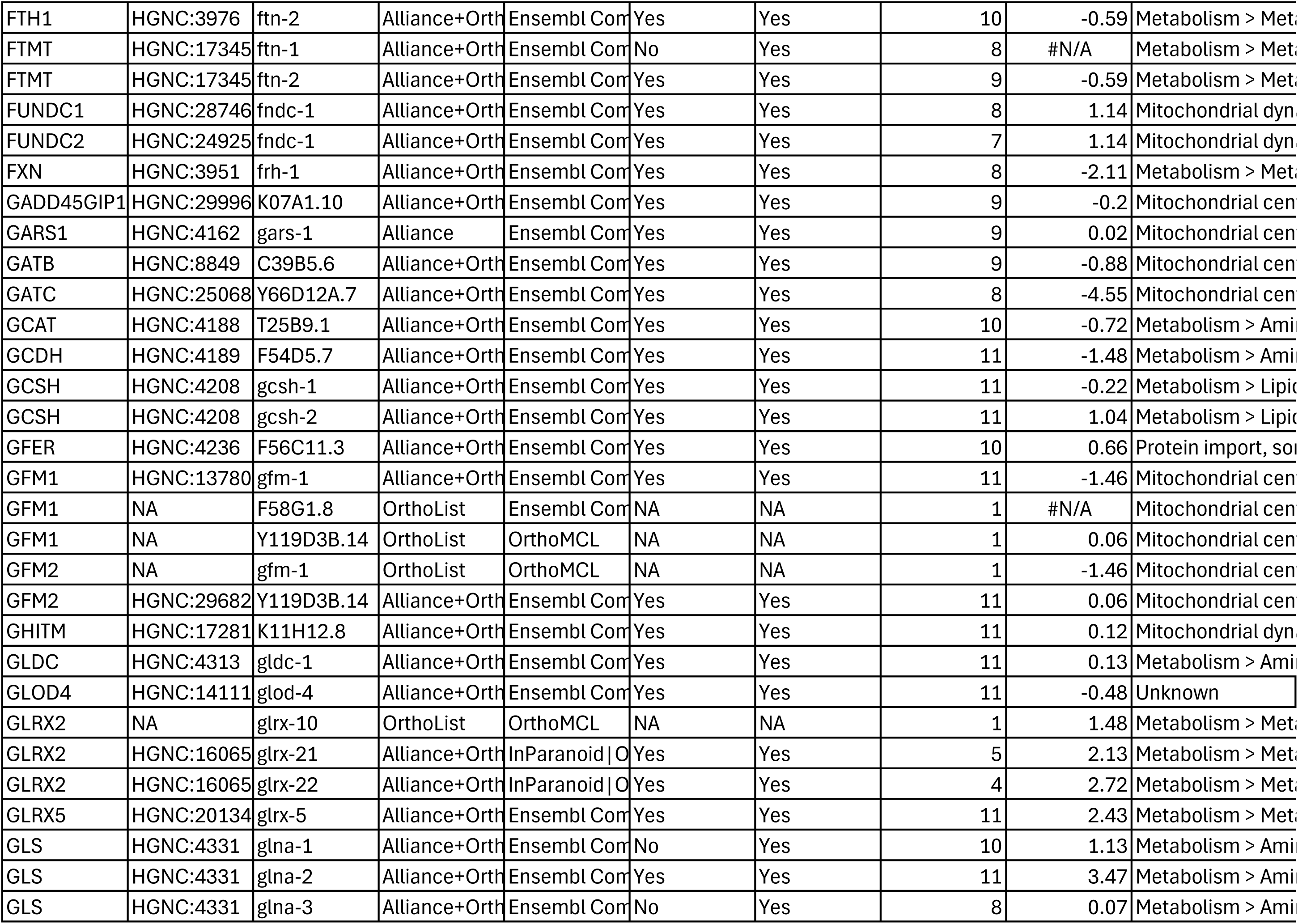
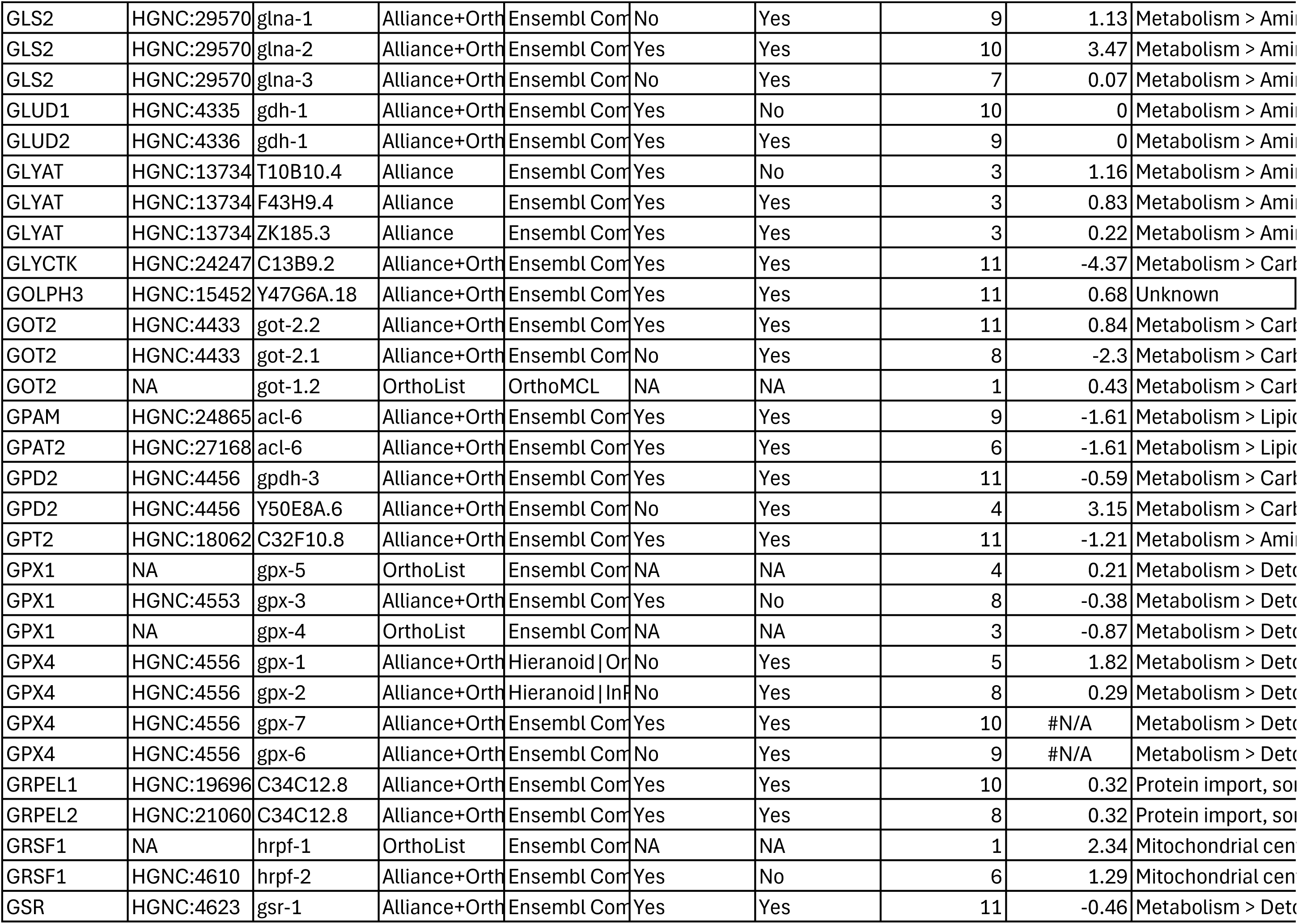
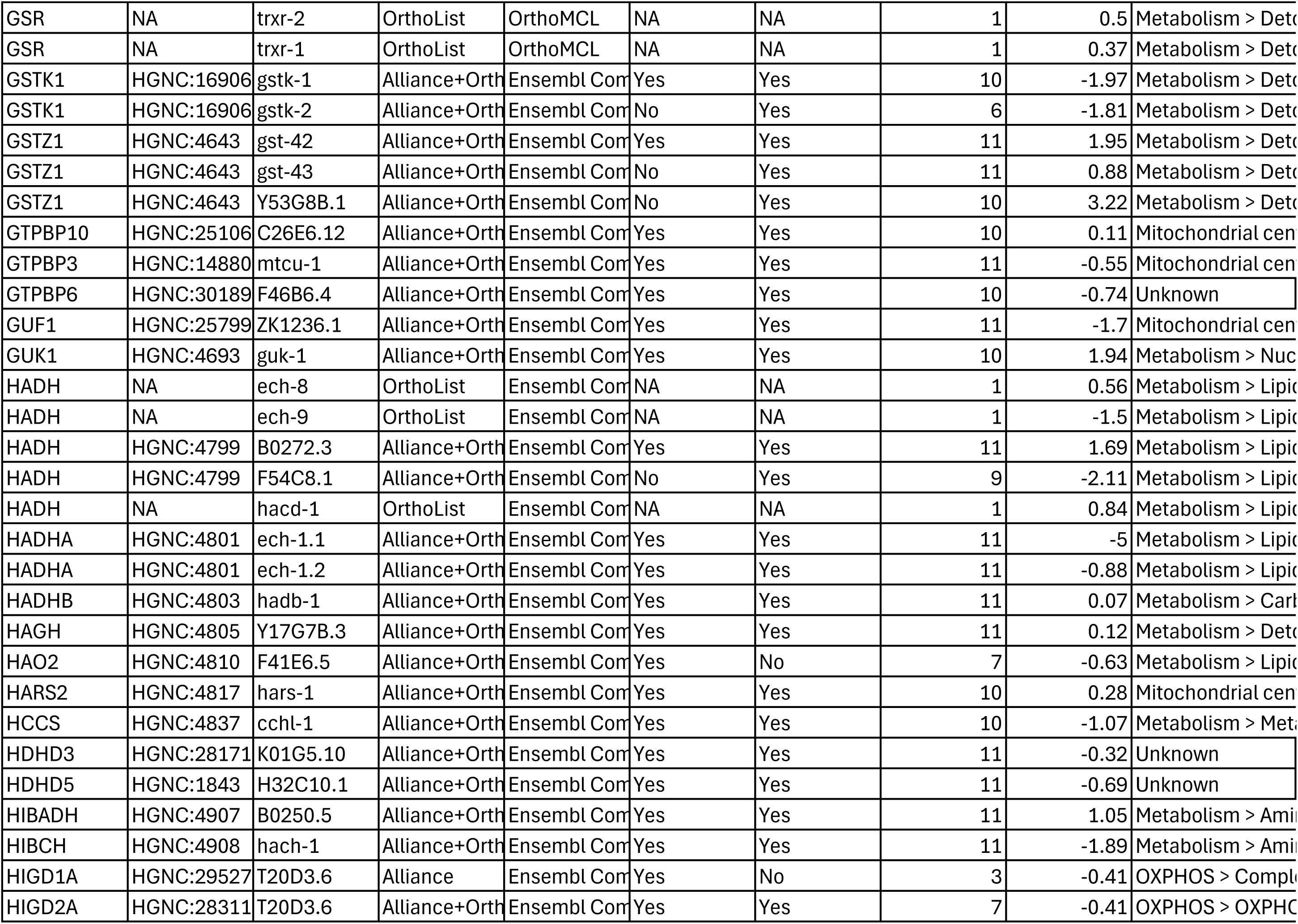
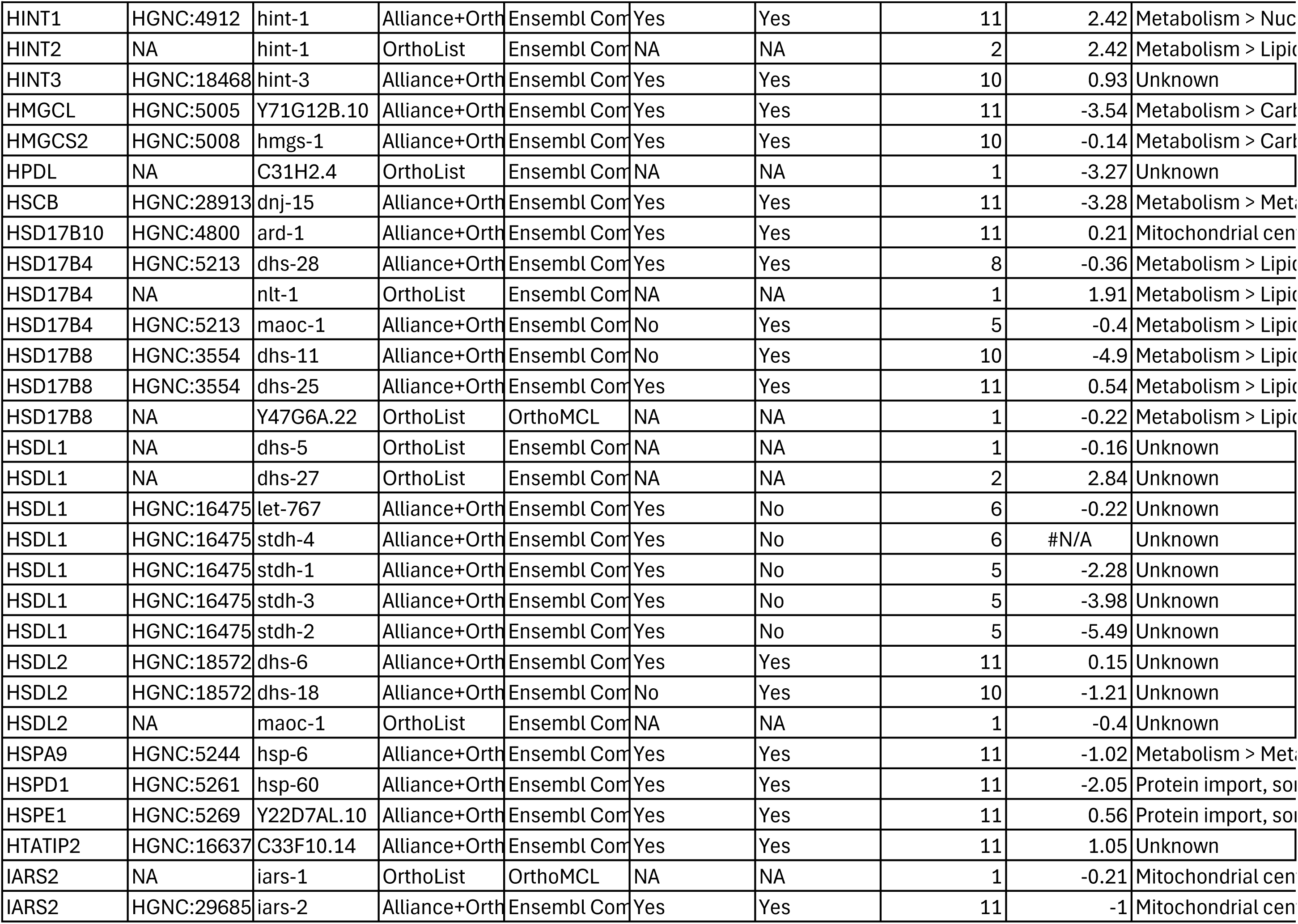
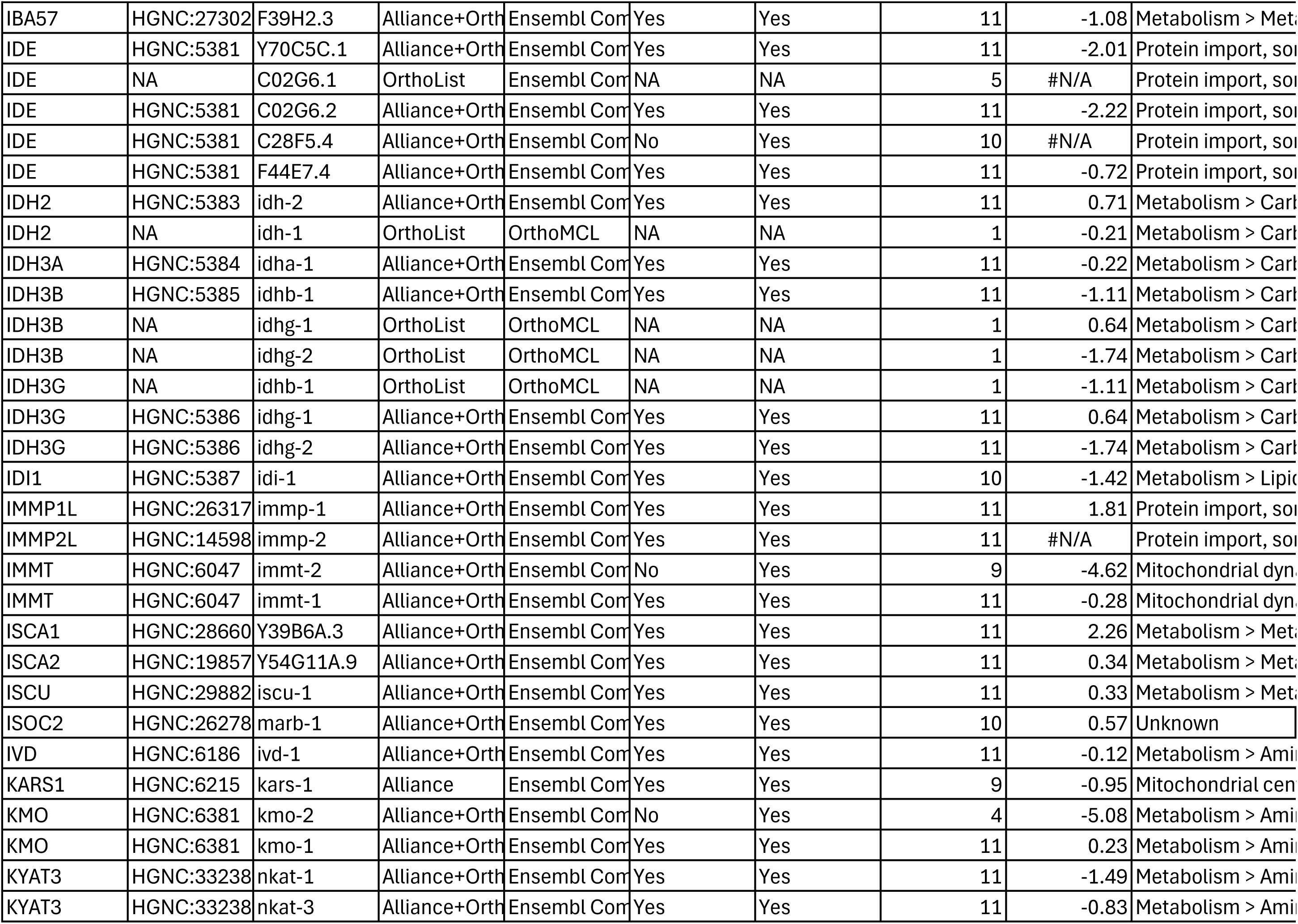
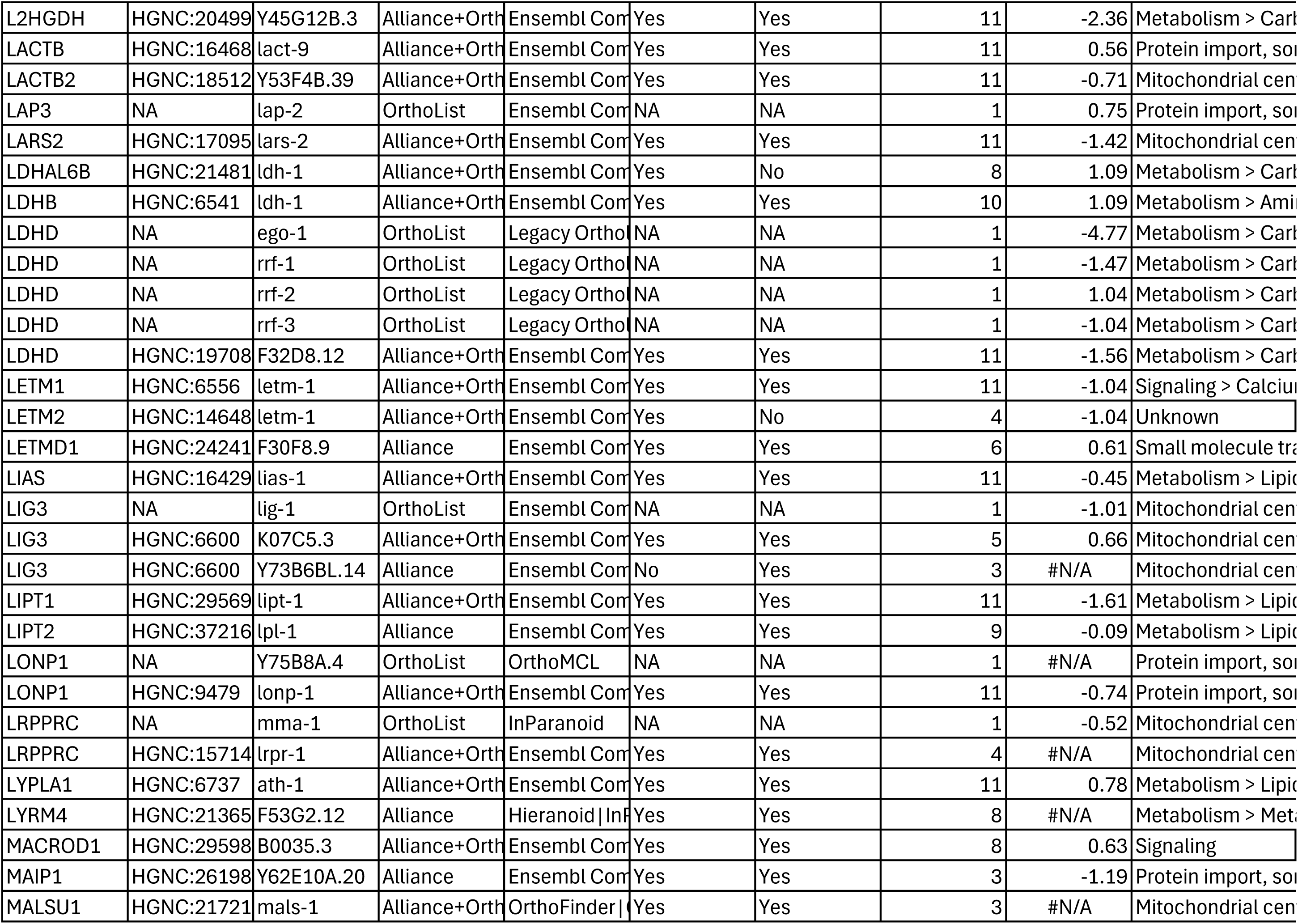
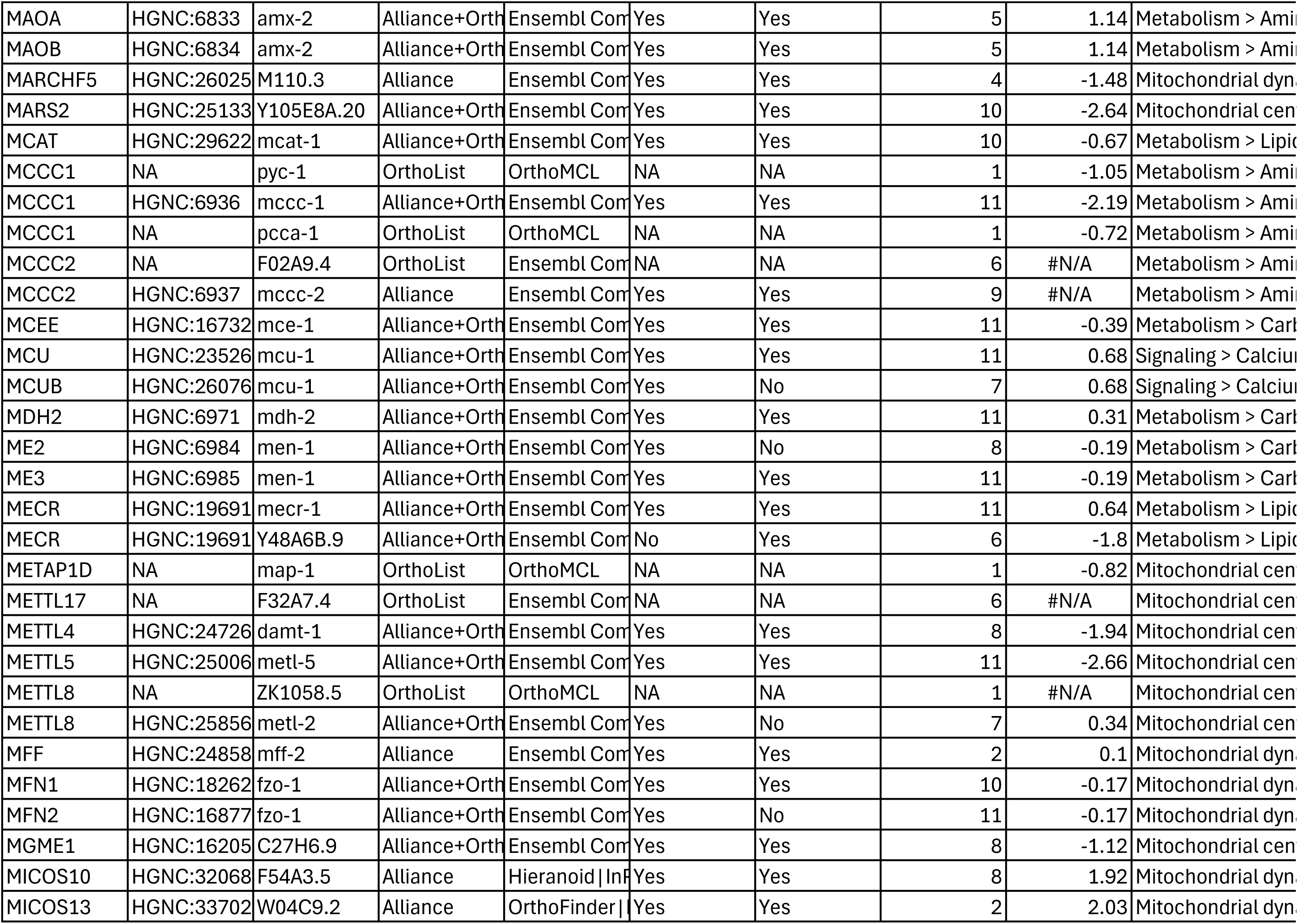
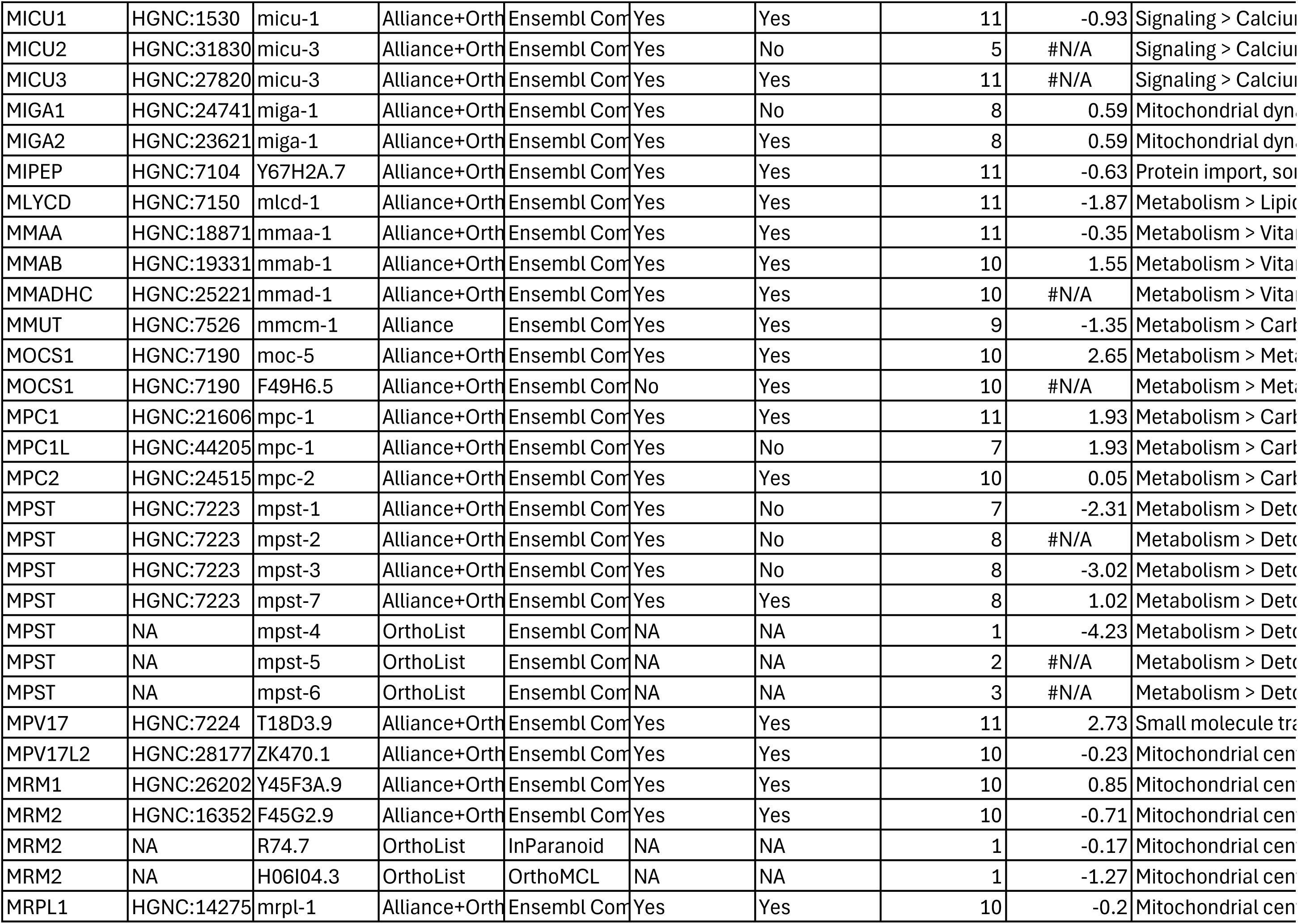
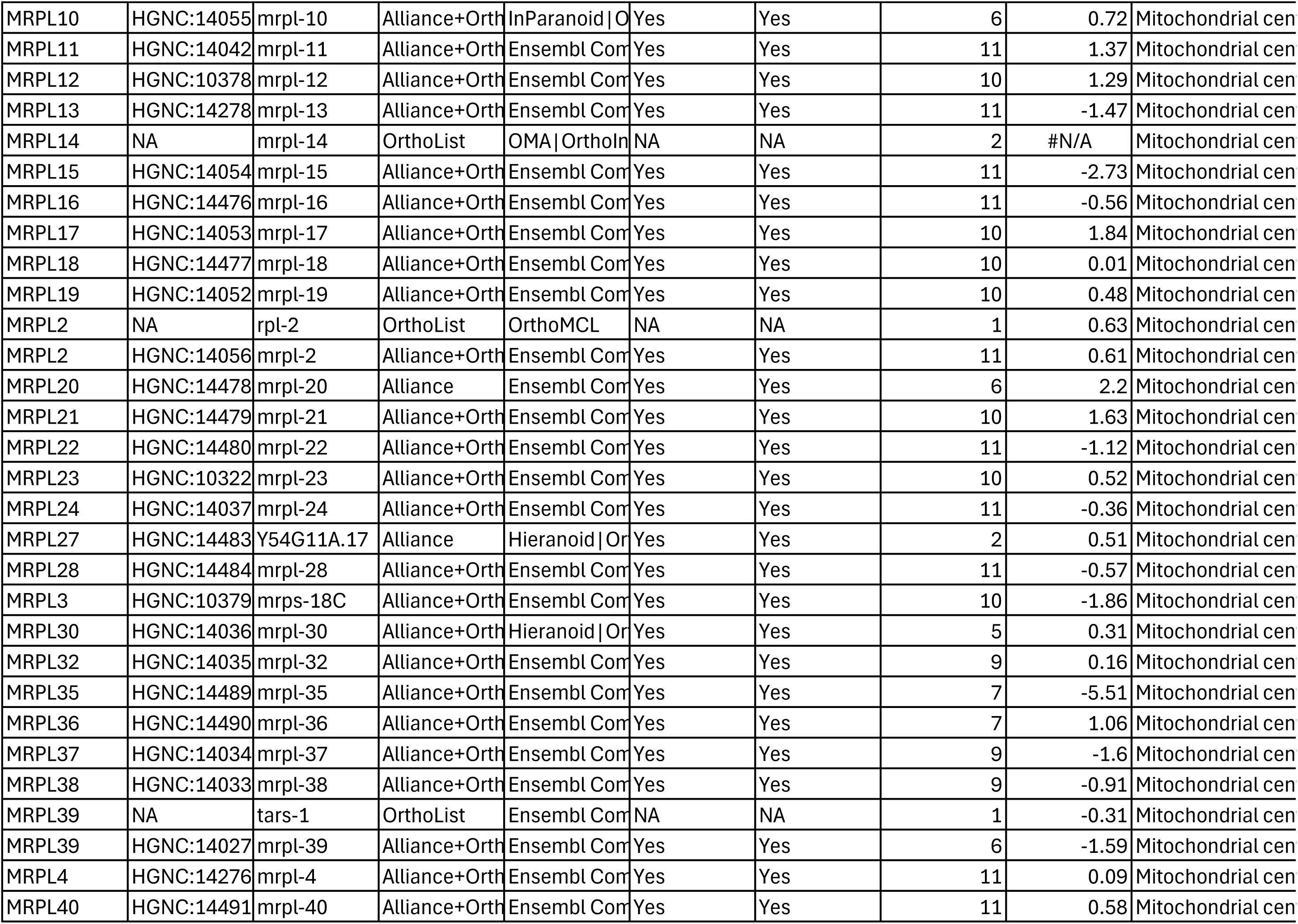
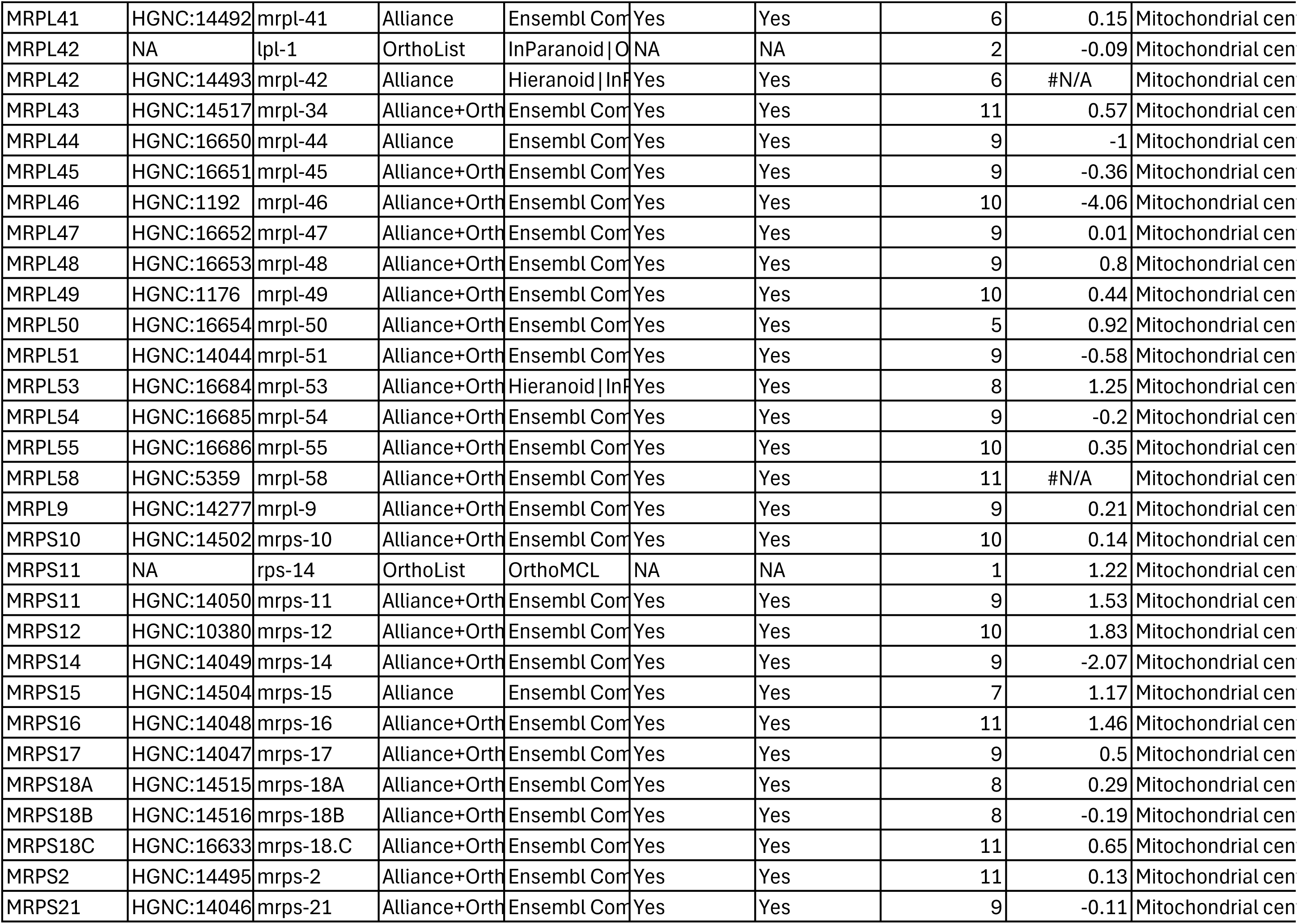
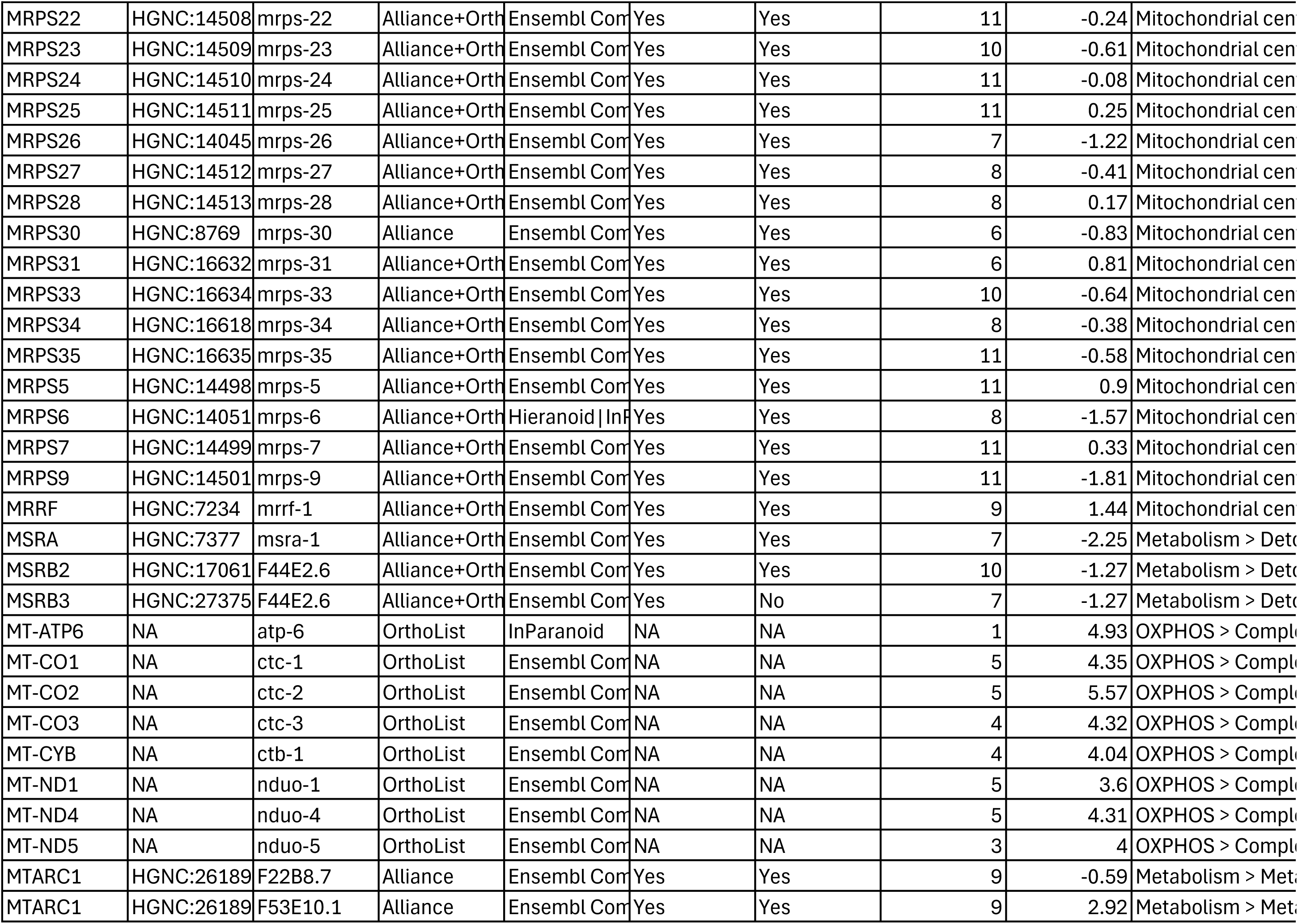
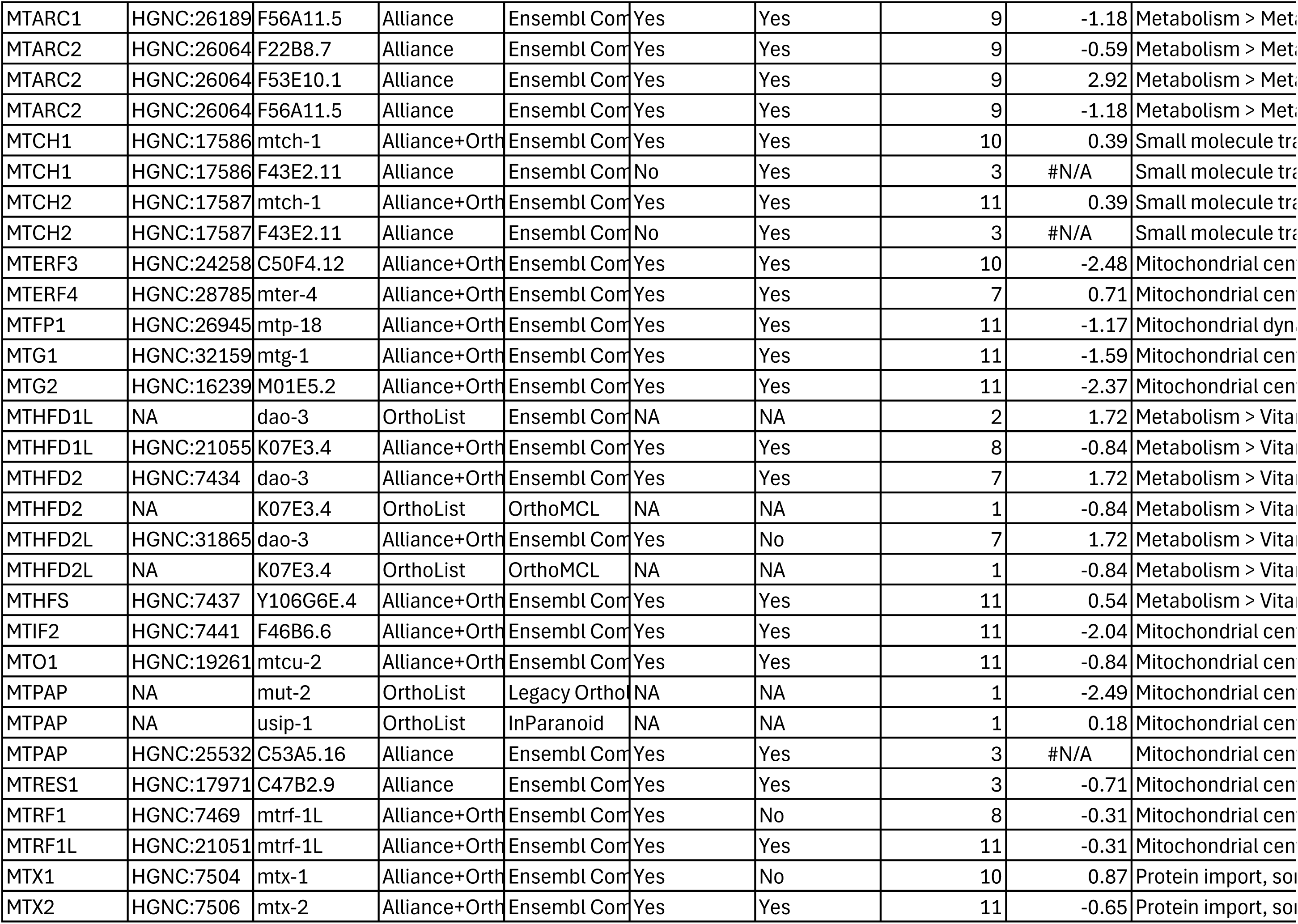
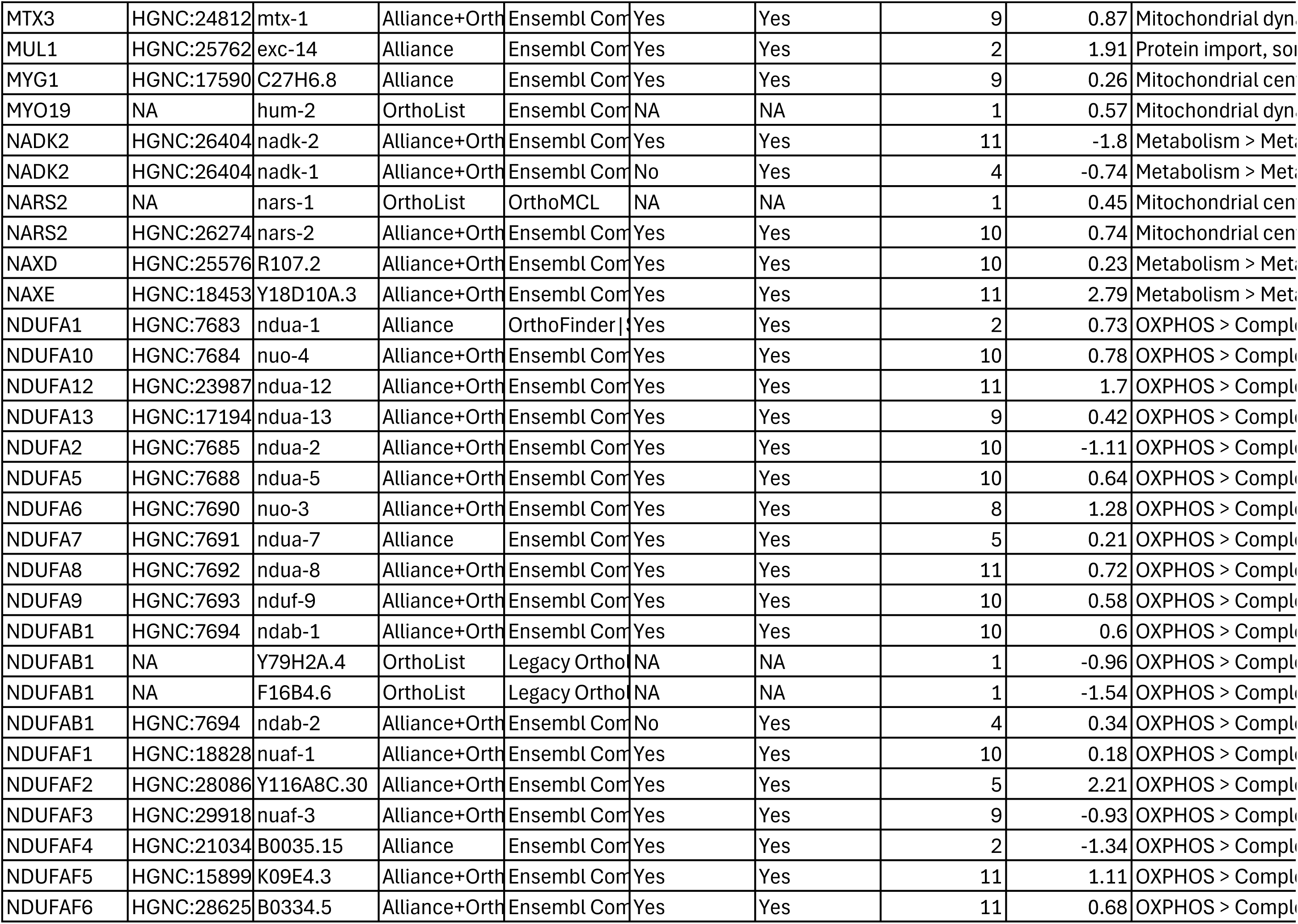
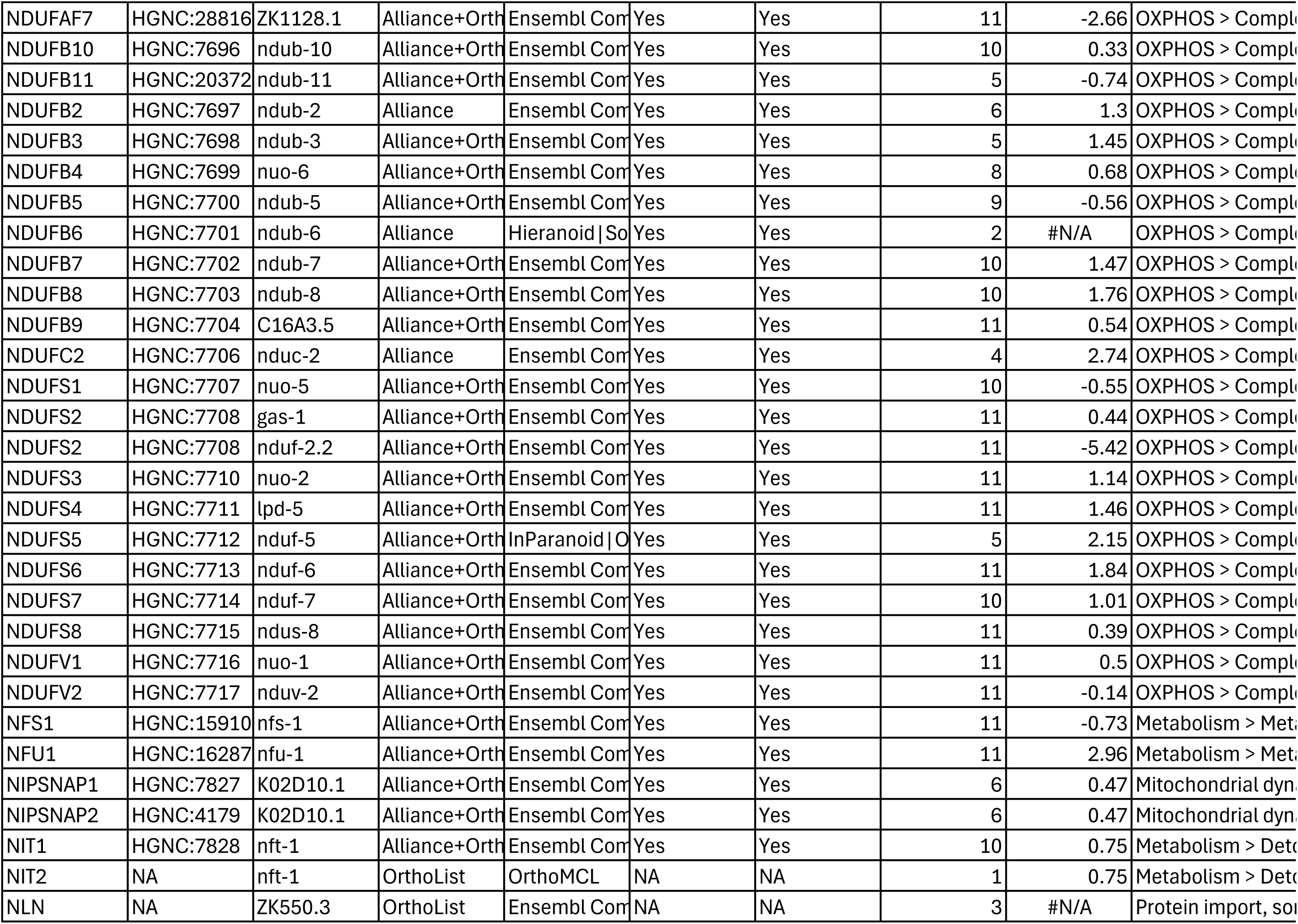
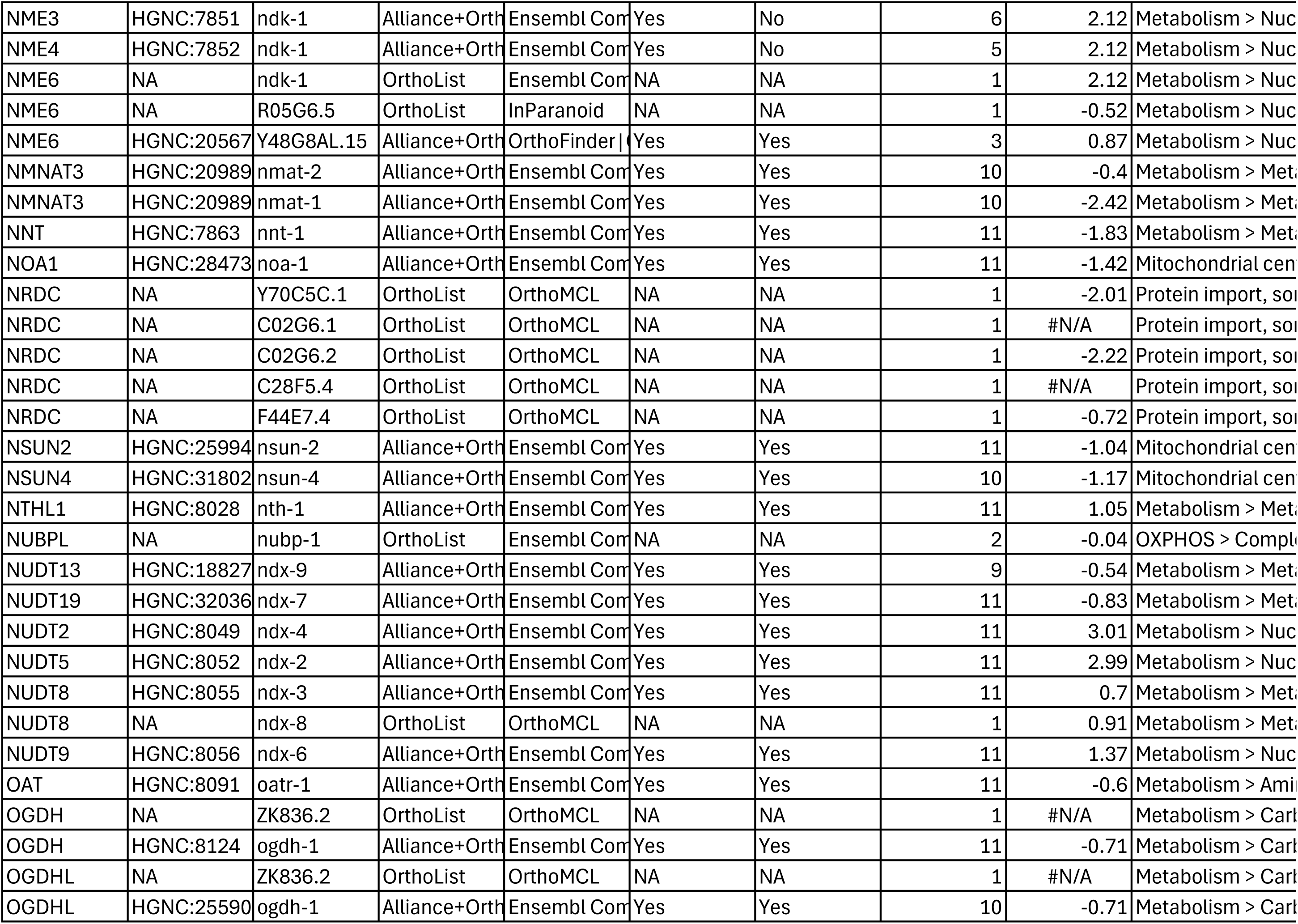
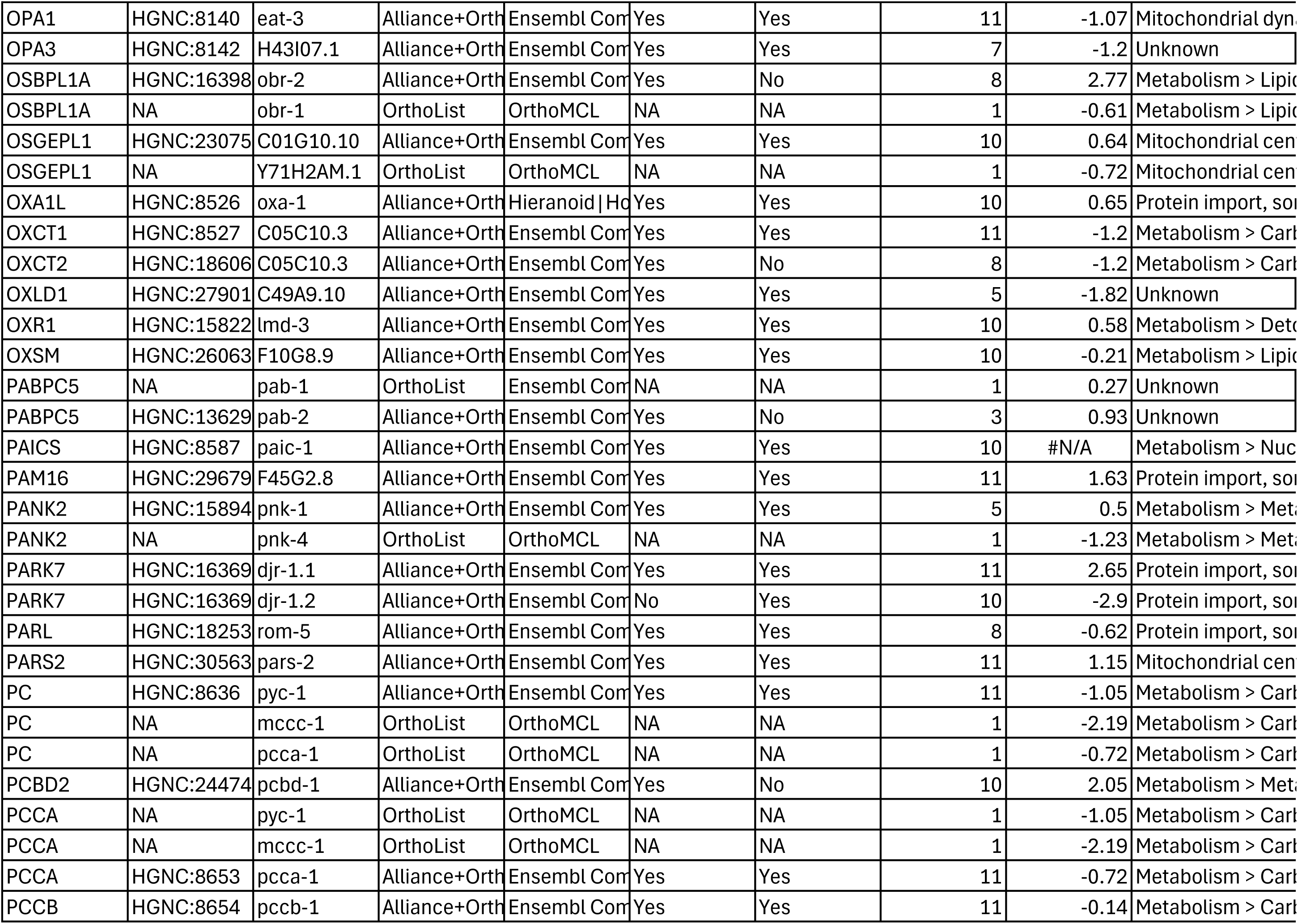
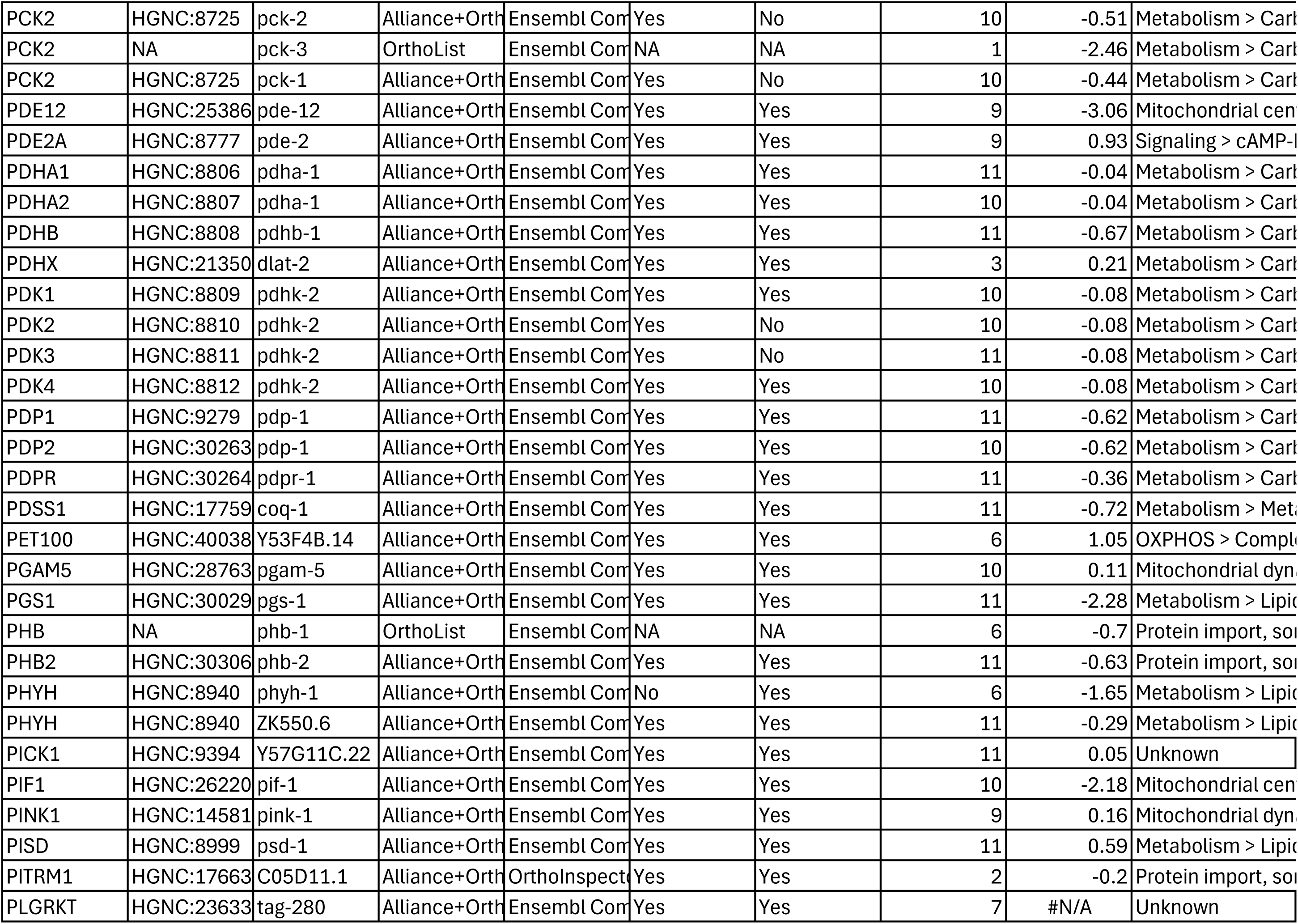
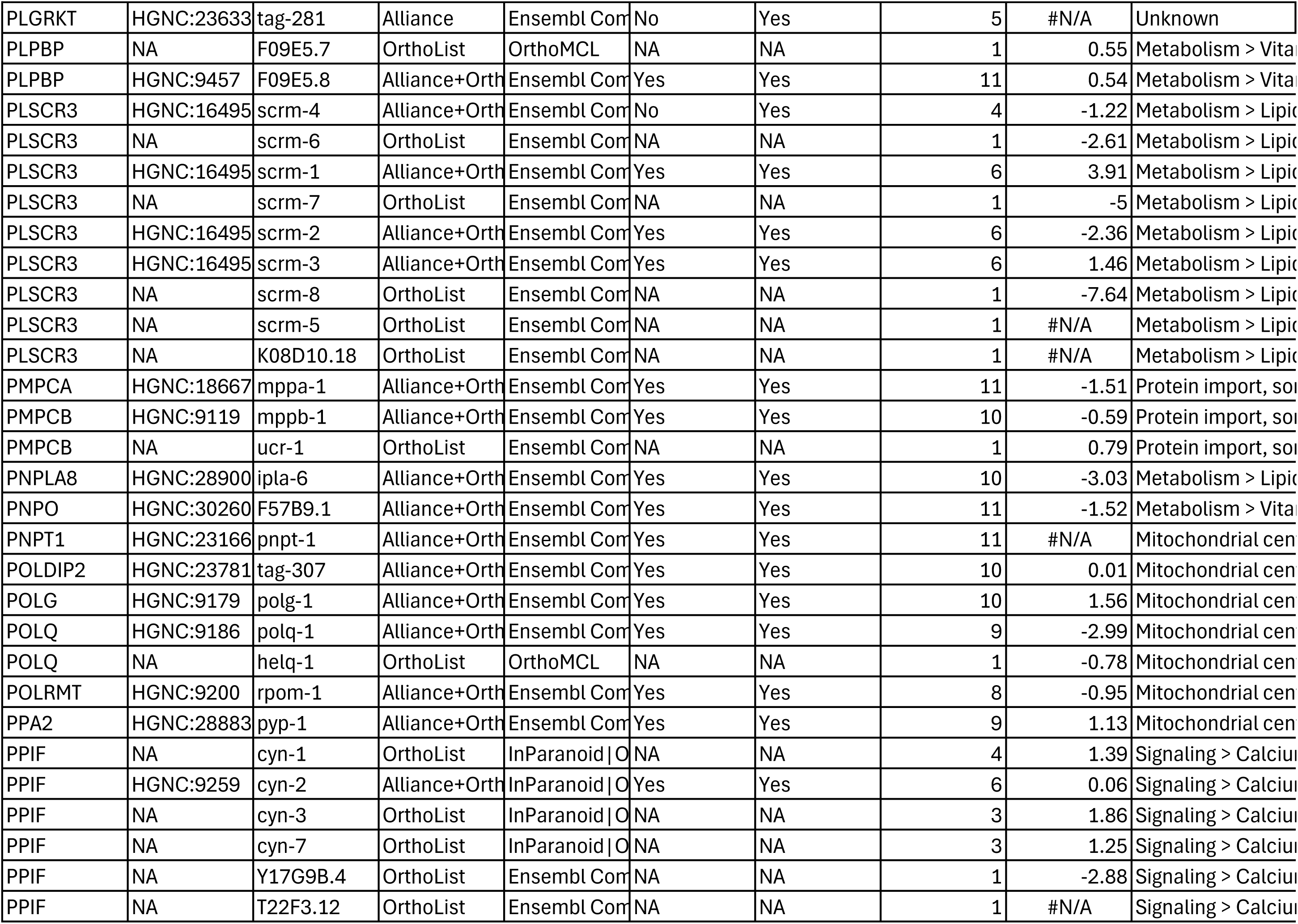
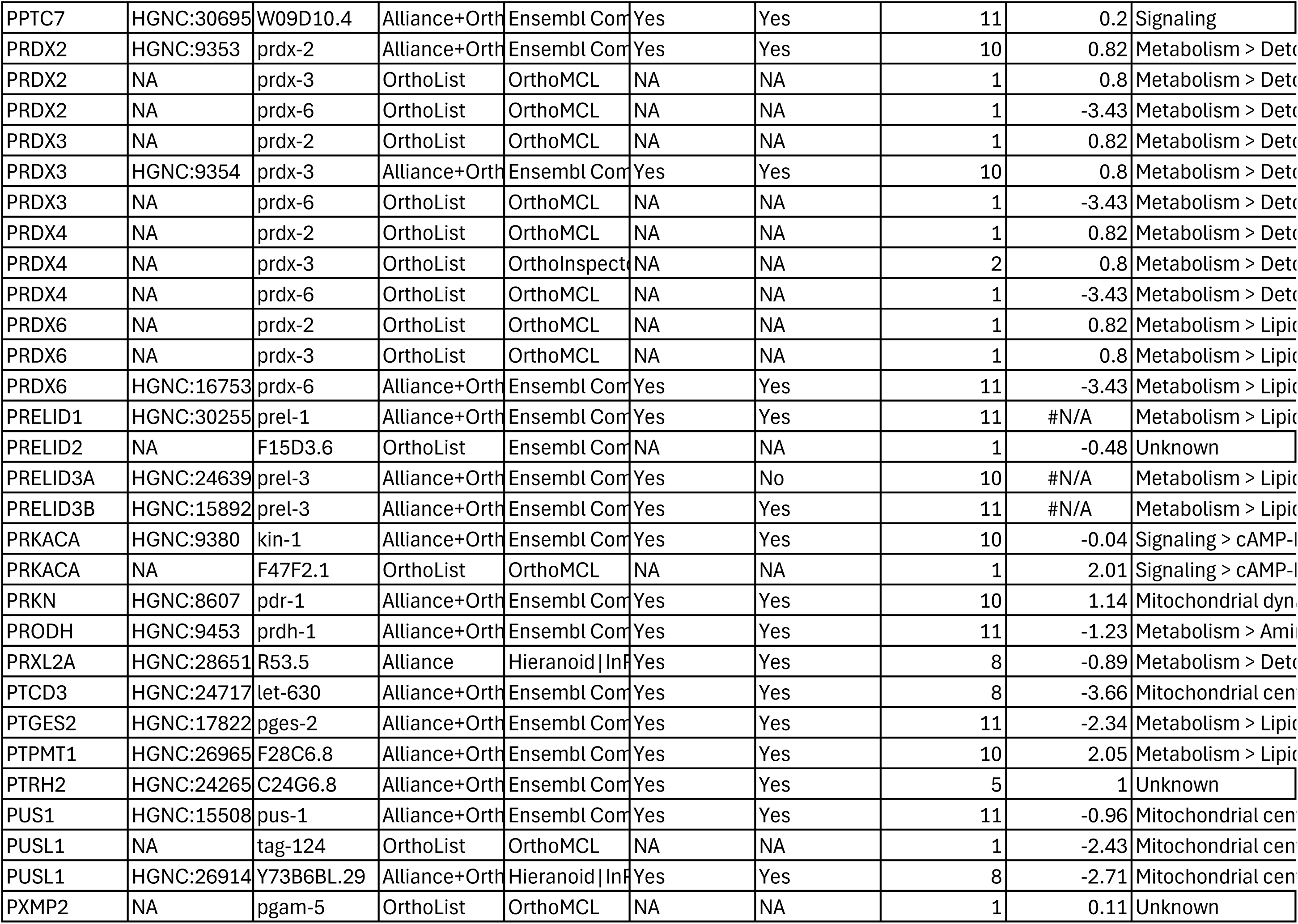
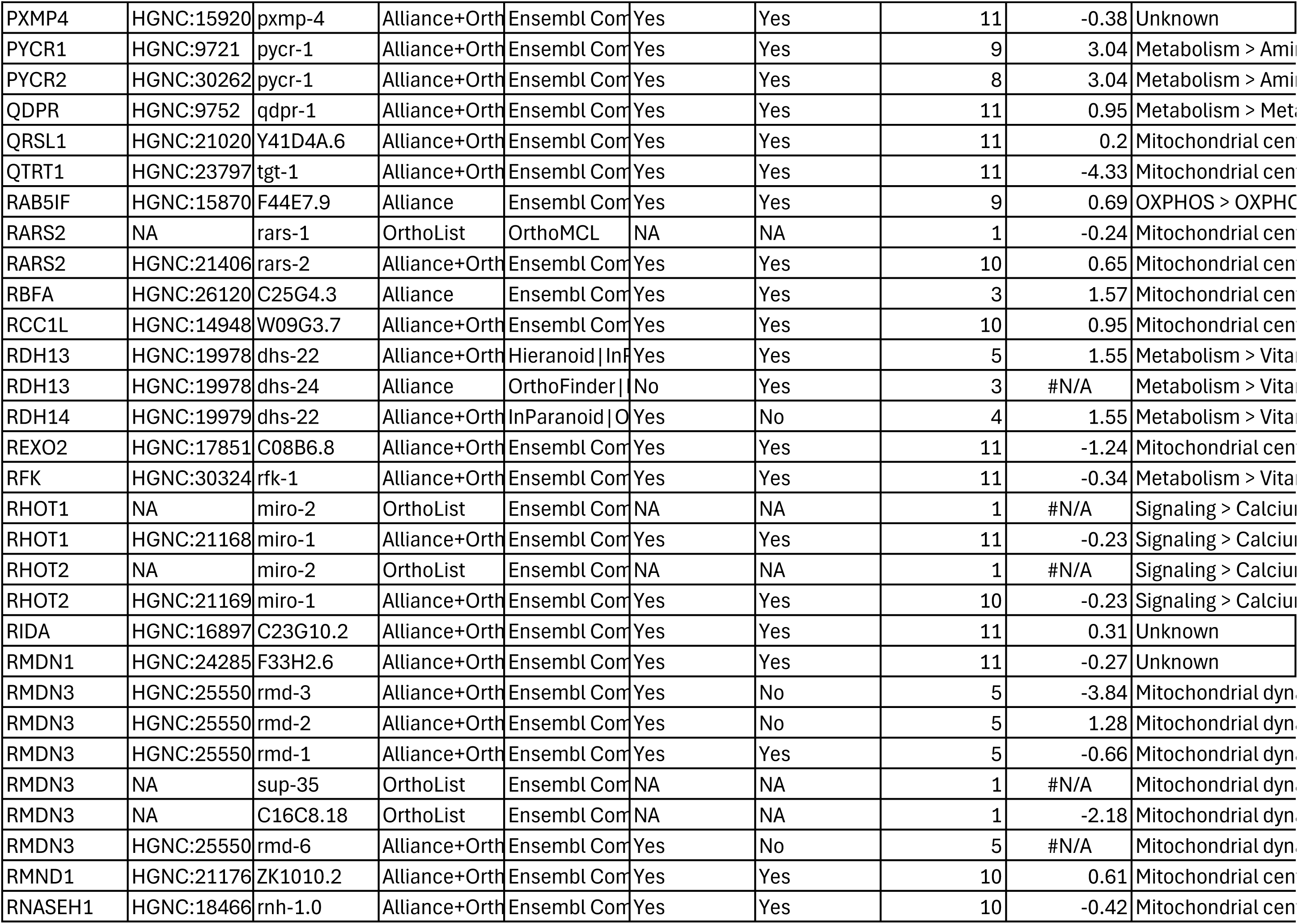
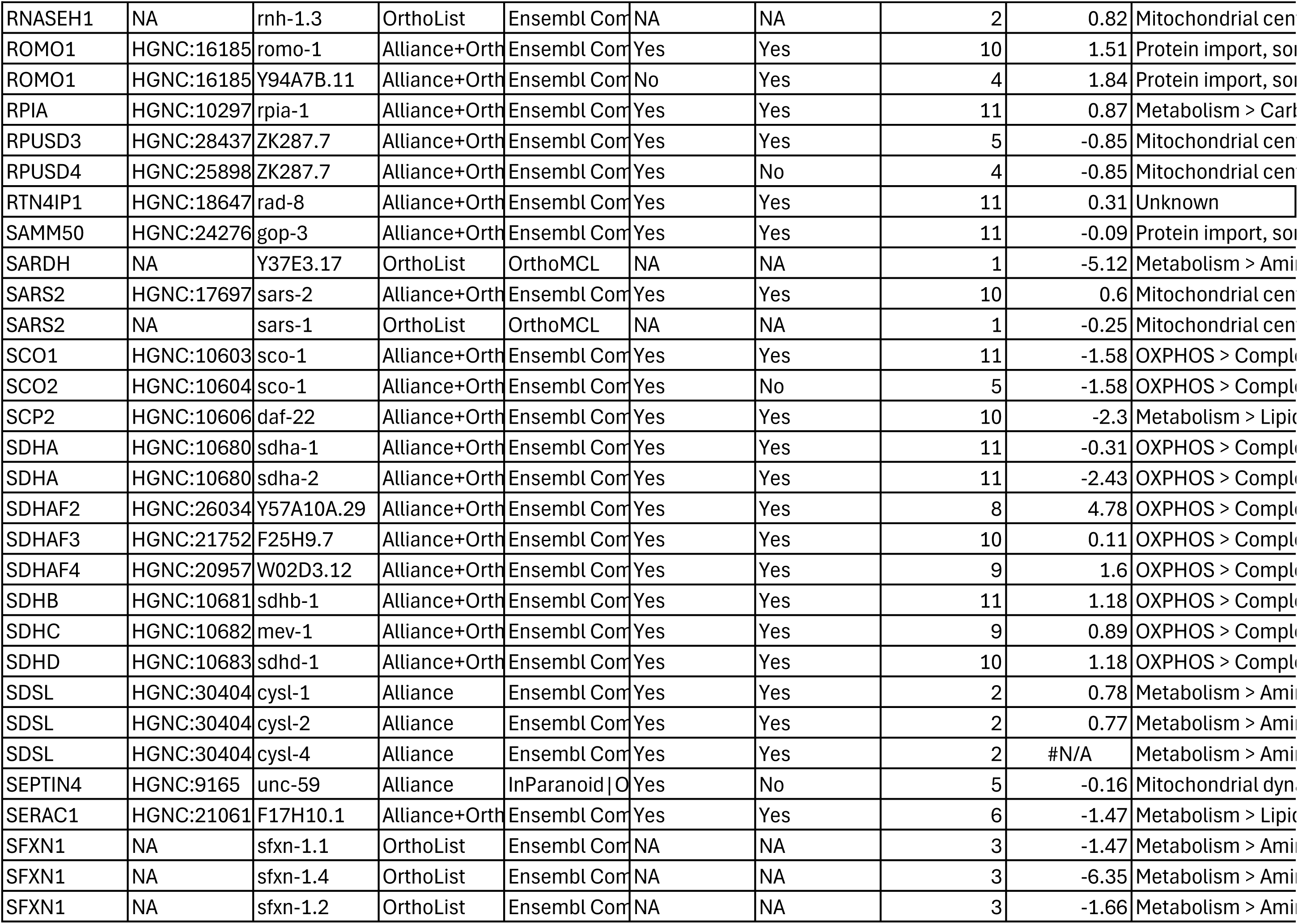
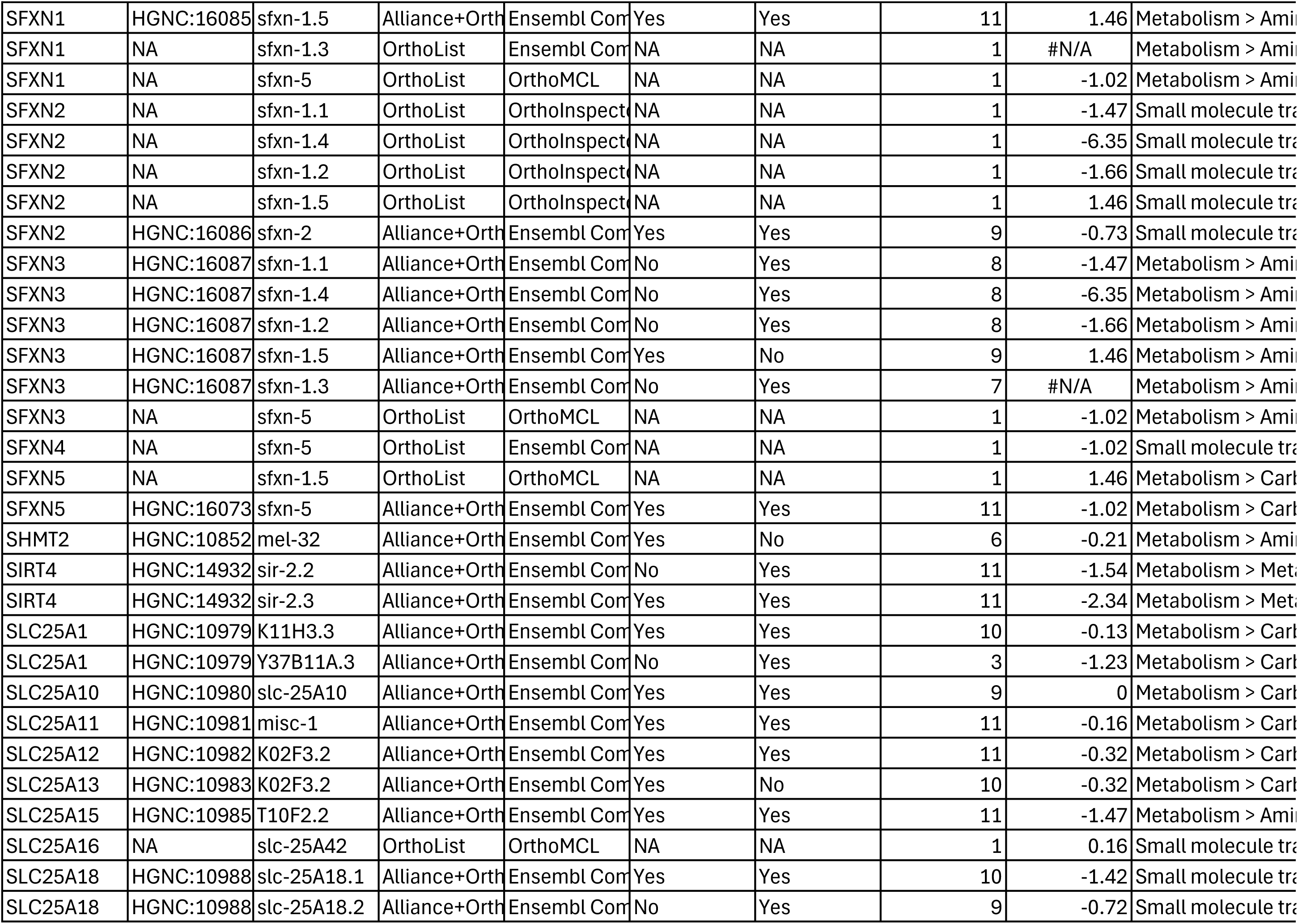
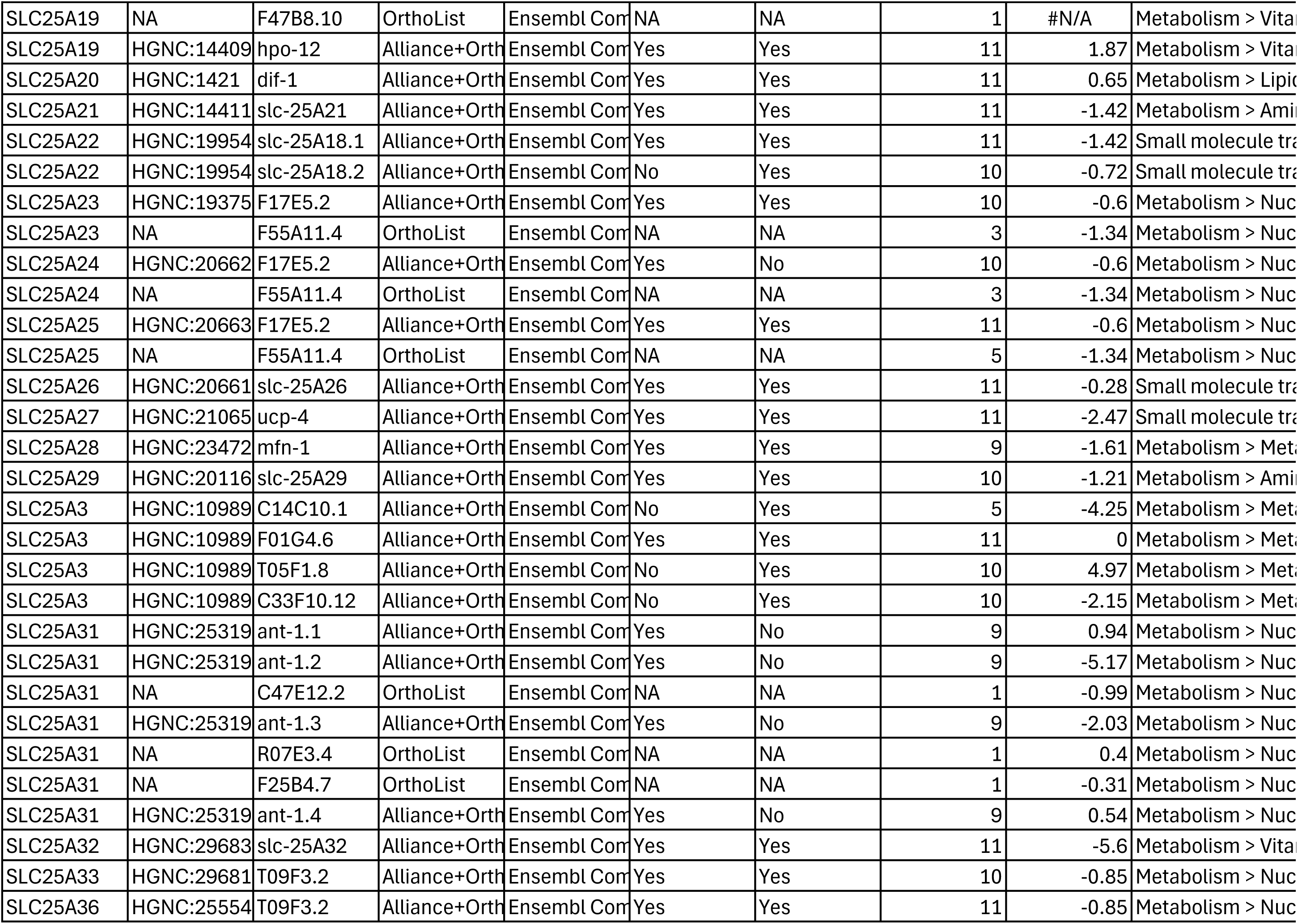
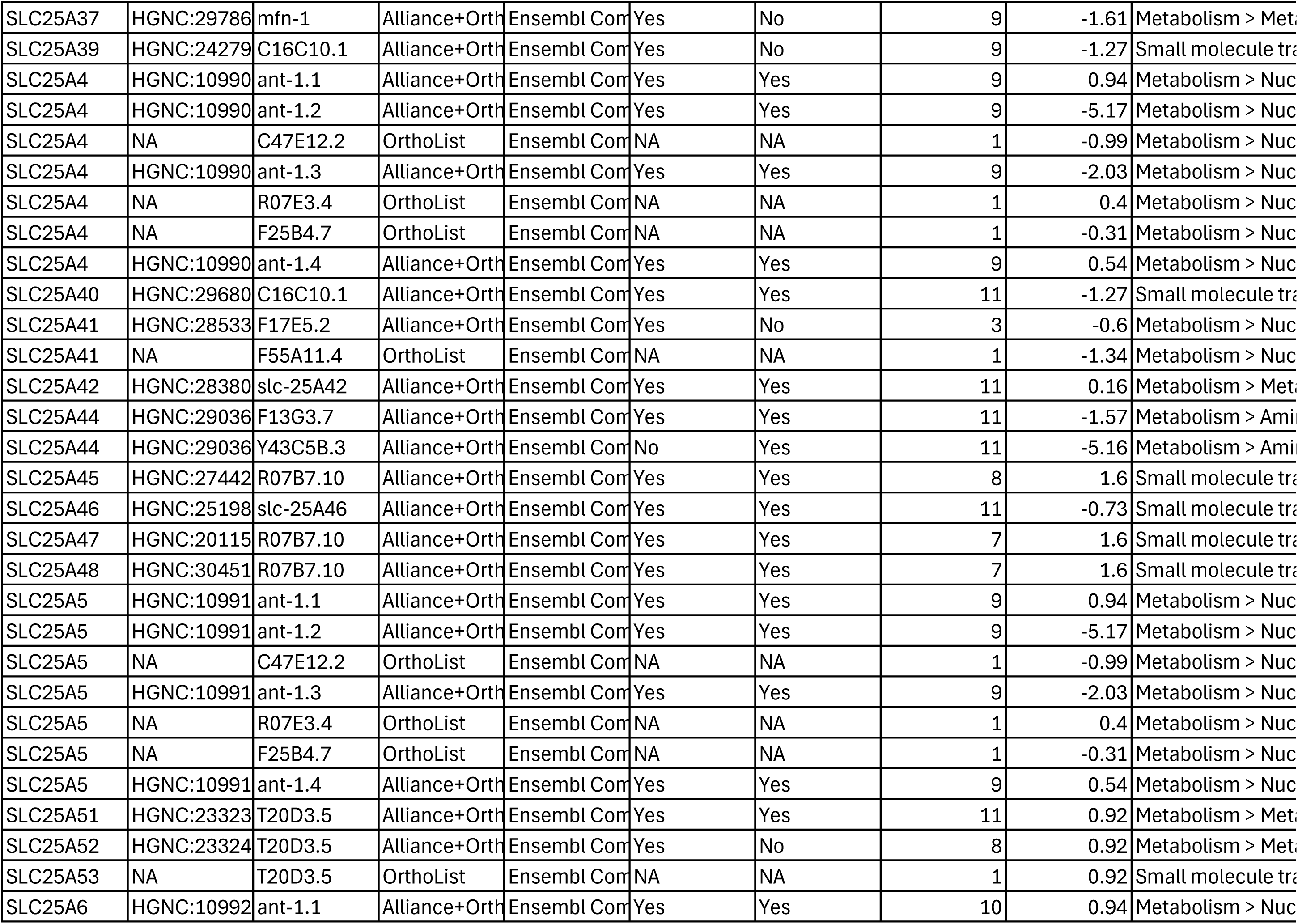
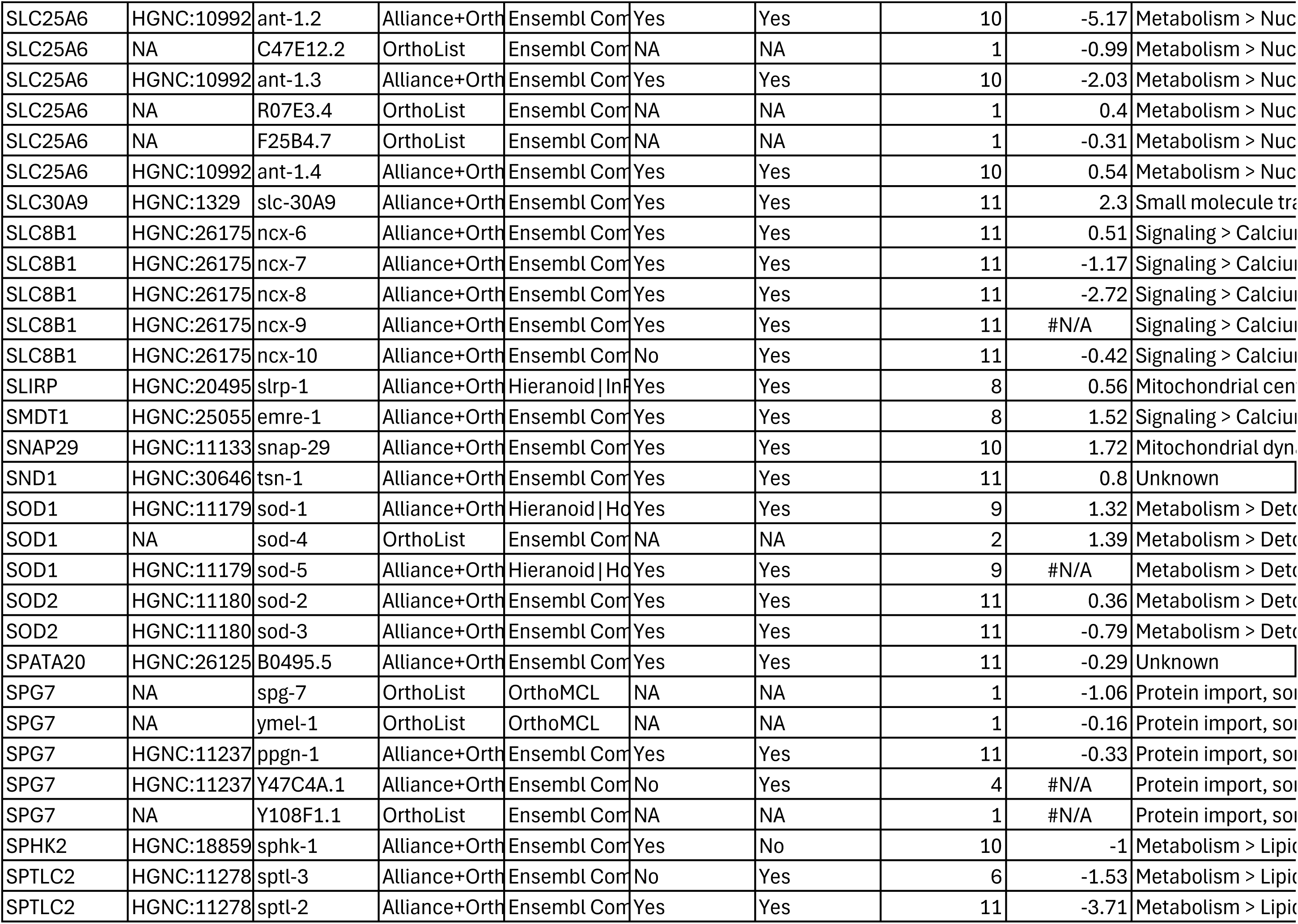
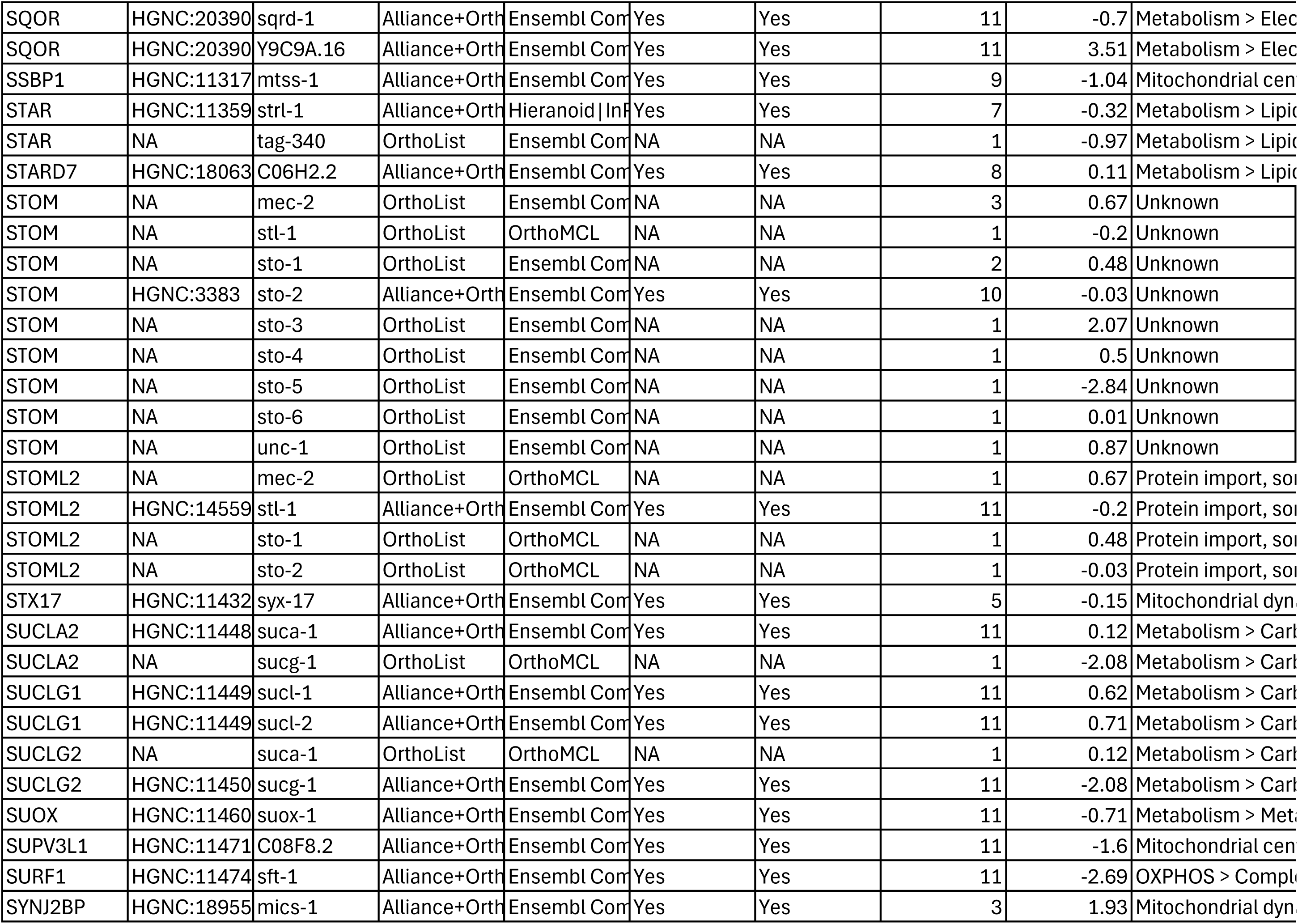
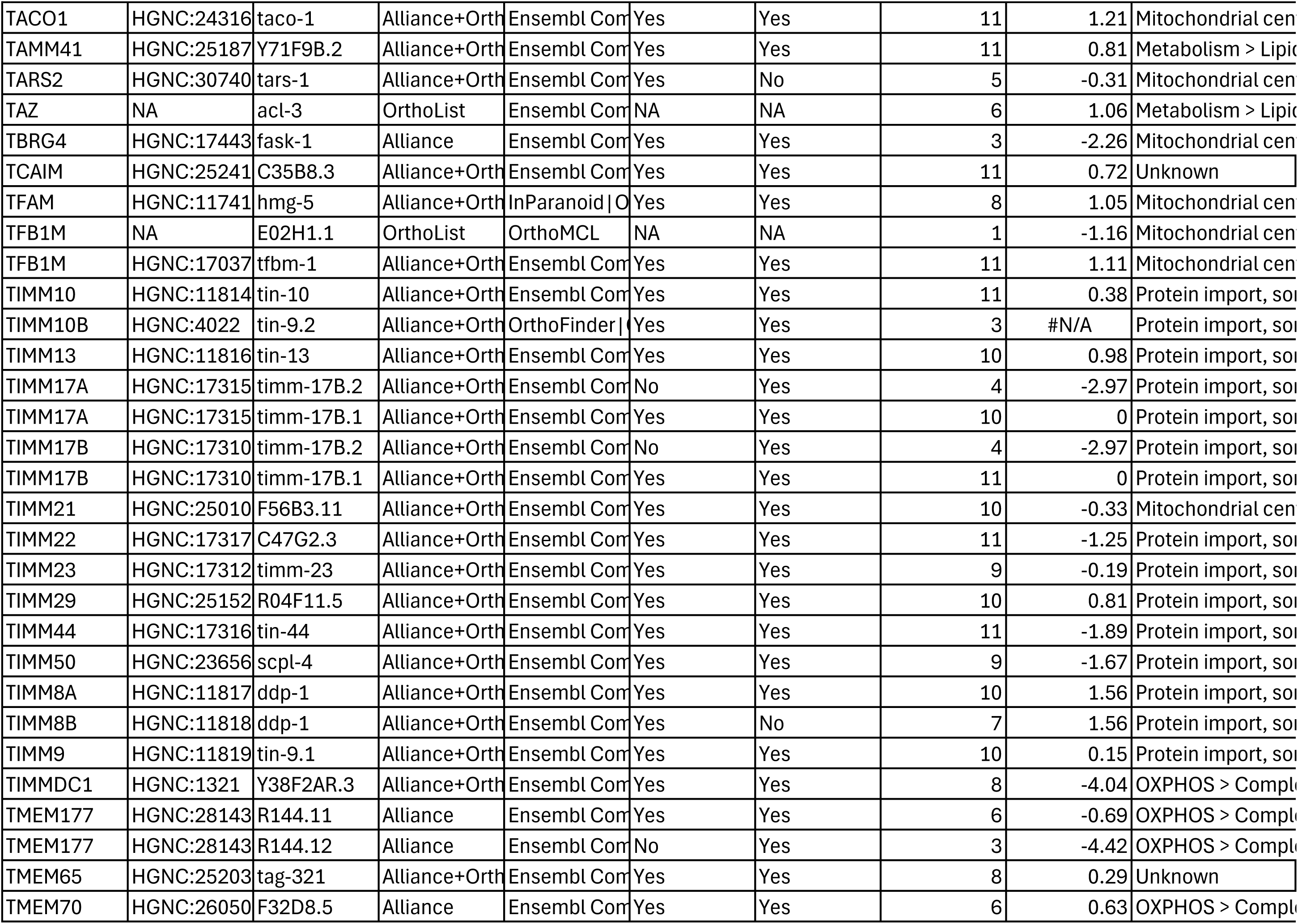
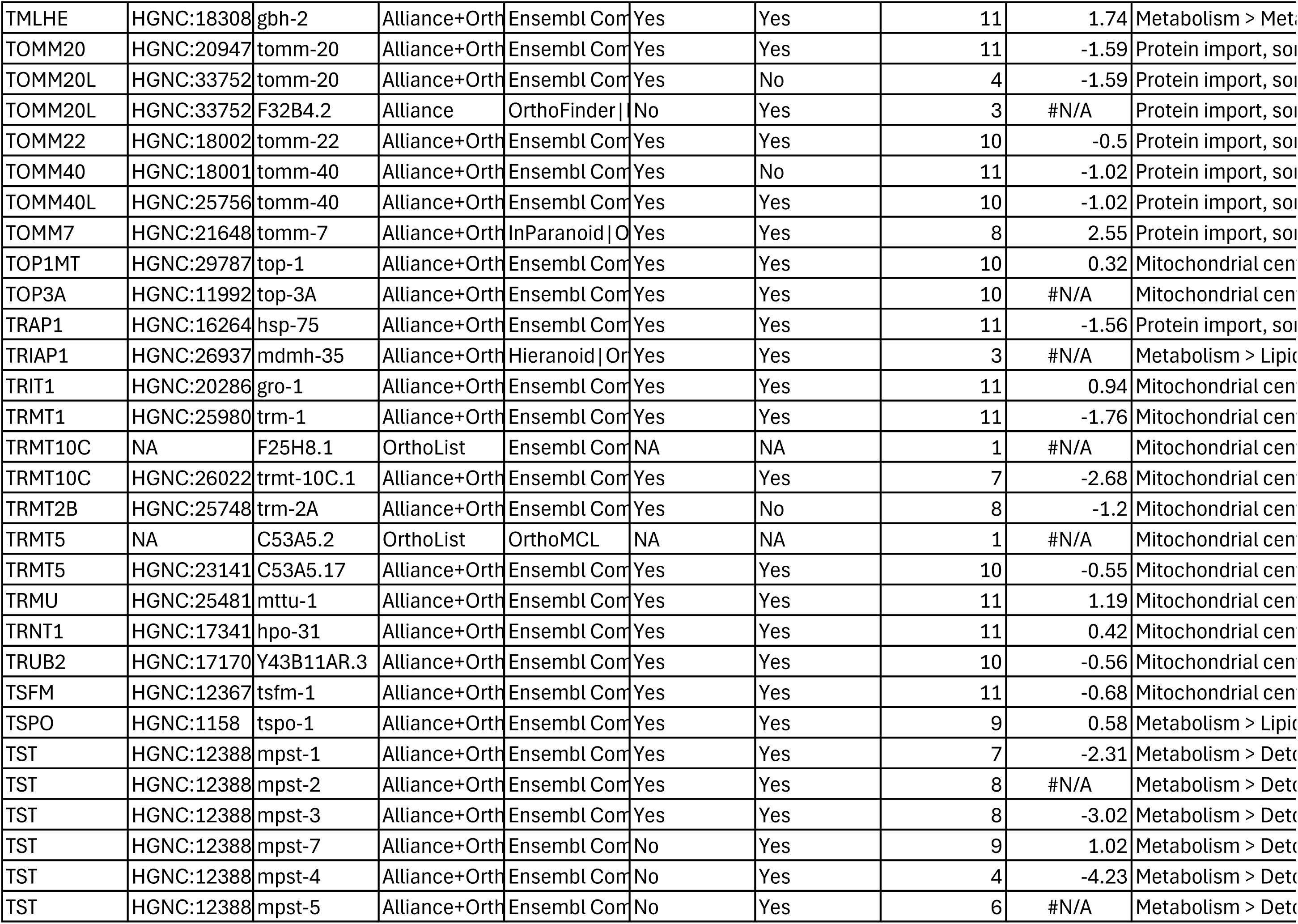
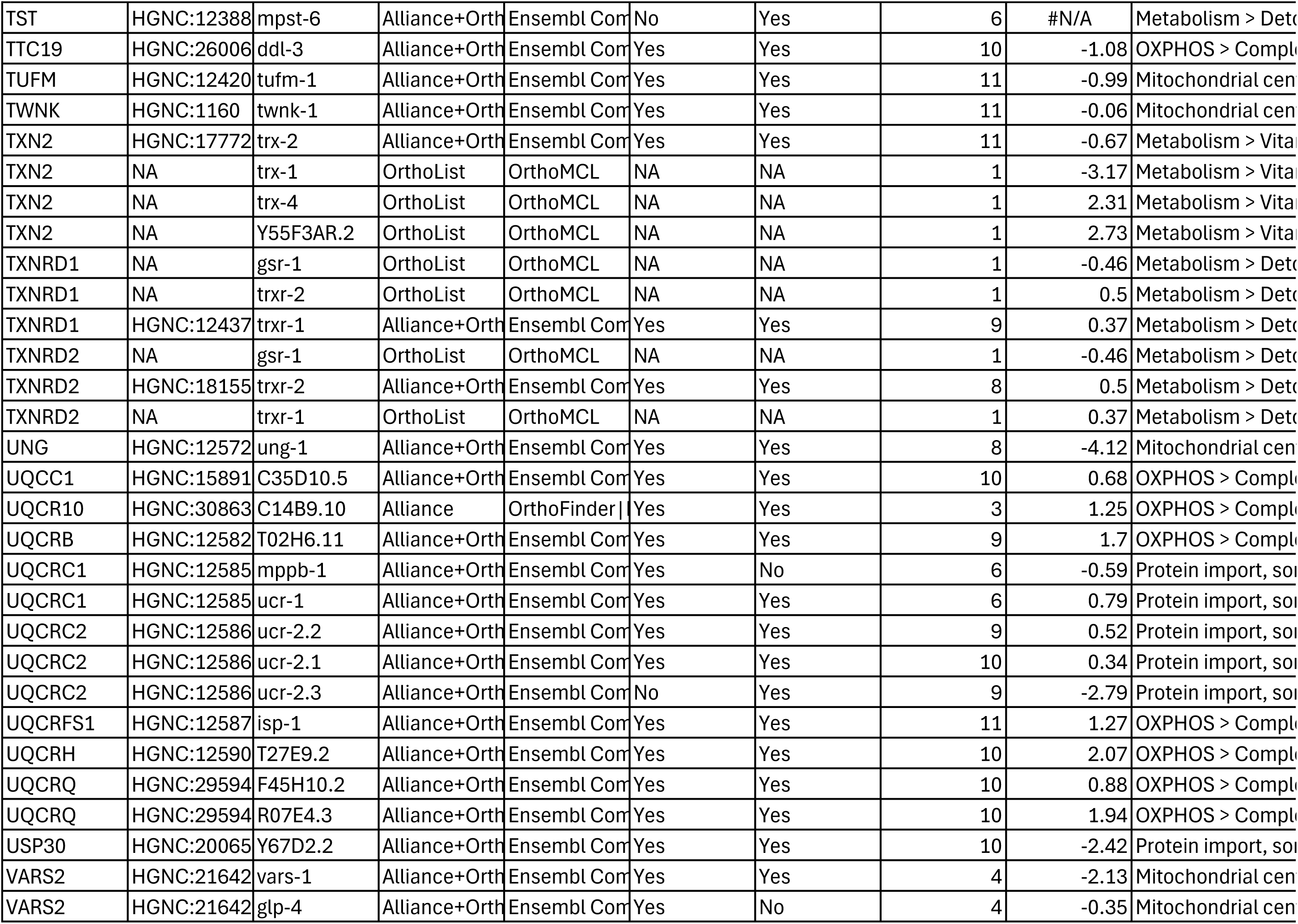
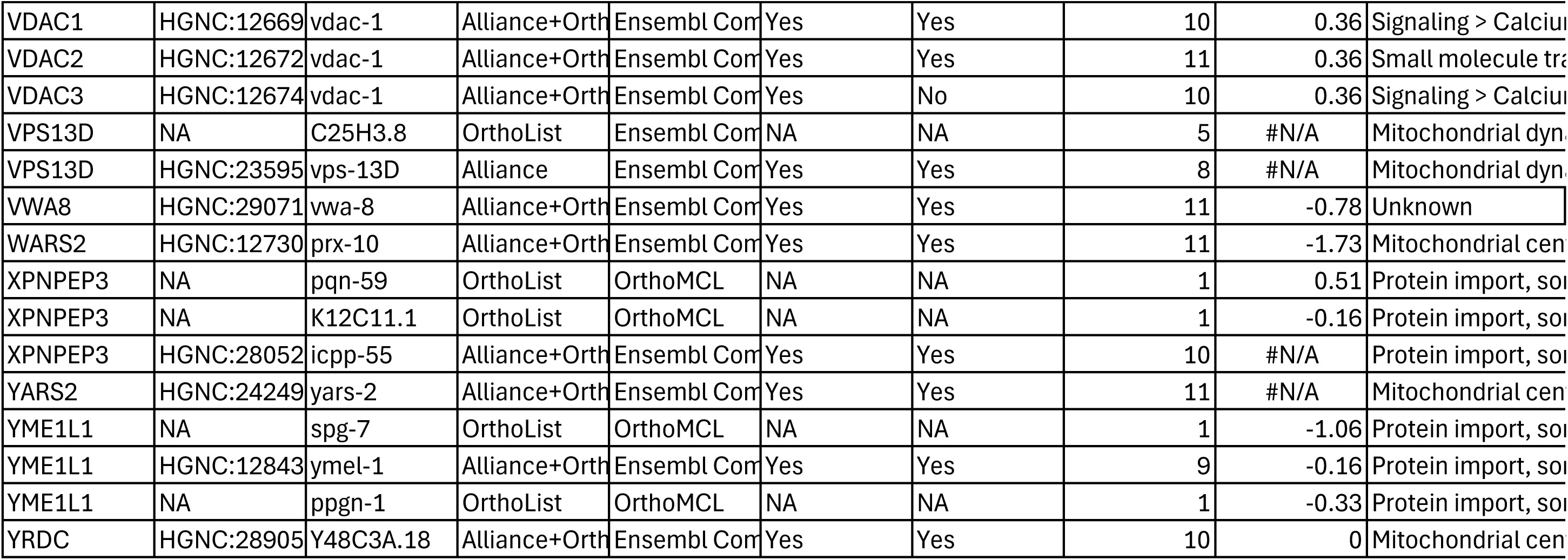
*C.elegans* MitoCarta.

**Table S3.**
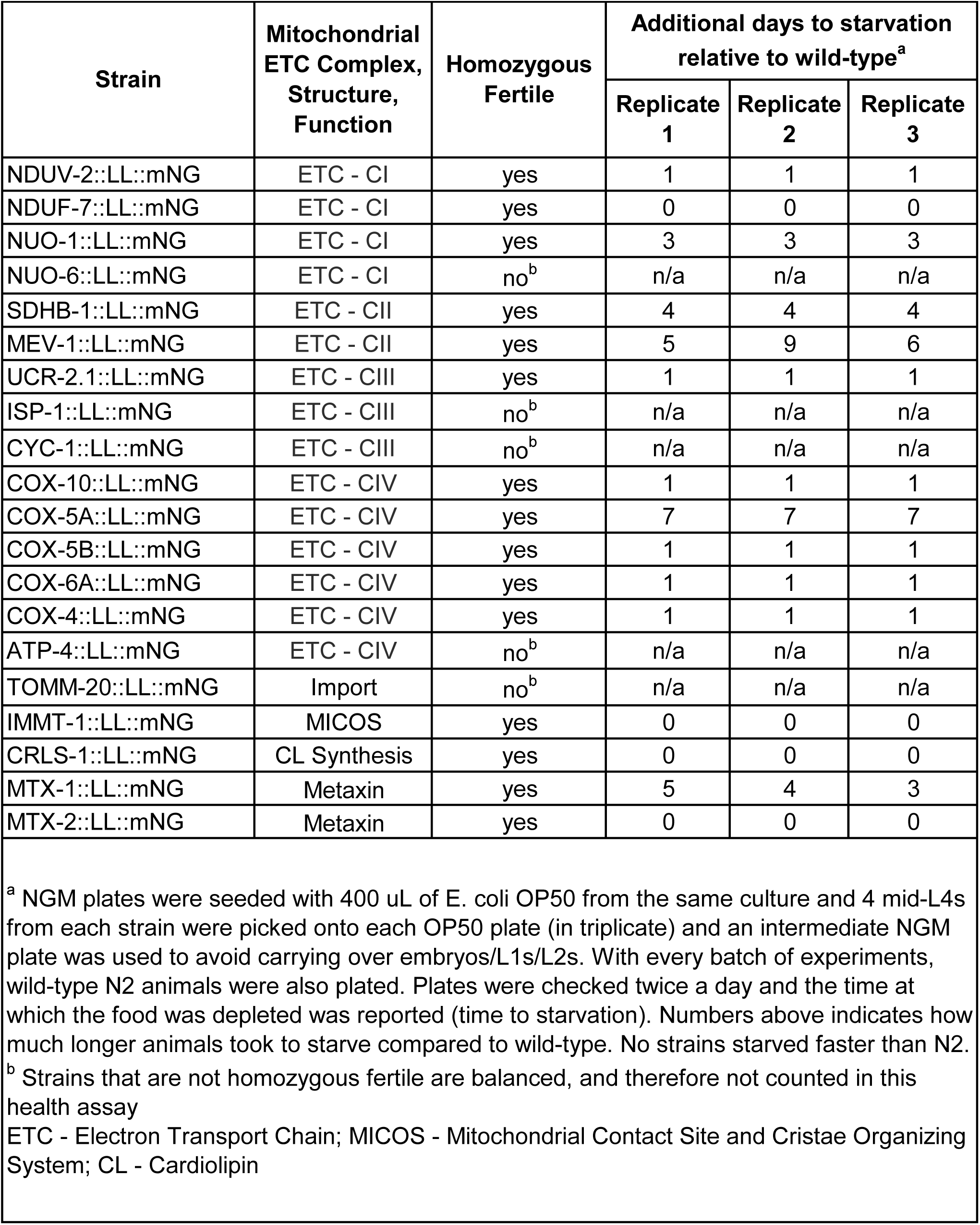
Health of fluorescently tagged mitochondrial strains.

**Table S4.** Mitochondrial Import and Cristae Screen.

| Human Gene | Role | RNAi Condition | Sequence ID | Decreased AC NUO-1 enrichment<sup>a</sup> | n | p-value<sup>b</sup> |
| --- | --- | --- | --- | --- | --- | --- |
| n/a | n/a | n/a | empty vector | 1.2% | 165 | n/a |
| TOMM40 | Import | *tomm-40* | C18E9.6 | 73.3% | 30 | <0.0001 |
| TOMM70 | Import | *tomm-70* | ZK370.8 | 46.7% | 30 | <0.0001 |
| TOMM20 | Import | *tomm-20* | F23H12.2 | 73.3% | 15 | <0.0001 |
| TIMM23 | Import | *timm-23* | F15D3.7 | 80.0% | 15 | <0.0001 |
| APOOL | MICOS | *moma-1* | K02F3.10 | 6.7% | 15 | 0.2309 |
| MICOS13 | MICOS | *W04C9.2* | W04C9.2 | 20.0% | 15 | 0.0043 |
| CHCHD3 | MICOS | *chch-3* | M176.3 | 0.0% | 15 | >0.9999 |
| MICOS10 | MICOS | *F54A3.5* | F54A3.5 | 6.7% | 15 | 0.2309 |
| DNAJC11 | MICOS | *dnj-9* | F11G11.7 | 40.0% | 30 | <0.0001 |
| IMMT | MICOS | *F36D4.6* | F36D4.6 | 20.0% | 15 | 0.0043 |
| IMMT | MICOS | *immt-2* | W06H3.1 | 26.7% | 15 | 0.0004 |
| SAMM50 | MICOS | *gop-3* | C34E10.1 | 0.0% | 15 | >0.9999 |
| IMMT | MICOS | *immt-1* | T14G11.3 | 33.3% | 30 | <0.0001 |
| SYNJ2BP | MICOS | *mics-1* | T21C9.1 | 33.3% | 30 | <0.0001 |
| CRLS1 | CL Synthesis | *crls-1* | F23H11.9 | 33.3% | 30 | <0.0001 |

<sup>a</sup> Decreased enrichment scored by eye on fluorescent compound microscope. If AC was not distinguishable from other uterine cells by NUO-1::mNG fluorescence intensity, then it was scored as having decreased NUO-1 enrichment
<sup>b</sup> One-way ANOVA with Tukey's post hoc test for multiple comparisons
MICOS - Mitochondrial Contact Site and Cristae Organizing System; CL - Cardiolipin

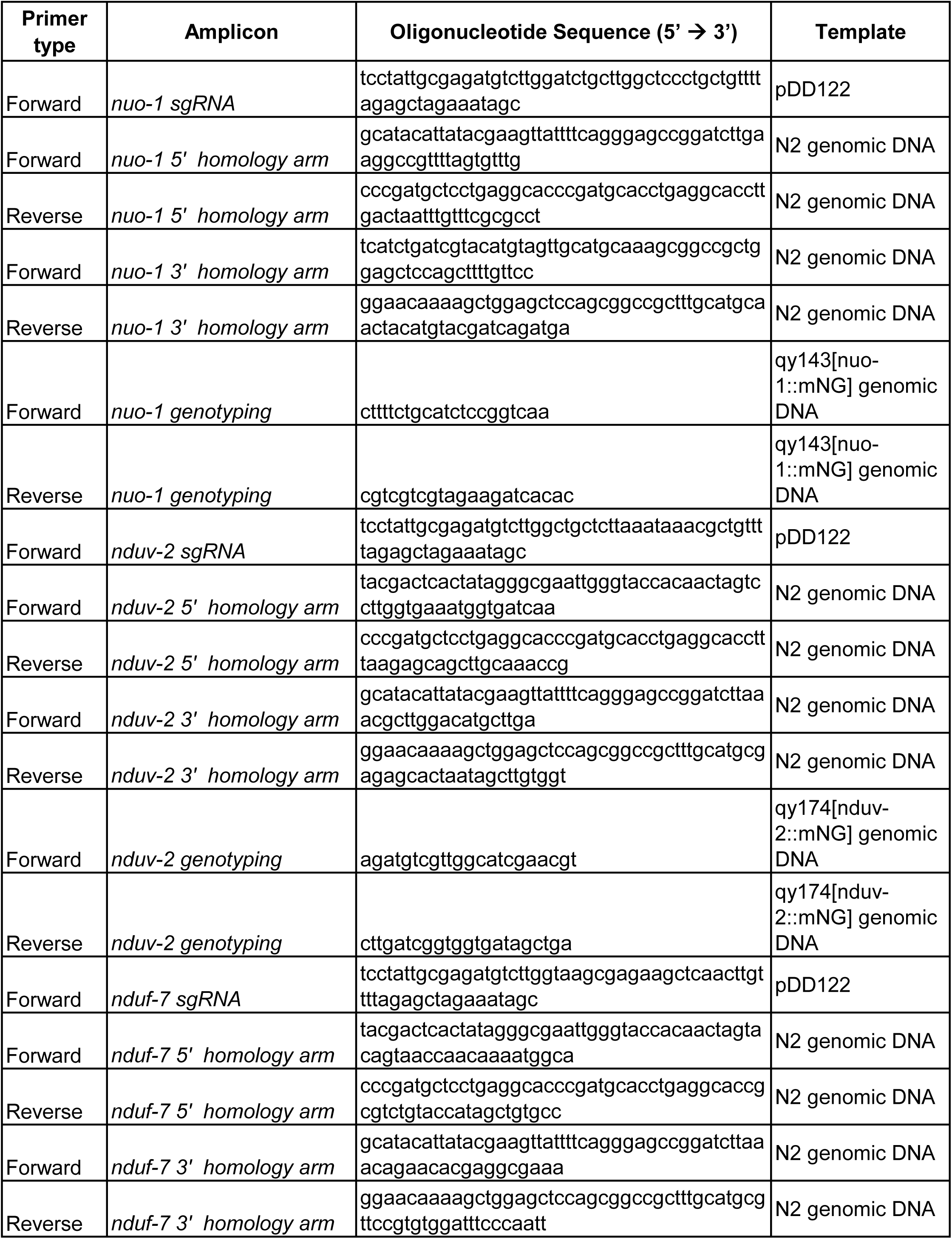
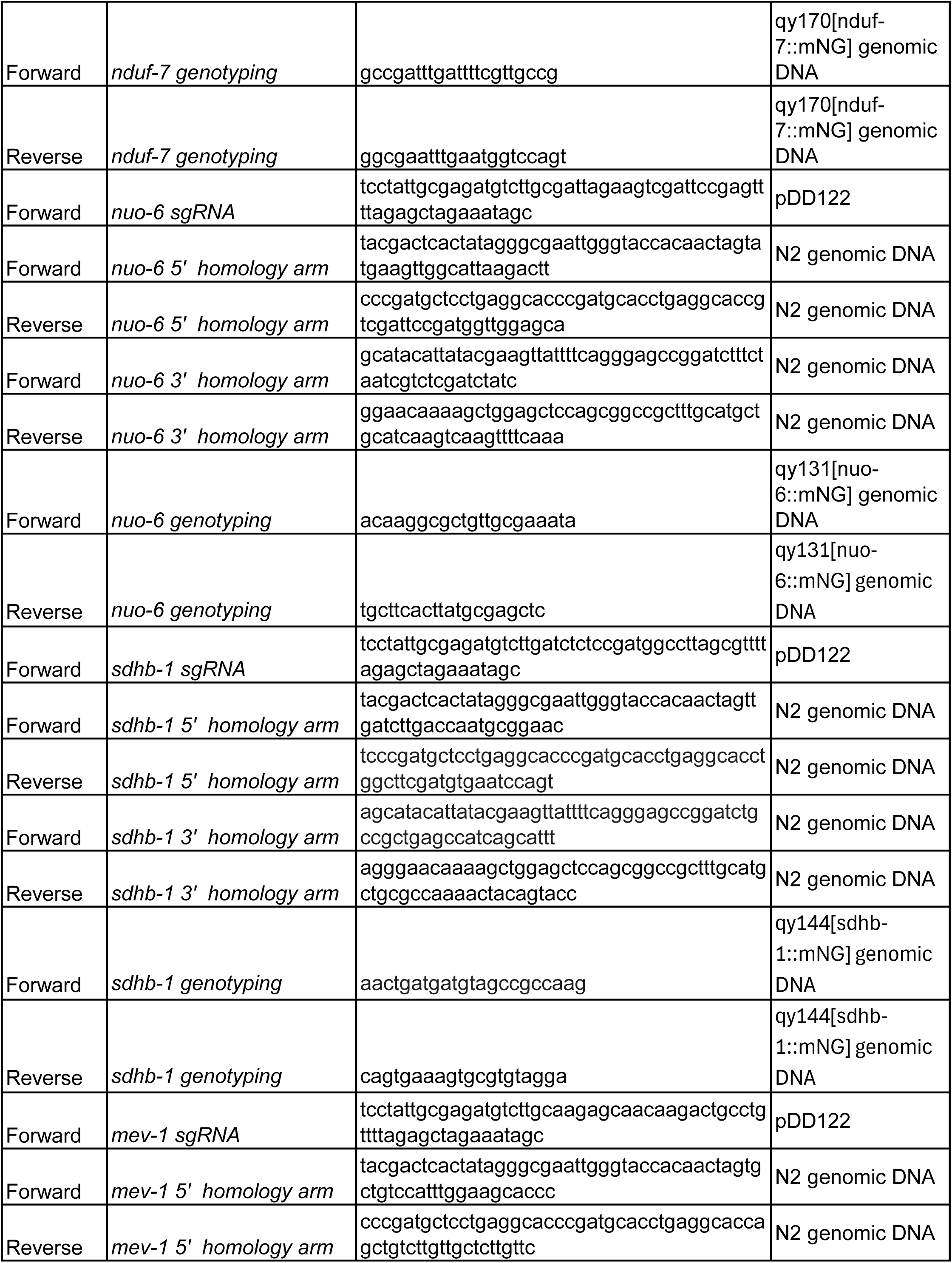
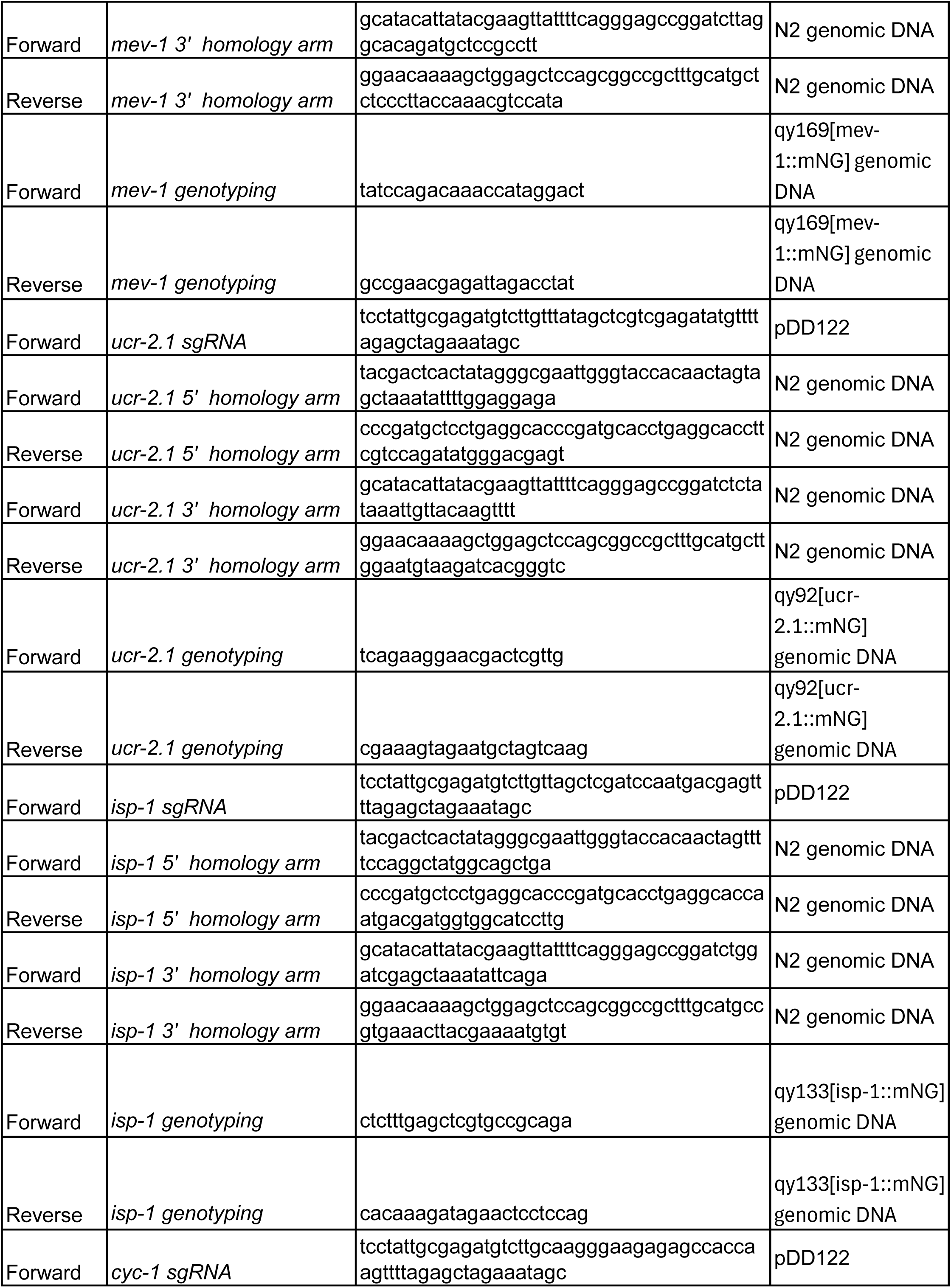
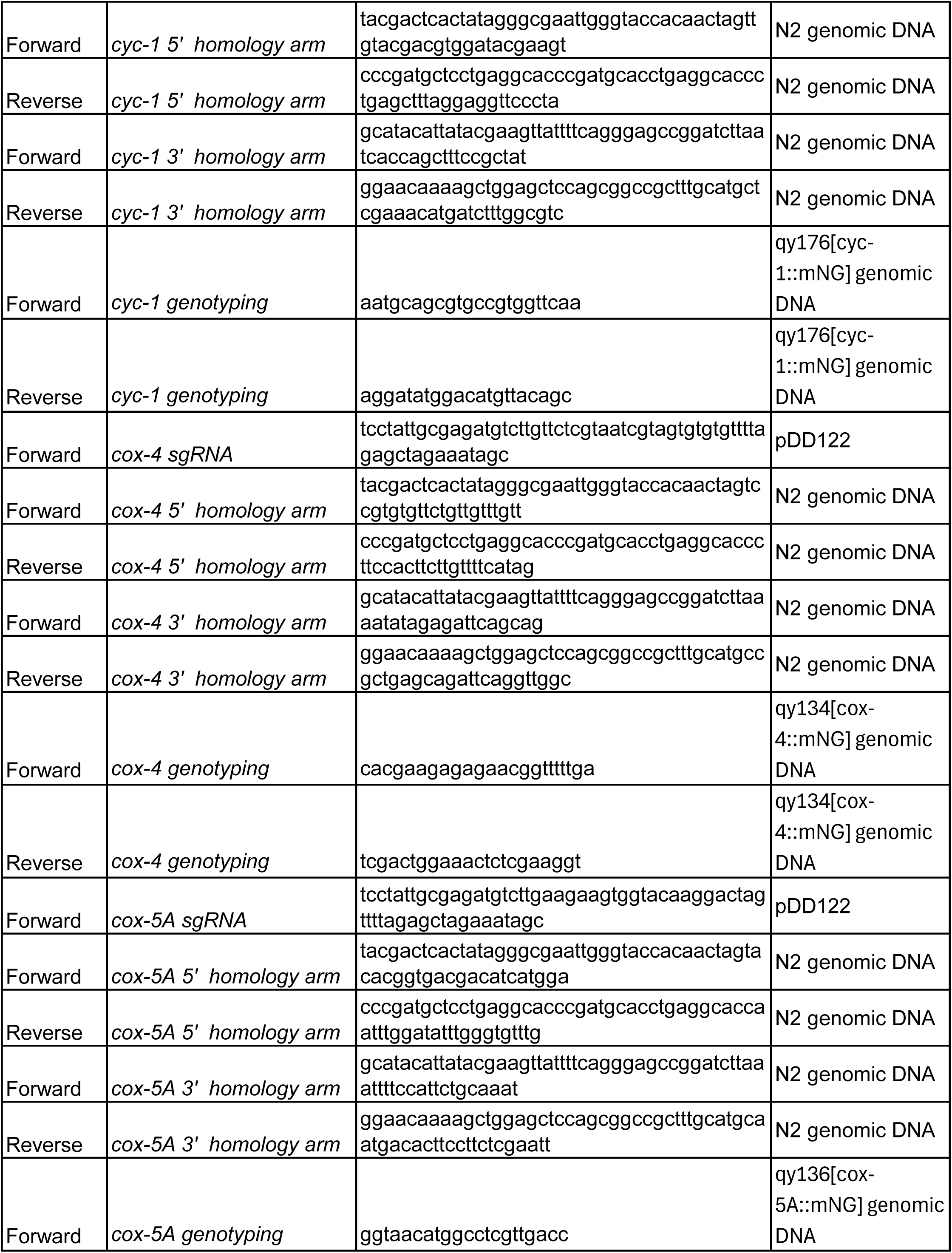
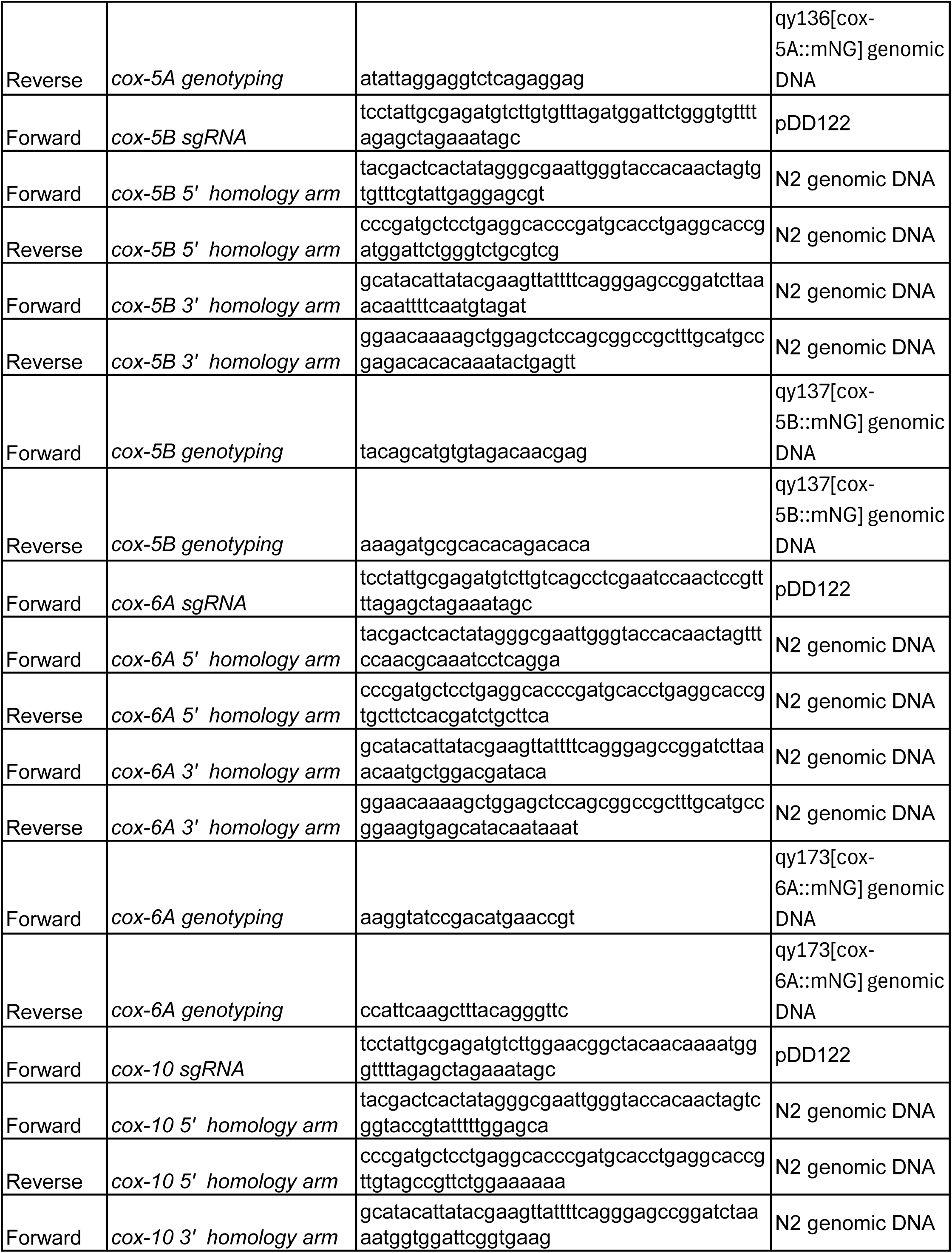
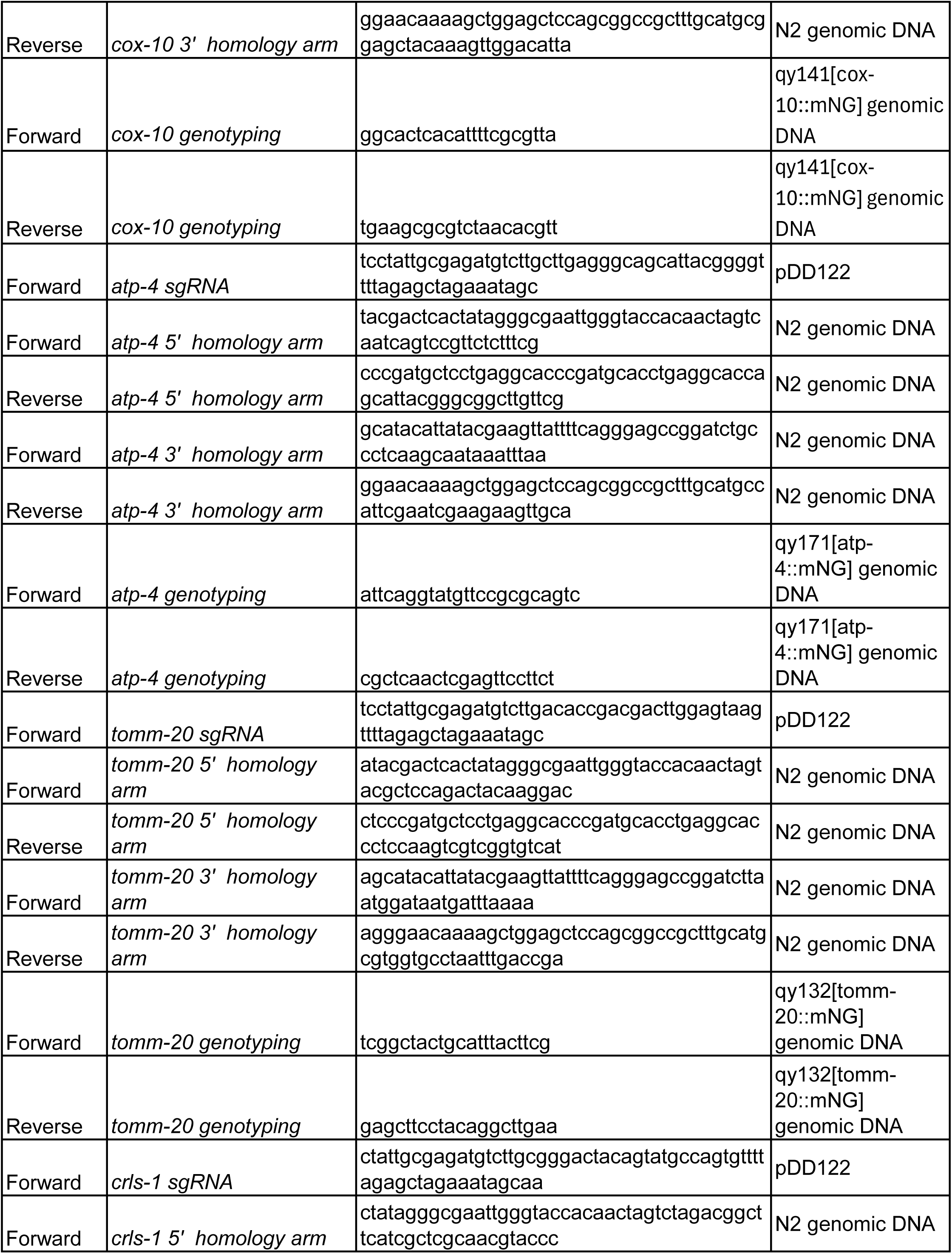
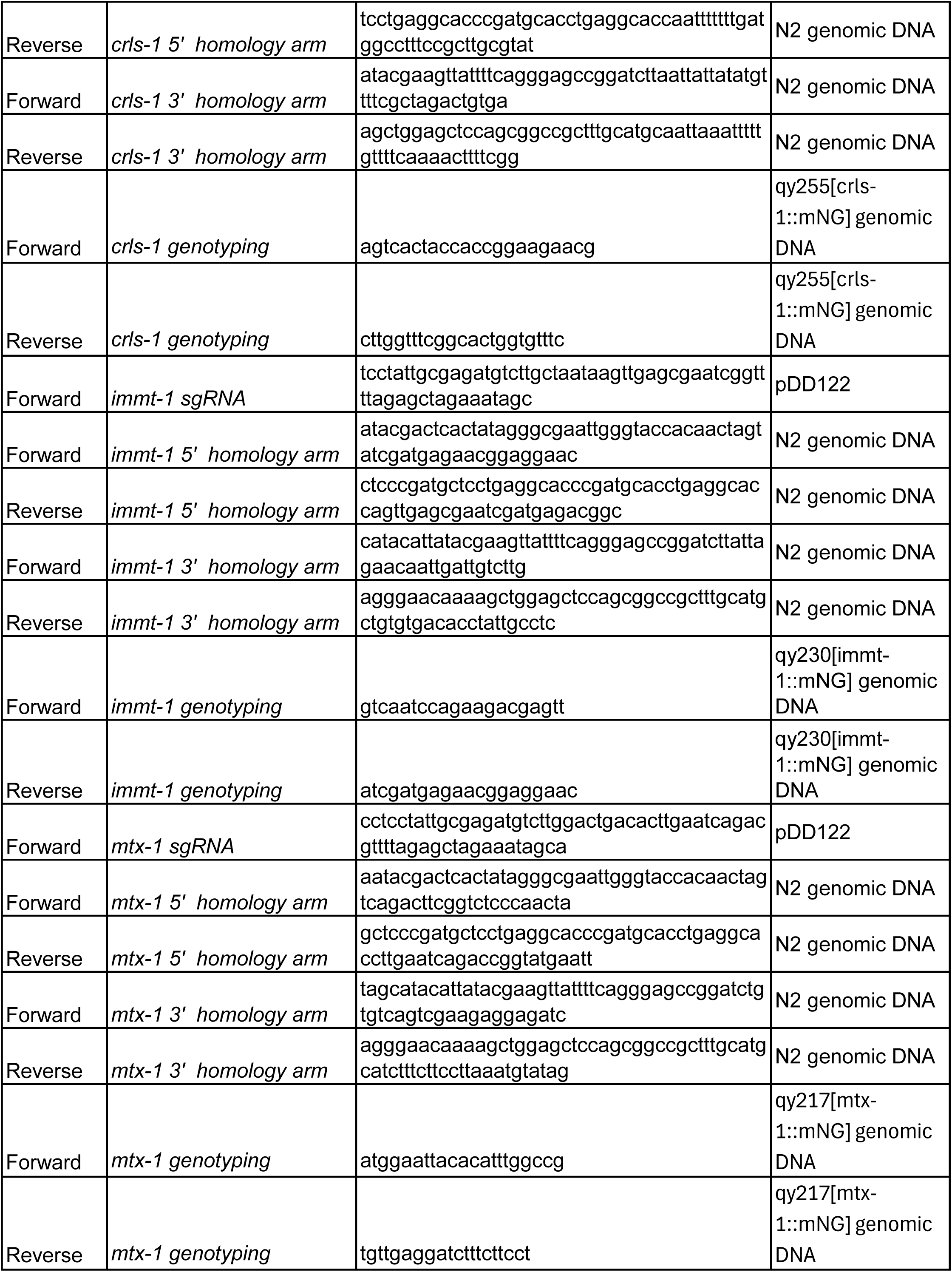
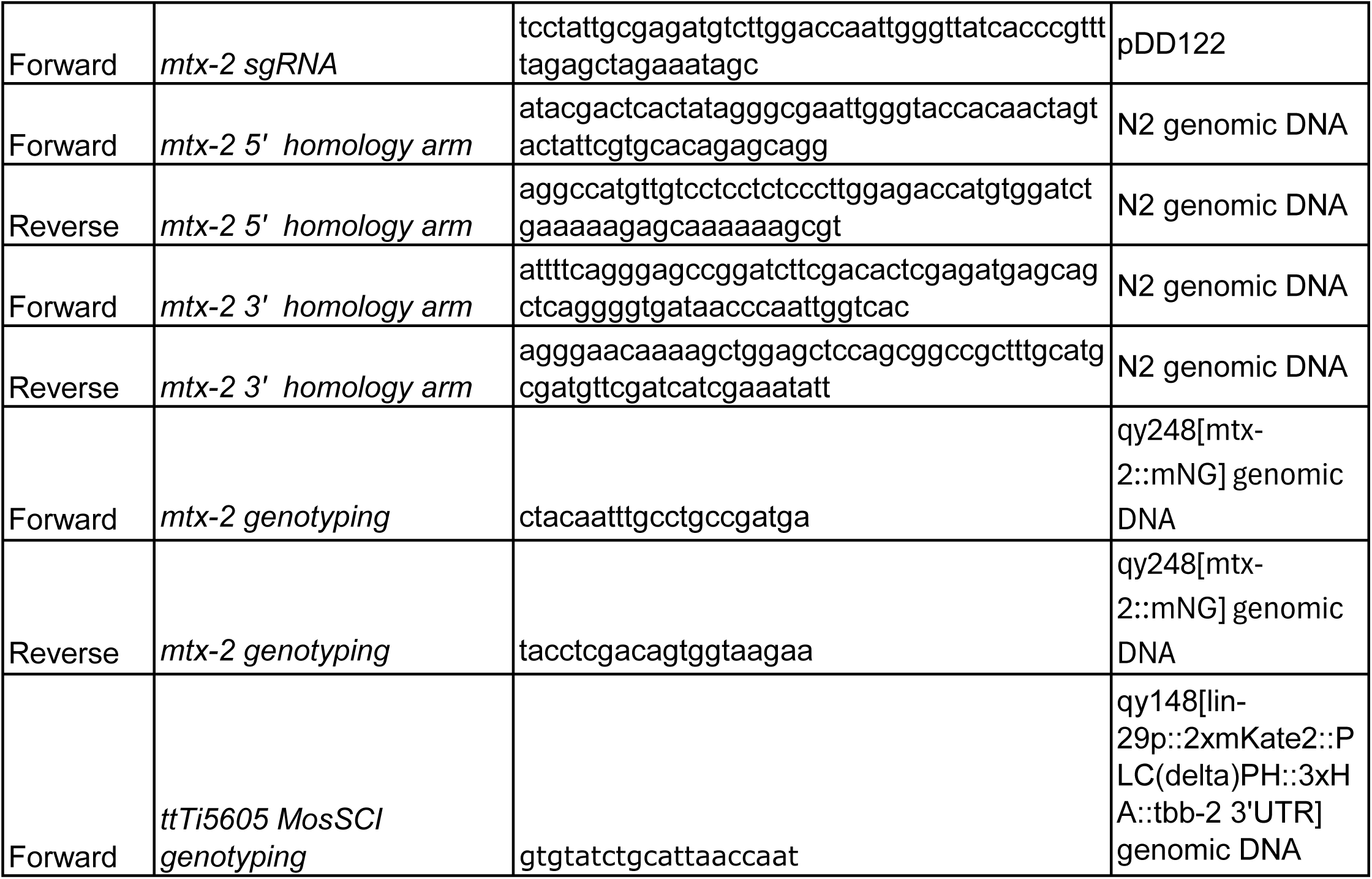
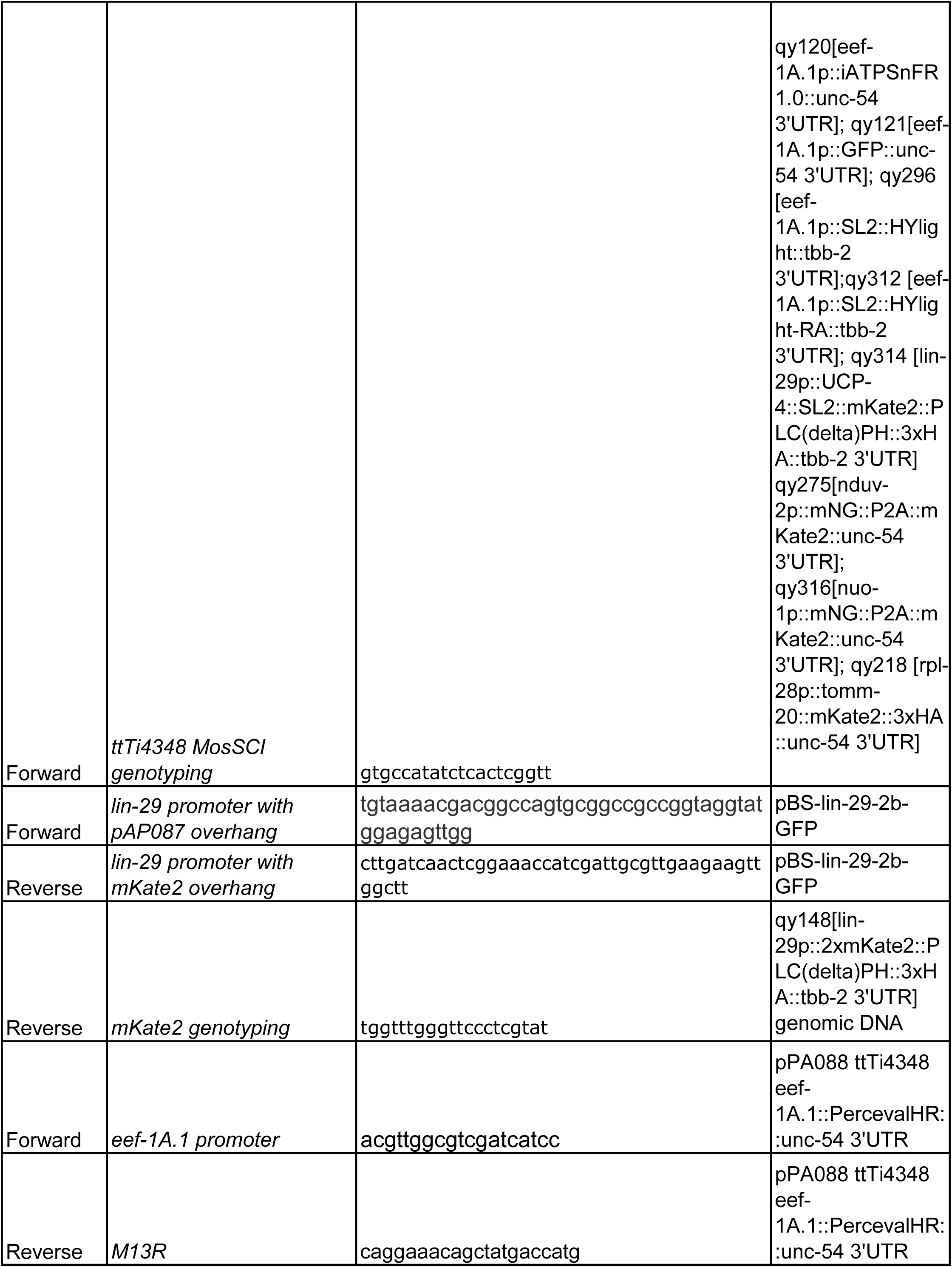
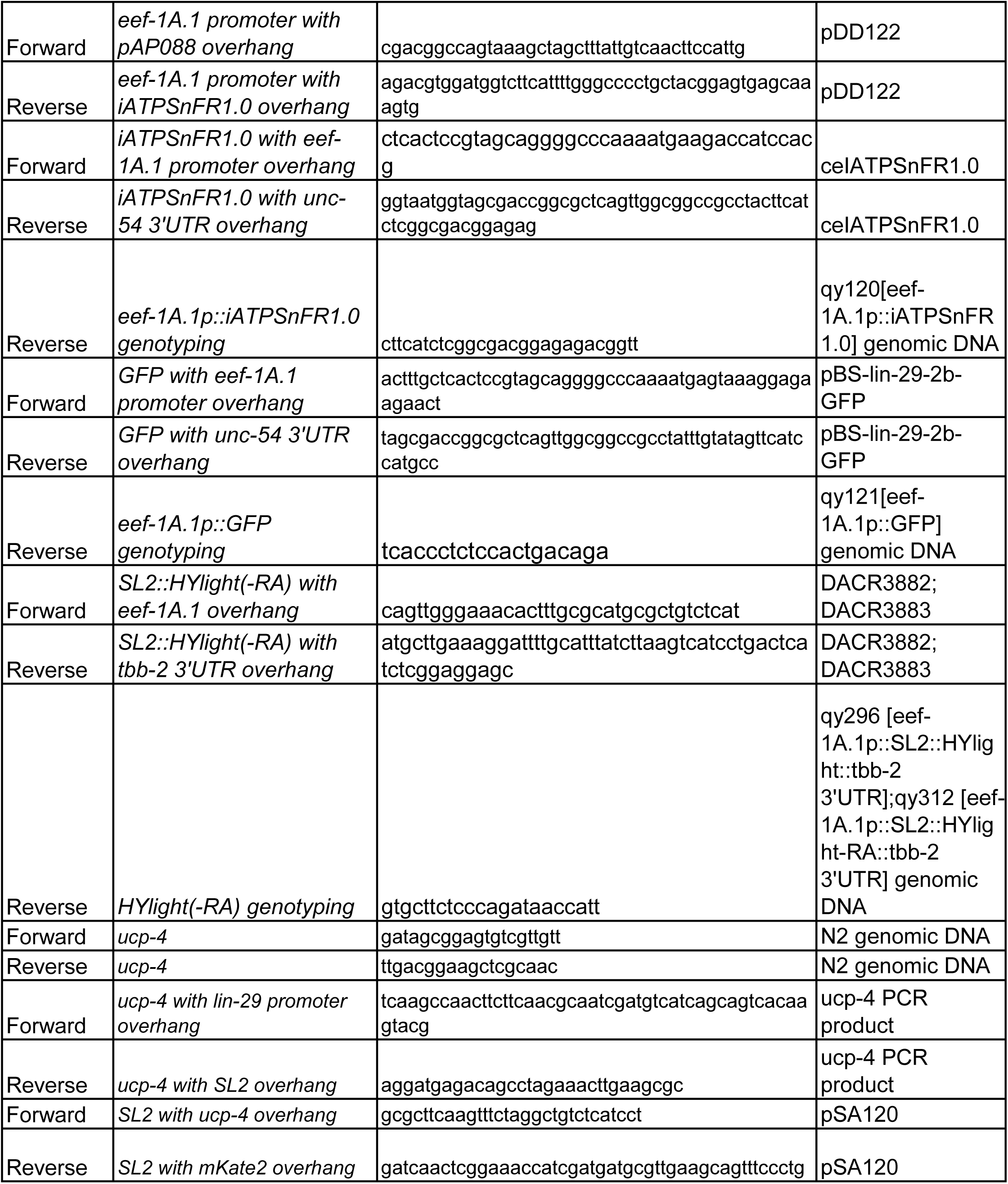
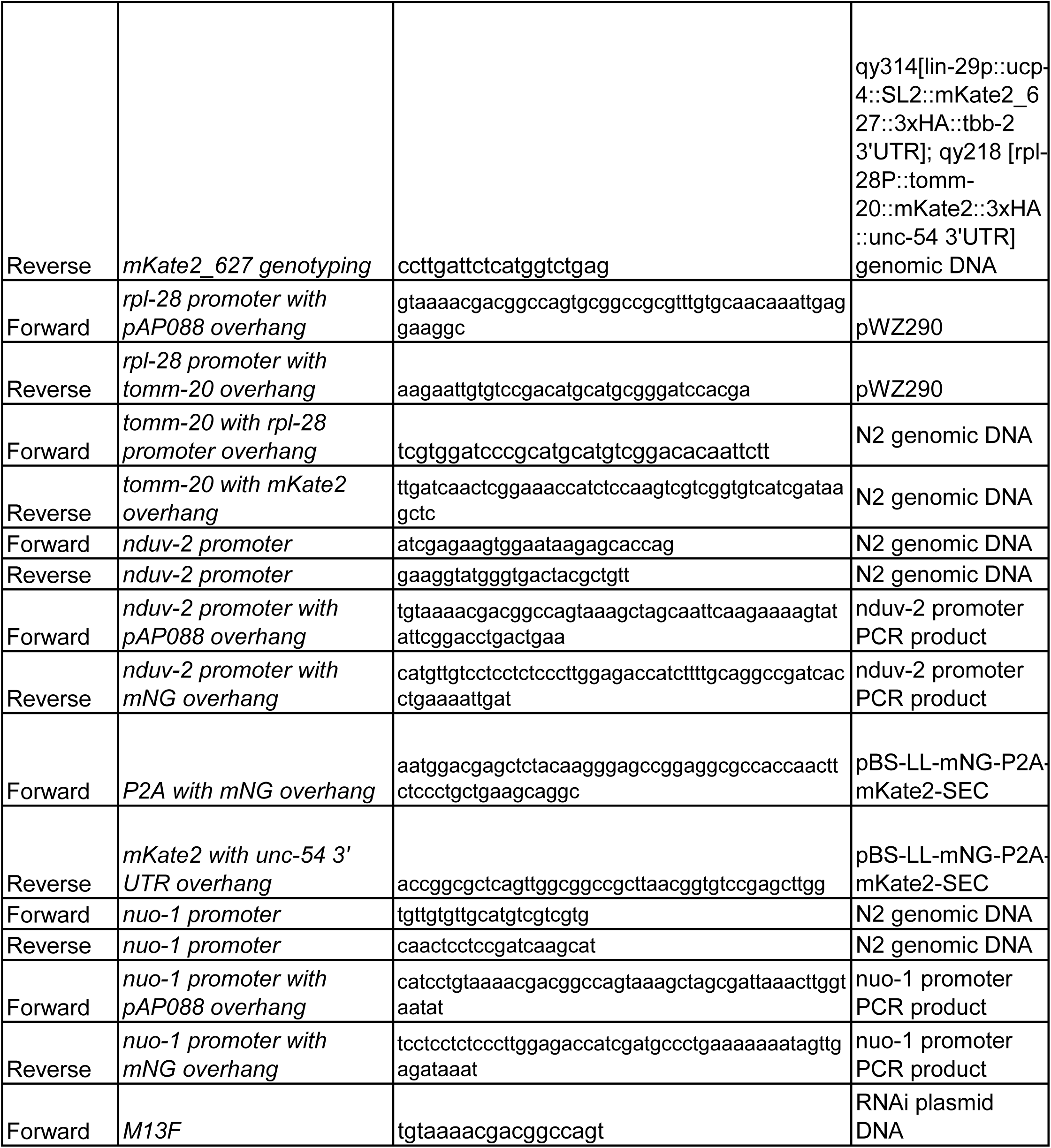

**Table S6.** RNAi knockdown efficiency.

| Gene targeted | Strain genotype<sup>a</sup> | Treatment | Percent protein knockdown (%) | n<sup>b</sup> |
| --- | --- | --- | --- | --- |
| *ucr-2.1* | *qy92[ucr-2.1::mNG] X* | *ucr-2.1 RNAi* | 63.8 | ≥ 27 |
| *nuo-1* | *qy143[nuo-1::mNG] II* | *nuo-1 RNAi* | 93.7 | ≥ 13 |
| *nduv-2* | *qy174[nduv-2::mNG] V* | *nduv-2 RNAi* | 66.7 | ≥ 11 |
| *immt-1* | *qy230[immt-1::mNG] X* | *immt-1 RNAi* | 71.1 | 13 |
| *mtx-1* | *qy217[mtx-1::mNG] I* | *mtx-1 RNAi* | 83.5 | ≥ 14 |
| *mtx-2* | *qy248[mtx-2::mNG] III* | *mtx-2 RNAi* | 61.3 | 8 |

<sup>a</sup> All RNAi knockdown efficiency was performed using strains endogenously edited with
<sup>b</sup> Number of animals imaged in each condition

## Notes

### Competing Interest Statement

The authors have declared no competing interest.

